# AssemblyTron: Flexible automation of DNA assembly with Opentrons OT-2 lab robots

**DOI:** 10.1101/2022.09.29.510219

**Authors:** John A. Bryant, Mason Kellinger, Cameron Longmire, Ryan Miller, R. Clay Wright

## Abstract

As one of the newest fields of engineering, synthetic biology relies upon a trial-and-error Design-Build-Test-Learn approach to simultaneously learn how function is encoded in biology and attempt to engineer it. Many software and hardware platforms have been developed to automate, optimize, and algorithmically perform each step of the Design-Build-Test-Learn cycle. However, there are many fewer options for automating the Build step. Build typically involves DNA assembly, which remains manual, low throughput, and unreliable in most cases, limiting our ability to advance the science and engineering of biology. Here, we present AssemblyTron: an open-source python package to integrate j5 DNA assembly design software outputs with build implementation in Opentrons liquid handling robotics with minimal human intervention. We demonstrate the versatility of AssemblyTron through several scarless, multipart DNA assemblies beginning from fragment amplification. We show that AssemblyTron can perform PCRs across a range of fragment lengths and annealing temperatures by using an optimal annealing temperature gradient calculation algorithm. We then demonstrate that AssemblyTron can perform Golden Gate and homology-dependent *in vivo* assemblies with comparable fidelity to manual assemblies by simultaneously building four four-fragment assemblies of chromoprotein reporter expression plasmids. Finally, we used AssemblyTron to perform site-directed mutagenesis reactions via homology-dependent *in vivo* assembly also achieving comparable fidelity to manual assemblies as assessed by sequencing. AssemblyTron can reduce the time, training, costs, and wastes associated with synthetic biology, which along with open-source and affordable automation, will further foster the accessibility of synthetic biology and accelerate biological research and engineering.

## 1. Intro

As one of the newest fields of engineering, biological engineering, or synthetic biology, is still largely performed by iterative, trial-and-error Design-Build-Test-Learn (DBTL) cycles, both because our knowledge of biology is incomplete and our ability to build biological devices is error-prone. Molecular biology developments in the past forty years have improved our ability to treat genes, promoters, and other DNA elements as modular parts to engineer increasingly complex devices with biology. However assembly of the DNA elements encoding these devices largely remains error-prone and requires significant tacit, knowledge, and iterative trials to master (1). The speed at which we can engineer biology and learn its design principles relies on our ability to perform DBTL cycles rapidly and error-free. Automation of the build step can accelerate each DBTL cycle, eliminate errors, and allow scientists and engineers to focus their energy on the creative design and learn steps addressing the biological questions they set out to answer, instead of optimizing the physical assembly process of the build step. Notably, a 2017 analysis of biological design automation tools found that there were fewer automation tools available for the Build step than Design, Test, or Learn (2).

To build biological devices researchers assemble multigene DNA constructs using an ever-growing variety of potential DNA assembly methods, including short homology recombination methods such as SLIC (3), Gibson (4), CPEC (5), AQUA (6), and IVA (7) assembly methods, restriction-ligation methods such as Golden Gate (8), and *de novo* synthesis (9,10). Each of these methods has advantages and disadvantages, and a combination of short homology recombination, restriction-ligation, and *de novo* synthesis methods are routinely used in most synthetic biology labs. Importantly, flexible combinatorial fragment assembly methods allow reassembly of different parts (*e*.*g*. various promoters and terminators with a coding sequence) for creating combinatorial construct libraries with high time- and cost-efficiency increasing DBTL cycle throughput. Thus, even as the cost of de novo DNA fragment synthesis continues to decrease, the need will remain for robust, automated assembly capabilities in order to accommodate higher throughput experimentation.

Build automation is key to improving the speed and accuracy of DNA assembly and accelerating the general DBTL cycle, as human-errors are frequent in manual assembly processes. The Edinburgh Genome Foundry has automation workflows that increase throughput 20-fold, while iBioFab at the University of Illinois at Urbana-Champaign has reduced the price of construct assembly by 97.7% (11,12). Though these impressive figures underpin the value of biofoundries, high experimental costs still prevent most of the academic community from taking advantage of these facilities or incorporating automation into their labs. Additionally, standard procedures in established biofoundries including software, hardware, and methods are not interoperable, so their services are often useful to only a small pool of researchers (13). Though a flexible, open-source build automation platform would be a good alternative, few exist (14,15), particularly those which make use of liquid handling robotics. Many of the build platforms that do exist are fairly inaccessible, difficult to find, and not interoperable (2,16).

Recently, low-cost open-source liquid handling robotics systems have created an opportunity to increase the throughput and decrease the human-error associated with the build step of the DBTL cycle (17). The Opentrons OT-2 is an open-source liquid handling robot with thermocycler capabilities. Its advanced API library for pipette manipulation and protocol development is highly flexible for a wide array of molecular biology applications. However, there is not yet an open-source python software package or protocols to enable the OT-2 to execute the popular short homology and Golden Gate DNA assembly protocols, such as those generated by automated DNA assembly design algorithms j5 and Cello (18,19). To our knowledge the only build automation software for the OT-2 is DNA-BOT (20) which executes the Biopart Assembly Standard for Idempotent Cloning (BASIC) method (21).

Numerous options exist for researchers to automate the generation of DNA device designs and optimize assembly protocols which implement commonly used, scarless DNA assembly methods, but these often do not integrate with liquid handling robotics. To minimize researcher-to-researcher variation in primer and assembly design and maximize the likelihood of assembly success, several software platforms have been developed to automate the design and optimization of DNA assemblies using vetted algorithms (18,19). The j5 construct design algorithm is a valuable tool for standardizing the design and assembly workflow while minimizing the need for DNA synthesis (18). Interpretation and implementation of the DNA assembly as specified in j5 output remains a bottleneck in DBTL cycle. There is a significant amount of training required for new researchers to master DNA assembly and a high potential for errors during training. This training barrier often prevents undergraduates from being successful in their first synthetic biology experiments. Software for processing output files from DNA assembly design software, such as j5, and generating protocols for this new generation of relatively affordable laboratory robotics, such as the OT-2, would expedite automation efforts in public laboratories, avert errors on the benchtop, and allow researcher to focus on the more critical test and learn steps.

We seek to minimize human error rate in the build step of the DBTL cycle by automating the physical assembly workflow with open-source software that implements j5 assembly designs in the OT-2 liquid handling robotics system with minimal human intervention. We aimed to create a build automation platform that supports existing DNA assembly protocols and conventions. We also focused on leveraging existing DNA design software and liquid handling robotics hardware platforms to expedite synthetic biology across academic research labs in an economically accessible way. Here, we present AssemblyTron: an open-source python package to implement DNA assembly design software output (currently j5 specifically) using the OT-2 liquid handling robot (22). This package is designed to automate the DNA assembly process between the design stage and transformation in *E. coli* with minimal human intervention. AssemblyTron automates enzyme-free homology-dependent assembly methods such as AQUA (6) and IVA (7), which serve as fast and cheap alternatives to Gibson since they depend on native *E. coli* machinery to ligate fragments. It also automates Golden Gate (8) assembly to offer a highly accurate and flexible assembly strategy option for complex designs or libraries. AssemblyTron is an open-source build automation framework that can reduce the time, training, costs, and excesses associated with molecular biology workflows. This allows researchers to spend more time asking questions and building new synthetic biology tools instead of troubleshooting their assembly workflow.

## 2. Methods

### 2.1 *E. coli* strains and reagents

Media and other reagents were prepared according to Sambrook and Russel 2012, unless otherwise specified (23). *E. coli* TOP10 chemically competent cells were prepared by the Hanahan method (24) (efficiency of 8.47×10^8 CFU/ug supercoiled pUC19). Lysogeny Broth, Miller (Fisher BioReagents) medium with kanamycin or ampicillin at 50 or 100 ug/ml respectively was used for growing *E. Coli* with the addition of 0.7% (w/v) bactoagar for plating. NEB Monarch Plasmid MiniPrep kits were used for isolating plasmid DNA from *E. coli*, and the Zymogen DNA clean and concentrate kits were used to purify PCR products prior to assembly or transformation. Agarose gels (1% (w/v) in 1X TAE) were stained with biotium GelRed® Nucleic Acid Gel Stain and bands were imaged using an iBright imaging system (Thermo Fisher). The 1 kb plus DNA ladder (NEB) was used for fragment length comparison via agarose gel electrophoresis.

### 2.2 Primers and plasmids

Primers were designed using j5 (18) and were purchased from Integrated DNA Technologies (IDT) (Table 1 and supplementary file 11). J5 is free for use in academic labs (j5.jbei.org). Plasmids used included pGP8A-ARF19 (gifted from the Nemhauser lab), pGP8A-ARF5-pdar, pGP8A-ARF7-pdar, pIDMv5K-J23100-tsPurple-B1006, pIDMv5K-J23100-YukonOFP-B1006, pIDMv5K-J23100-aeBlue-B1006, and pIDMv5K-J23100-fuGFP-B1006 (gifted from Sebastian Cocioba) (Table 2 and supplementary file 12).

### 2.3 DNA Fragment construction and plasmid assembly

PCRs were performed in 25 μL volumes using Phusion® High-Fidelity DNA polymerase (NEB) or Q5® High-Fidelity DNA Polymerase (NEB) with 0.1 uM primers and 0.5 ng linearized plasmid template DNA. Template plasmids were linearized with restriction enzymes cutting outside of the desired amplicons to improve amplification. Amplification was performed according to the following protocol: 30 sec at 98 °C, 34 or 36 cycles of 10 sec at 98 °C, 30 sec at annealing temperatures specified by AssemblyTron/j5, extension times as specified by AssemblyTron at 72 °C, and a final 5 min extension at 72 °C. For plasmid template digestion, 19 μL water, 5 μL rCutSmart Buffer (NEB), and 1 μL DpnI (NEB) was added and incubated for 30 min at 37 °C prior to deactivation at 65 °C for 20 min.

Final assemblies for Golden Gate (Figure 3) and fragment mixes for homology dependent assembly (Figure 5) were cleaned and concentrated with DNA Clean & Concentrator-5 columns (Zymo), and DNA was eluted with 10 μL molecular grade water. This elution was transformed into 50 μL of TOP10 competent cells. The other 5 μL was used to measure DNA concentration for transformation efficiency calculations with a NanoDrop-2000c Microvolume Spectrophotometer (Thermo Fisher). Transformation mixes were incubated for 30 min on ice. Mixes were then heat shocked at 42°C for 60 sec and recovered at 37°C for 60 min with 250 μL LB with Catabolite repression (LB + 0.2% (w/v) dextrose) medium added. Depending on the predicted assembly efficiency, 50-200 μL of the base mix or a 10X dilution was plated onto LB-agar plates with the appropriate antibiotics for incubation at 37°C overnight. Colonies were manually counted and reported as Colony-Forming Units per μg of DNA plated (CFU/μg). Plasmid assembly was assessed by chromoprotein expression, sanger sequencing aligned with A Plasmid Editor (ApE)(25), and whole plasmid sequencing (Plasmidsaurus).

Fragment mixes for Figure 4 were transformed into a separate batch of TOP10 competent cells and were recovered with Super Optimal Broth with Catabolite repression (SOC). These transformations had comparable transformation efficiency to those above, however Figure 4 transformation efficiency should not be directly compared with Figures 3 and 5.

### 2.4 Liquid handling protocol

DNA constructs were designed using j5 (File 3, File 4, File 5, File 6) (Figure 1A). Following design, a python setup script was run on the anaconda command prompt to launch AssemblyTron. First, the setup script used subprocess to call an R script included in AssemblyTron to parse the combinatorial j5 designs (Figure 1B) into 4 CSV files: oligo.csv, assembly.csv, combinations.csv, and pcr.csv (Figure 1C). Parsed CSV files were then automatically transferred from our shared google drive to the AssemblyTron working directory. The setup script then generates a custom text file with instructions on how to set up the deck for the current protocol (Table 7) (Figure 1D). Next, it prompts a pop-up window for specifying template concentrations, modifying PCR volumes, etc. and saves these parameters as another CSV in the working directory (Figure 1E). The setup script then runs the optimal annealing algorithm, saves the gradient as a CSV, and attaches tube positions to the pcr.csv file (Figure 1G). Finally, AssemblyTron calculates location, concentration, and volume of reagents and DNA and saves the information as CSV input files (Figure 1F). It then makes copies of the automated dilution and protocol scripts and produces another instructions file for positioning PCR tubes in the thermocycler (Table 8). A batch script is then called via subprocess from the setup script to copy all CSV input files to the robot working directory and is included in the package download. The resulting Protocol scripts in the working directory were then uploaded to the OT-2 run app where they have been thoroughly screened and debugged. The AssemblyTron Golden Gate script is based on Engler et al. (8), and the homology dependent assembly script is inspired by Garcia-Nafria et al. (7) and Beyer et al. (6,23). PCR reactions were manually transferred to a Bio-Rad C100 gradient thermocycler. Cycles were performed as stated above and the reactions were then returned to the OT-2 for assembly. Output files are archived in the working directory following the run (Figure 1H).

**Figure 1:**
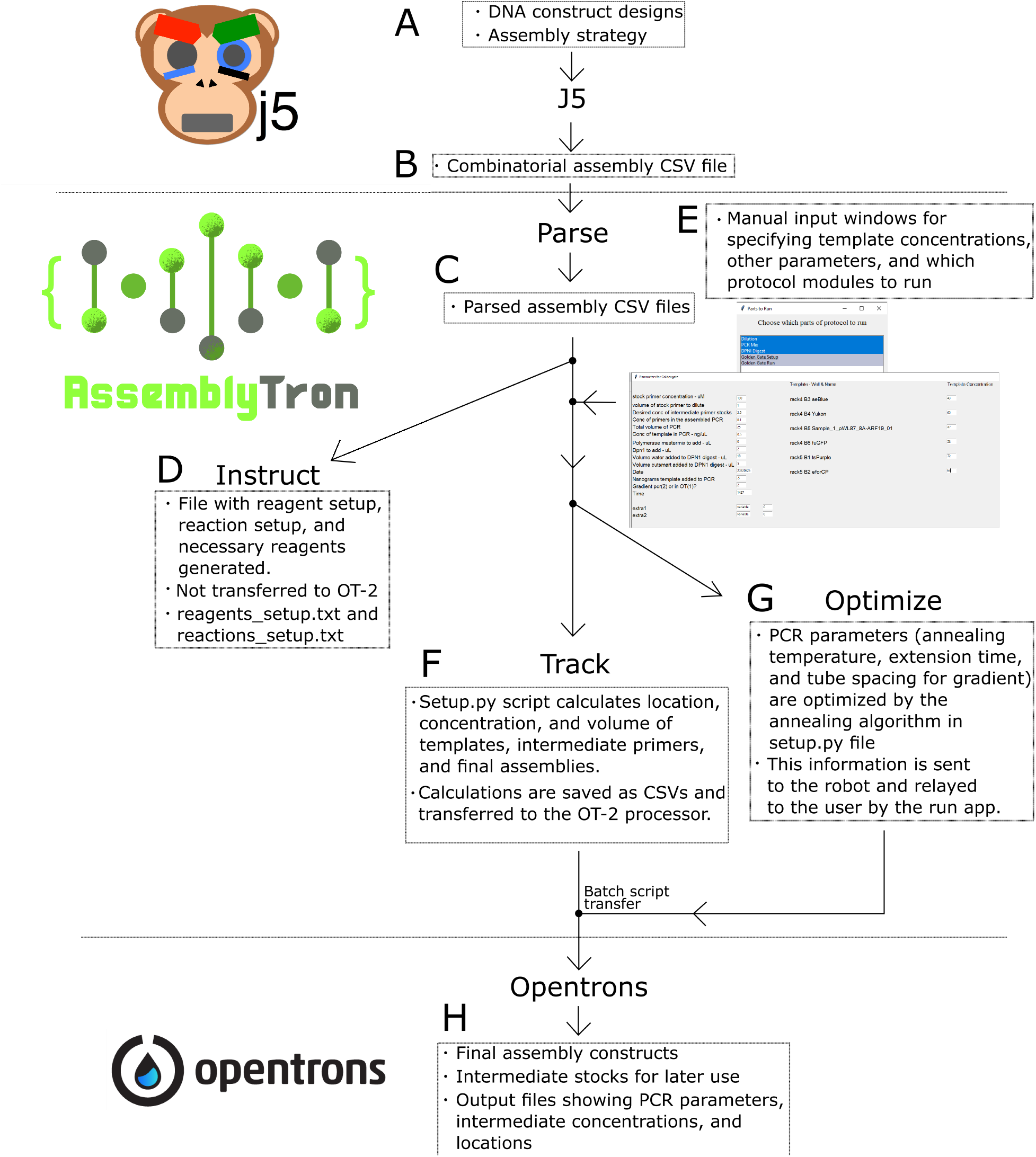
AssemblyTron Workflow: A) User must devise a DNA construct design and choose the appropriate assembly strategy (Golden Gate, Gibson, etc.). B) After putting together the design in J5, the user receives a combinatorial design file as an output from J5. C) AssemblyTron R script parses the J5 combinatorial assembly file and divides it up into separate CSV files for different stages of assembly. This script runs automatically in Setup.py script. D) After initiating the AssemblyTron Setup.py script, and specifying the location of the parsed design files, the user receives a reagents_setup file as an output (Table 7). E) The user is manually prompted to input template concentration, which parts of the protocol to run, etc. F) Setup.py calculates and tracks the location, concentration, and volume of all primers, templates, and final assemblies. This information is provided as CSV output files. G) AssemblyTron optimizes the annealing temperature and extension time of PCRs with its optimal annealing algorithm. Instruction for arranging 100 μL PCR tubes in the OT-2 thermocycler are provided in reactions_setup.txt. H) At the conclusion of the protocol run, AssemblyTron prompts the OT-2 to provide the user with output files containing all tracking information. The user is left with final assembly constructs as well as intermediate stocks.

## 3. Results

To demonstrate the utility and performance of the AssemblyTron package, we have used it to build two sets of combinatorial plasmid assemblies, one consisting of two yeast expression vectors containing plant transcription factors assembled from two fragments, and another consisting of four chromoprotein *E. coli* expression vectors assembled from four fragments each. The chromoprotein assemblies enable high-throughput quantification of the efficiency and accuracy of assembly.

To use AssemblyTron, the user must first begin with a combinatorial DNA construct design and select an appropriate assembly strategy (Figure 1A). This information is then used as an input for the j5 algorithm (18) which generates combinatorial assembly files (Figure 1B). Combinatorial assembly files specify the primer sequences, PCR parameters, and fragment assembly strategy. Several combinatorial assembly files for different sets of constructs from different users can be combined with j5 tools to batch many assemblies across a lab or biofoundry user base. AssemblyTron in its current iteration is limited to assemblies with a maximum of 96 total primers and templates.

AssemblyTron then converts combinatorial design files into Opentrons protocols and Instructions documents for the operator. A single combinatorial assembly file from j5 is used as the input to the AssemblyTron package. The user begins by initiating the Setup.py script in AssemblyTron, which facilitates all file processing. The combinatorial assembly file is first parsed by an AssemblyTron R script called by Setup.py into 4 individual CSV files: oligo.csv, pcr.csv, assembly.csv, and combinations.csv (Figure 1C). The script then moves the new CSV files to the AssemblyTron working directory and processes them to generate an instructions file (Figure 1D), which instructs the user which reagents to retrieve and how to set up the OT-2 deck for the run. Next, Setup.py prompts the user to input the concentrations of template stocks for dilutions (Figure 1E). Other parameters, such as PCR volumes, primer concentrations, etc. can be modified but are prefilled for convenience. Additionally, there are two slots in the graphical user interface for extra parameters in the case that a user modifies the source code.

Following parameter confirmation, Setup.py calculates concentrations for working stocks of templates, primers, intermediates, and final products for the run. This means that AssemblyTron tracks and records every item through each step of the reaction and saves the information as CSV input and output files (Figure 1H), which could in the future interface with lab inventory management systems. Setup.py then transfers protocol scripts to the OT-2 via a batch script (Figure 1 F). Setup.py also defines an optimal thermocycler gradient and block position (annealing temperature) for each PCR as well as an optimal extension time for the set of reactions (Figure 1G). Since this optimization algorithm relies on implementation of a gradient annealing step, the current OT-2 thermocycler module is insufficient for our workflow. The user must manually transfer PCR tubes to a separate thermocycler and manually enter the PCR parameters; however, PCR tube spacing and gradient parameters are calculated and given by AssemblyTron in the reaction instructions text file. Alternatively, during the assembly design steps in j5, the user may specify stringent primer parameters (*i*.*e*. Primer Max Tm Diff, Primer Max TM, and Primer Min TM), although this method is less robust to difficult sequences. PCR products may either be purified or directly returned to the OT-2, and the assembly protocol is performed. The user is then left with assembled constructs to be transformed, remaining intermediate dilutions for later use, and output files in which each step of the protocol and location of reagents is tracked (Figure 1H).

### 3.1 PCR

First, we demonstrate the software’s ability to automatically prepare PCRs directly from input files (File 3, File 4, File 5, File 6). AssemblyTron is currently capable of performing PCRs with up to 96 combined primers and templates in standard microcentrifuge tubes which are diluted into a 96 well plate. Our script first calculates an optimal annealing temperature gradient that accommodates the annealing temperature of each fragment for assembly within 0.4 °C. J5 also includes a *delta* parameter, which specifies the tolerable variance from primer annealing temperatures for each PCR. If the delta parameter is less than 0.4 °C, the script will adjust the gradient to where these more rigid annealing temperatures are accommodated. The annealing temperature gradient accommodates each PCR in a single run. A reaction tube number is assigned to each temperature step in the gradient in order to specify tube placement (Table 8). These PCR tube positions are relayed to the user via a pop-up text file. The script uses the gradient to choose the tube positions, and once reactions are mixed, they are ready for gradient amplification. The OT-2 thermocycler module does not currently have gradient capability, so it is necessary to move mixed reactions into a separate gradient thermocycler to run the reaction. However, all calculations, optimization, and setup for the run are determined by our software package. This reduces the cognitive burden of molecular cloning, reduces the rate of failure, and eliminates need for any manual protocol optimization. Finally, our package prompts the Opentrons user interface to pause and instruct the user how to set up the gradient PCR run. The main drawback of PCR is the trial-and-error optimization process, and our package harnesses machine precision to automate and resolve PCR parameter calculation using j5 design output files.

To demonstrate our PCR script and gradient algorithm, we amplified 5 fragments with variable sizes and annealing temperature means and differences (Figure 2A). Our setup.py script contains the gradient algorithm, working directory generation code, and parameter input window for staging the protocol. Input files can be found in supplementary Tables 2-6 (File 3, File 4, File 5, File 6).

**Figure 2:**
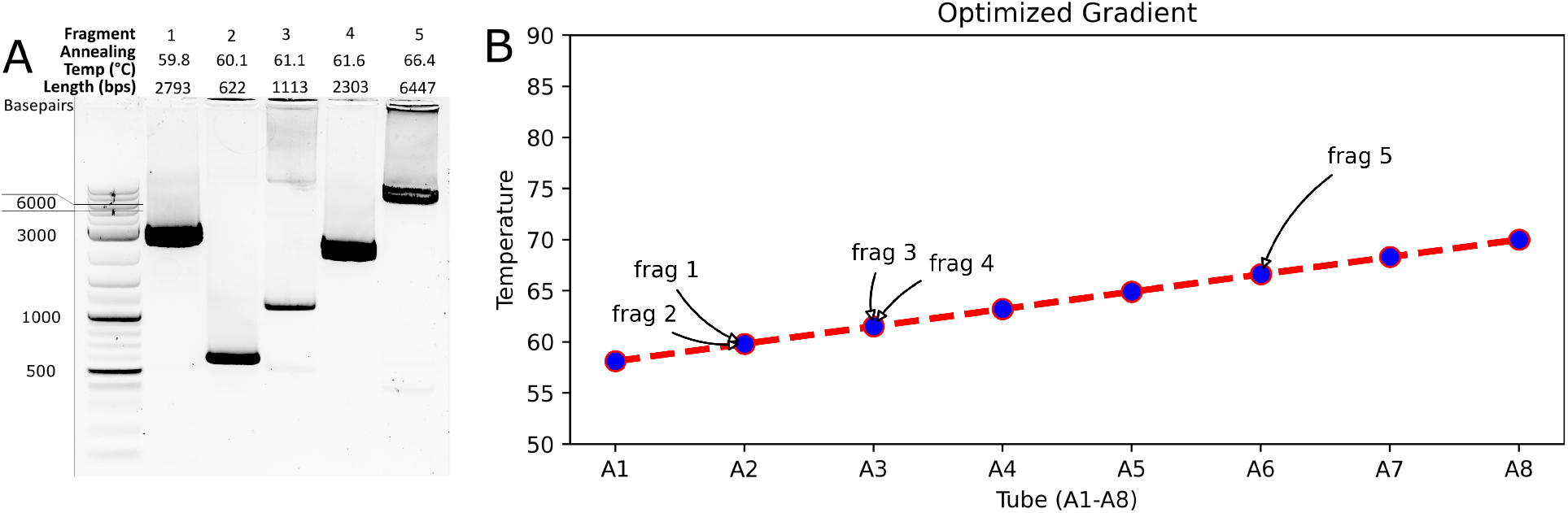
PCR with AssemblyTron (A) AssemblyTron PCR protocol was verified by successful amplification of DNA fragments with variable annealing temperatures and lengths. (B) A graphical depiction of the optimal annealing temperature gradient calculated for the PCR. Each tube is assigned a temperature in the linear gradient, which corresponds to an annealing temperature of one or more fragments.

Gel electrophoresis of final PCR products indicated that we successfully amplified all fragments and obtained the expected sizes from a single gradient thermocycler run (Figure 2A). It should be noted that fragment 5 had a significantly higher annealing temperature (>4 degrees) than the other four fragments. To demonstrate how our algorithm accommodates this difference, Figure 2B provides a visual plot of the optimal linear temperature gradient calculated by our algorithm. The algorithm determined that fragment 5 should be in a tube located in row A at column 6 which is higher in the temperature gradient than the other fragments. This information was then relayed to the robot which assembled the reaction mix in the correct position. The reactions are transferred to the thermocycler, and the gradient cycle was set as outlined in the instructions document (Table 8). All reactions were performed successfully as shown in Figure 2A.

### 3.2 Golden Gate Assembly

AssemblyTron also includes a Golden Gate assembly feature, which allows the user to go from primers and templates specified in a j5 Golden Gate assembly file to fully assembled, transformation-ready plasmids in a single day. This feature begins by performing a gradient-optimized PCR run for each fragment in separate tubes. Primers and templates specified from j5 design files are used for the PCR run (Figure 3A). Next, a Dpn1 digestion is performed to remove residual template. Our package then prompts the robot to automatically combine amplified fragments in equimolar ratios for each assembly reaction, assuming equivalent yields from each PCR. If any reactions of shorter fragments are likely too concentrated, a dilution will be performed to accommodate accurately pippetable equimolar additions of each fragment. Next, reagents are added prior to automatic initiation of the assembly in the Opentrons thermocycler module. The Opentrons thermocycler module is practical for Golden Gate assemblies since there is no parameter gradient required across reactions, in contrast to PCR. Although there is no need to relocate the reaction to a separate thermocycler, the protocol will pause so that reactions can be transferred in case the lab does not have access to the Opentrons thermocycler module. Once the Golden Gate thermocycler protocol is completed, the reactions can be removed, optionally cleaned and concentrated, and transformed into *E. coli*. Golden Gate assembly reactions performed using AssemblyTron are accurate and efficient (Figure 3D).

**Figure 3:**
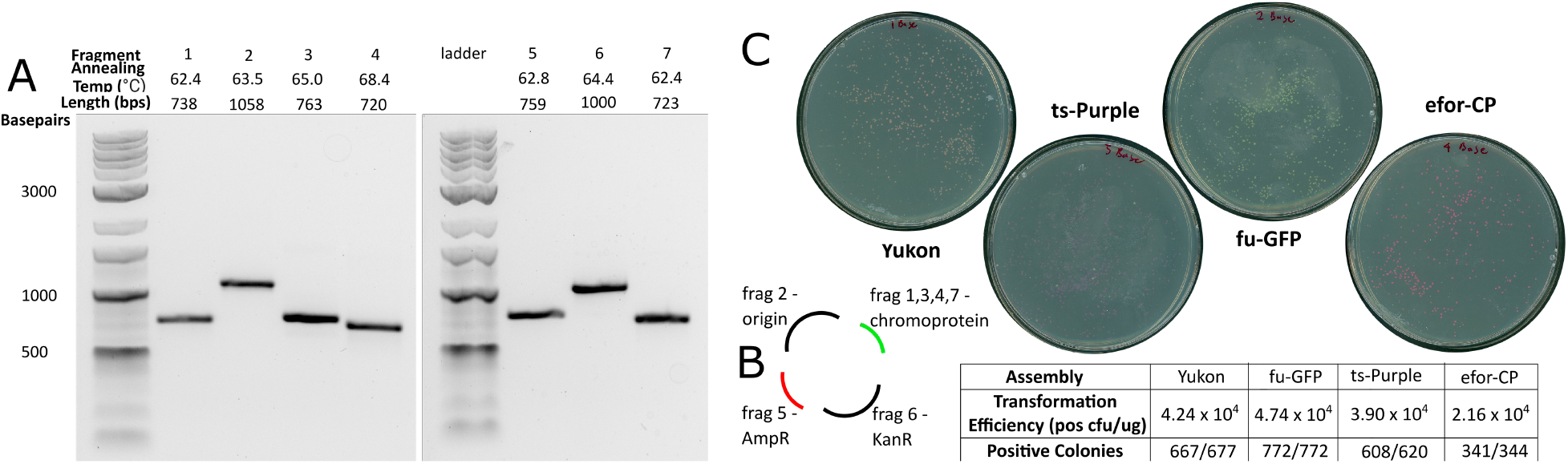
Golden Gate Assembly with AssemblyTron (A) AssemblyTron protocol was verified by successful amplification of each chromoprotein assembly fragment. (B) Schematic to specify how fragments are assembled to yield final constructs. (C) High transformation efficiency, high accuracy, and correct chromoprotein expression validate the robustness of the Golden Gate protocol.

To test the AssemblyTron combinatorial Golden Gate assembly feature, we also assembled chromoprotein *E. coli* expression vectors using this method. The assembly consists of four final plasmids, which were each assembled from 4 fragments. The fragments include two separate plasmid backbone fragments, a kanamycin selection gene, and the chromoprotein gene, which differs in each assembly. First, PCR reagents for each respective fragment were mixed and annealing temperature, and extension times were calculated using the PCR optimization algorithm (Figure 2C). After thermal cycling, samples of each reaction were removed for gel electrophoresis. Amplified fragments of the correct lengths are shown in figure 3A where fragments 2, 5, and 6 are backbone parts, and fragments 1, 3, 4, and 7 are chromoprotein parts (Figure 3B).

Following fragment amplification, the reactions were returned to the Opentrons robot where AssemblyTron automatically performed a Dpn1 digestion to eliminate residual template DNA. Before proceeding to assembly step, each fragment was cleaned and concentrated to remove polymerase, which we found to interfere with assembly. To do this, the protocol was paused, fragments were removed for cleaning, and then they were returned to their positions on the thermocycler block. In future work, we plan to add an automated magnetic bead purification step. Next, a volume proportional to the fragment length for each resulting fragment was added to the respective assembly reaction in a new well. Again, for simplicity, we assumed that each fragment PCR is similarly efficient and so this volume proportional to length will result in a roughly equimolar mixture of fragments. This avoids the need for further purification and quantification of DNA in most cases. Following the assembly, constructs were transformed into competent *E. coli* (produced in-house with transformation efficiency greater than 10^8^ CFU per μg pUC19)(24). Between 300 and 800 colonies were formed from each assembly with transformation efficiencies on the order of 10^4^ CFU per ug of total assembly reaction DNA transformed. Greater than 98% of colonies were positive based on chromoprotein expression. The Golden Gate Assembly feature also includes the option to use a destination plasmid that contains internal BsaI sites specifically for Golden Gate cloning, as opposed to assembling the plasmid backbone from PCR amplified fragments.

### 3.3 Short Homology Based Assembly

Since Gibson assembly is the most popular cloning strategy(26), we provide AssemblyTron users with a script to accommodate this approach. However, instead of creating a Gibson protocol with a lengthy assembly cycle requiring more reagents and time, we use a homology dependent assembly strategy inspired by the *in vivo* assembly technique (IVA) (7) and AQUA cloning (6). Similar to Gibson, IVA and AQUA cloning are based on recombination 20+ bps of homology between the ends of adjacent fragments. However, these strategies do not depend on enzyme-catalyzed creation of sticky-ends, instead relying on assembly via homologous recombination *in vivo*, in *E. coli*. While we have successfully assembled two-fragment constructs in one-pot PCRs, as specified in IVA cloning (7), we experienced consistent failure when attempting to assemble constructs with more than two fragments in one-pot. Perhaps setting tighter tolerances for primer annealing temperature in the initial j5 design would improve one-pot fragment amplification. In order to ensure reproducibility and consistent success, we also developed a script to generate protocols using separate PCRs for each fragment as in AQUA cloning (6). The protocol for one-pot IVA reactions is still available in AssemblyTron, but we recommend the separate PCR script for more complex designs. In these protocols, following thermal cycling, PCRs, either combined or separate are digested with Dpn1. For separate fragment PCRs the products are then combined using volumes proportional to fragment length. This fragment mixture is then transformed into *E. coli* where native E. coli machinery was responsible for assembling fragments.

To demonstrate short homology based assembly with AssemblyTron, we performed two sets of combinatorial assemblies. First, we did a one-pot IVA to implement site-directed mutagenesis on a construct frequently used in our lab. After initial PCR amplification of fragments and Dpn1 digestion to eliminate residual template, we analyzed a small fraction of these reactions by gel electrophoresis to confirm correct fragment sizes (Figure 4A). Since both fragments in lane 1 are approximately the same size, they are indistinguishable. However, fragment sizes in lane 2 differed by approximately 1000 bp, and slight separation between bands can be seen in Figure 4A, indicated by cyan arrows and the fragment number. After amplification, Dpn1 digestion, and column purification, we transformed these fragment mixtures into *E. coli*. We recorded transformation efficiencies on the order of approximately 10^2^ CFU/μg for these IVA reactions (Figure 4B). Sequence verification determined that 2/6 colonies contained the correct mutation for assembly 1, and 3/6 colonies were correct for assembly 2. Incorrect colonies either had the original sequence or random insertions and deletions (supplementary file 9 and file 10).

**Figure 4:**
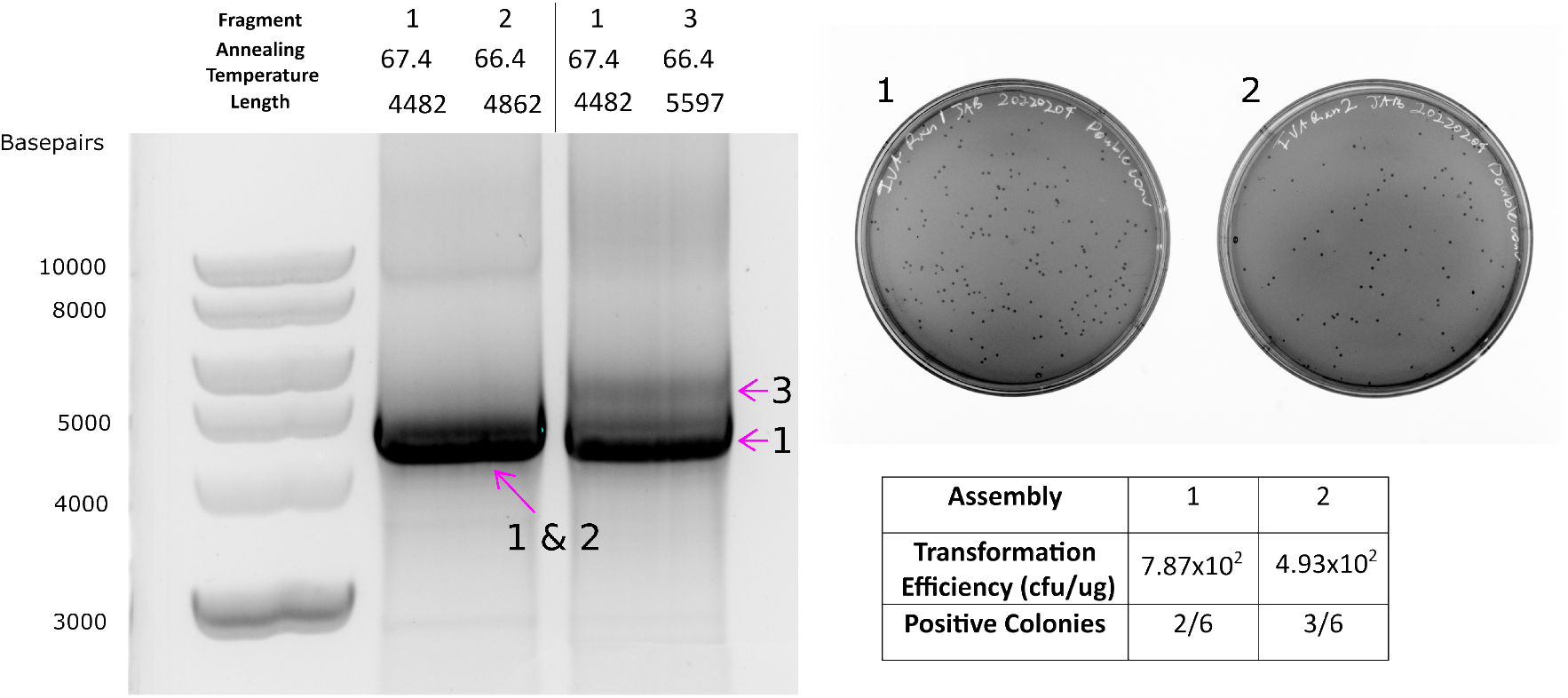
Homology dependent one-pot *in Vivo* Assembly (IVA) with AssemblyTron. A) AssemblyTron one-pot IVA assembly was verified by amplification of appropriate bands via gel electrophoresis. Fragments 1 and 2 are indistinguishable due to size similarity, however there is a slight resolution between fragments 1 and 3. B) Consistent transformation efficiency and sequence verification validate the one-pot IVA protocol.

In addition, we performed the same chromoprotein assembly described in figure 3 using the AQUA assembly strategy. Correct bands were amplified (Figure 5A), fragment mixes were transformed into *E. coli*, and dilutions were plated to determine transformation efficiency. We recorded transformation efficiencies on the order of approximately 10^3^ cfu/μg for these IVA assemblies (Figure 5C). Recombination-based assembly techniques are characteristically lower in efficiency than restriction-ligation-based assemblies like Golden Gate, so these results were expected (27). Chromoprotein expression revealed that 68.7% of colonies in the Yukon assembly were positive for the Yukon chromoprotein, and >80% of colonies were positive for the remaining colors. Negative colonies expressed either the blue chromoprotein from the template backbone vector or no color.

**Figure 5:**
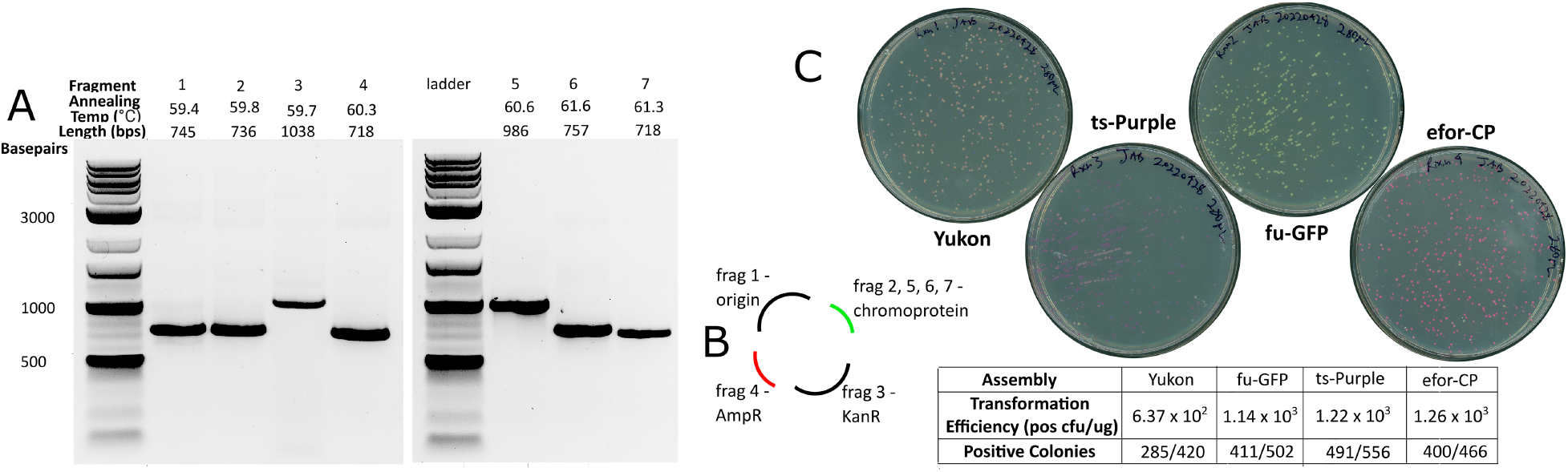
Homology dependent AQUA Assembly with AssemblyTron. (A) AssemblyTron AQUA protocol was verified by successful amplification of each chromoprotein assembly fragment. (B) Schematic to specify how fragments are assembled to yield final constructs. (C) Consistent transformation efficiency, correct antibiotic resistance, and correct chromoprotein expression further validate the AQUA protocol.

## 4. Discussion

The error-prone nature of molecular cloning inhibits the training of a large, robust workforce of synthetic biologists. AssemblyTron provides an open-source framework for automating cloning and is a tool to improve the productivity of labs plagued by failing workflows and limited funds (22). Its simple interface can be operated by an undergraduate student with minimal prior cloning experience. We demonstrate that our software is capable of seamlessly integrating with j5 DNA assembly designs. Thus, paired with j5, AssemblyTron provides reproducible algorithmic solutions to the design and build steps of the Design-Build-Test-Learn cycle ubiquitous to synthetic biology. AssemblyTron’s ability to perform optimized PCRs and efficient DNA assemblies will allow researchers to spend more time asking questions and learning from experiments instead of scrutinizing error-prone cloning. AssemblyTron will also lighten the training barrier for undergraduates, since all calculations and nuanced details are automatically handled by AssemblyTron.

Our data provides evidence that AssemblyTron delivers robust results for cloning workflows. We successfully performed PCRs with variable parameters and multi-part assemblies via both IVA and Golden Gate. Assemblies performed with AssemblyTron in the OT-2 have similar transformation efficiency and accuracy to the manual procedures.

AssemblyTron is open source, under the Apache License 2.0, so any public lab interested in automating their workflow has access. The OT-2 liquid handling robot is one of the most affordable and can currently be purchased, complete with temperature module, magnetic module, and thermocycler module, for $16,500. This price point is ten times less expensive than similar systems (28). By reducing opportunities for human error, decreasing human labor time, and increasing throughput, automated cloning with AssemblyTron and the OT-2 can realistically save time and money in the long run.

Although AssemblyTron has expedited cloning in our lab and has the potential to do so in other labs as well, there are still numerous opportunities for improvement and expansion. The complexity of assemblies that can currently be handled by the AssemblyTron software is limited by our 96 combined primer and template capacity. However, as demand for multiplex assemblies increases, we will scale our platform to accommodate 384 well plates for the primer and template dilution steps. This modification will be simple to implement and serves as an example of the scalability of AssemblyTron.

Another drawback is that in the Golden Gate protocol, amplified fragments must be cleaned and concentrated before mixing with restriction enzymes and ligase, to prevent sticky ends from being filled in by residual polymerase. In the future, we will integrate the magnetic module to perform DNA cleaning within the OT-2 to avoid the use of expensive, time-consuming, and human error-prone manual purification kits. Several open-source protocols for magnetic bead purification already exist for the OT-2 (29), which can be integrated in future versions of AssemblyTron.

Additionally, we are in the early stages of integrating a transformation protocol into AssemblyTron, which has also been achieved in an automated context in the past by DNA-BOT (20). This will save even more time for researchers if transformations can be performed directly in the OT-2. The more we can automate, the more time we will have to develop the field of synthetic biology.

## Material Availability Statement

The software, AssemblyTron, used to produce the results of this study is openly available on Zenodo at http://doi.org/10.5281/zenodo.7062951. The latest iteration of AssemblyTron is available on PyPI at https://pypi.org/project/AssemblyTron/.

## Data Availability Statement

The authors confirm that the data supporting the findings of this study are available within the article and its supplementary materials.

## Acknowledgments

We thank Jennifer L Nemhauser and Sebastian Cocioba for providing plasmids used in this work. Research in the Wright Plant Synthetic Biology Laboratory is supported by the USDA NIFA AFRI Plant Breeding for Agricultural Production GRANT13367799 and Hatch Project VA-1021738, Virginia Space Grant Consortium, VT Institute for Critical Technologies and Sciences Junior Faculty Award, and VT CALS Strategic Plan Advancement 2021 Integrated Internal Competitive Grants through the Center for Advanced Innovation in Agriculture.

## Author contributions

J.A.B and R.C.W. conceived of the project and designed experiments. All authors wrote code and revised code for the software package. J.A.B acquired data. J.A.B and R.C.W. analyzed and interpreted data. J.A.B and R.C.W drafted and revised the manuscript. All authors approved of the final manuscript.

## References

1. Casini A, Storch M, Baldwin GS, Ellis T. Bricks and blueprints: methods and standards for DNA assembly. Nat Rev Mol Cell Biol. 2015 Sep;16(9):568–76.

2. Appleton E, Densmore D, Madsen C, Roehner N. Needs and opportunities in bio-design automation: four areas for focus. Current Opinion in Chemical Biology. 2017 Oct 1;40:111– 8.

3. Li MZ, Elledge SJ. SLIC: A Method for Sequence- and Ligation-Independent Cloning. In: Peccoud J, editor. Gene Synthesis [Internet]. Totowa, NJ: Humana Press; 2012 [cited 2022 Sep 9]. p. 51–9. (Methods in Molecular Biology; vol. 852). Available from: http://link.springer.com/10.1007/978-1-61779-564-0_5

4. Gibson DG. Synthesis of DNA fragments in yeast by one-step assembly of overlapping oligonucleotides. Nucleic Acids Research. 2009 Nov 1;37(20):6984–90.

5. Quan J, Tian J. Circular Polymerase Extension Cloning of Complex Gene Libraries and Pathways. PLOS ONE. 2009 Jul 30;4(7):e6441.

6. Beyer HM, Gonschorek P, Samodelov SL, Meier M, Weber W, Zurbriggen MD. AQUA Cloning: A Versatile and Simple Enzyme-Free Cloning Approach. PLOS ONE. 2015 Sep 11;10(9):e0137652.

7. Garcĺa-Nafrĺa J, Watson JF, Greger IH. IVA cloning: A single-tube universal cloning system exploiting bacterial In Vivo Assembly. Scientific Reports. 2016 Jun 6;6(1):27459.

8. Engler C, Gruetzner R, Kandzia R, Marillonnet S. Golden Gate Shuffling: A One-Pot DNA Shuffling Method Based on Type IIs Restriction Enzymes. PLOS ONE. 2009 May 14;4(5):e5553.

9. Kosuri S, Church GM. Large-scale de novo DNA synthesis: technologies and applications. Nat Methods. 2014 May;11(5):499–507.

10. Palluk S, Arlow DH, de Rond T, Barthel S, Kang JS, Bector R, et al. De novo DNA synthesis using polymerase-nucleotide conjugates. Nat Biotechnol. 2018 Aug;36(7):645–50.

11. Chao R, Liang J, Tasan I, Si T, Ju L, Zhao H. Fully Automated One-Step Synthesis of Single-Transcript TALEN Pairs Using a Biological Foundry. ACS Synth Biol. 2017 Apr 21;6(4):678–85.

12. Edinburgh Genome Foundry [Internet]. The University of Edinburgh. [cited 2022 Jun 3]. Available from: https://www.ed.ac.uk/biology/research/facilities/edinburgh-genome-foundry

13. Hillson N, Caddick M, Cai Y, Carrasco JA, Chang MW, Curach NC, et al. Building a global alliance of biofoundries. Nat Commun. 2019 May 9;10(1):2040.

14. Vrana J, de Lange O, Yang Y, Newman G, Saleem A, Miller A, et al. Aquarium: open-source laboratory software for design, execution and data management. Synthetic Biology. 2021 Feb 17;6(1):ysab006.

15. Walsh DI, Pavan M, Ortiz L, Wick S, Bobrow J, Guido NJ, et al. Standardizing Automated DNA Assembly: Best Practices, Metrics, and Protocols Using Robots. SLAS TECHNOLOGY: Translating Life Sciences Innovation. 2019 Jun 1;24(3):282–90.

16. Hérisson J, Duigou T, du Lac M, Bazi-Kabbaj K, Sabeti Azad M, Buldum G, et al. The automated Galaxy-SynBioCAD pipeline for synthetic biology design and engineering. Nat Commun. 2022 Aug 29;13(1):5082.

17. Tegally H, San JE, Giandhari J, de Oliveira T. Unlocking the efficiency of genomics laboratories with robotic liquid-handling. BMC Genomics. 2020 Dec;21(1):729.

18. Hillson NJ, Rosengarten RD, Keasling JD. j5 DNA Assembly Design Automation Software. ACS Synthetic Biology. 2012;8.

19. Jones TS, Oliveira SMD, Myers CJ, Voigt CA, Densmore D. Genetic circuit design automation with Cello 2.0. Nat Protoc. 2022 Apr;17(4):1097–113.

20. Storch M, Haines MC, Baldwin GS. DNA-BOT: a low-cost, automated DNA assembly platform for synthetic biology. Synthetic Biology. 2020 Jan 1;5(1):ysaa010.

21. Storch M, Casini A, Mackrow B, Fleming T, Trewhitt H, Ellis T, et al. BASIC: A New Biopart Assembly Standard for Idempotent Cloning Provides Accurate, Single-Tier DNA Assembly for Synthetic Biology. ACS Synth Biol. 2015 Jul 17;4(7):781–7.

22. Bryant J, plantsynbio, mkellinger, CameronLongmire, Wright C, RCMiller-PSB, et al. PlantSynBioLab/AssemblyTron: AssemblyTron [Internet]. Zenodo; 2022 [cited 2022 Sep 9]. Available from: https://zenodo.org/record/7062951

23. Green MR, Sambrook J, Sambrook J. Molecular cloning: a laboratory manual. 4th ed. Cold Spring Harbor, N.Y: Cold Spring Harbor Laboratory Press; 2012. 3 p.

24. Hanahan D, Jessee J, Bloom FR. [4] Plasmid transformation of Escherichia coli and other bacteria. In: Methods in Enzymology [Internet]. Academic Press; 1991 [cited 2022 May 15]. p. 63–113. (Bacterial Genetic Systems; vol. 204). Available from: https://www.sciencedirect.com/science/article/pii/007668799104006A

25. Davis MW, Jorgensen EM. ApE, A Plasmid Editor: A Freely Available DNA Manipulation and Visualization Program. Frontiers in Bioinformatics [Internet]. 2022 [cited 2022 Sep 12];2. Available from: https://www.frontiersin.org/articles/10.3389/fbinf.2022.818619

26. Kahl LJ, Endy D. A survey of enabling technologies in synthetic biology. J Biol Eng. 2013;7(1):13.

27. Pryor JM, Potapov V, Pokhrel N, Lohman GJS. Rapid 40 kb genome construction from 52 parts [Internet]. Synthetic Biology; 2020 Dec [cited 2022 Jul 12]. Available from: http://biorxiv.org/lookup/doi/10.1101/2020.12.22.424019

28. Opentrons | Open-source Lab Automation, starting at $5,000 [Internet]. [cited 2022 Jun 5]. Available from: https://opentrons.com/

29. Magnetic Module GEN2 | Opentrons OT-2 Modular Hardware [Internet]. [cited 2022 Jul 12]. Available from: https://opentrons.com/modules/magnetic-module/

