## Supplementary material for "AssemblyTron: Flexible automation of DNA assembly with Opentrons OT-2 lab robots": Table 7

Date: 20220829 Time: 1605

Absolute Path: C:\Users\Public\Documents\opentrons\src\AssemblyTron\Golden_Gate

Place the coldtuberack in slot 1.

Put 300uL tips in slot 6 and 9, and 10uL tips in slot 5.

Put oWL00375_(backbone)_forward in deckslot4 A1

Put oWL00376_(backbone)_reverse in deckslot4 A2

Put oWL00377_(Yukon)_forward in deckslot4 A3

Put oWL00378_(Yukon)_reverse in deckslot4 A4

Put oWL00379_(bb2)_forward in deckslot4 A5

Put oWL00380_(bb2)_reverse in deckslot4 A6

Put oWL00381_(AmpR)_forward in deckslot5 A1

Put oWL00382_(AmpR)_reverse in deckslot5 A2

Put oWL00383_(fuGFP)_forward in deckslot5 A3

Put oWL00384_(fuGFP)_reverse in deckslot5 A4

Put oWL00385_(tsPurple)_forward in deckslot5 A5

Put oWL00386_(tsPurple)_reverse in deckslot5 A6

Put oWL00387_(eforCP)_forward in deckslot4 B1

Put oWL00388_(eforCP)_reverse in deckslot4 B2

NOTE: if a template is listed twice, (ie, pwl106 in B6 and C3) then skip the second position, and move remaining templates up a slot

This is ok because this setup sheet and df object in the script are both set up from pcr.csv, except df just takes out repeasts.

Put aeBlue in deckslot4 B3

Put Yukon in deckslot4 B4

Put Sample_1_pWL87_8A-ARF19_01 in deckslot4 B5

Put fuGFP in deckslot4 B6

Put tsPurple in deckslot5 B1

Put eforCP in deckslot5 B2

After Dilutions, place the following reagets in 24 tube rack in slot 1.

Place empty tube in C4 for the T4/BSA mix

Place T4 ligase in C5

Place 100X BSA in C6

Place T4 buffer in D2

Place DPNI in D3

Place cutsmart buffer in D4

Place BsaI in D5

Place Q5 DNA polymerase in D6

Place 24 well tuberack in slot 2. Add 27 empty 1.5 mL tubes to the rack in the same positions.

**Table 7: reagent_setup.txt example file with instructions for setting up the OT-2 deck and slots. This file corresponds to the chromoproteins assembly in Figure 3.**
