## Supplementary material for "AssemblyTron: Flexible automation of DNA assembly with Opentrons OT-2 lab robots": Table 8

PCR gradient tube positions:

Date: 20220829 Time: 1605

Absolute Path:

C:\Users\Public\Documents\opentrons\src\AssemblyTron\Golden_Gate\202208291605_GoldenGate

Put a 100 uL PCR tube in A4

Put a 100 uL PCR tube in A2

Put a 100 uL PCR tube in A3

Put a 100 uL PCR tube in B4

Put a 100 uL PCR tube in B2

Put a 100 uL PCR tube in C2

Put a 100 uL PCR tube in A7

Final assembly 100 uL PCR tube:

Put a 100 uL PCR tube in B8

Put a 100 uL PCR tube in B9

Put a 100 uL PCR tube in B10

Put a 100 uL PCR tube in B11

Fragment dilution 100 uL PCR tubes:

Put a 100 uL PCR tube in F4

Put a 100 uL PCR tube in F3

Put a 100 uL PCR tube in F2

Put a 100 uL PCR tube in G4

Put a 100 uL PCR tube in F4

Put a 100 uL PCR tube in F3

Put a 100 uL PCR tube in G2

Put a 100 uL PCR tube in G4

Put a 100 uL PCR tube in F4

Put a 100 uL PCR tube in F3

Put a 100 uL PCR tube in H2

Put a 100 uL PCR tube in G4

Put a 100 uL PCR tube in F4

Put a 100 uL PCR tube in F3

Put a 100 uL PCR tube in F7

Put a 100 uL PCR tube in G4

**Table 8. reactions_setup.txt example file with instructions for setting up the thermocycler with 100 µL PCR tubes. This file corresponds to the chromoproteins assembly in Figure 3.**
