## Supplementary material for "AssemblyTron: Flexible automation of DNA assembly with Opentrons OT-2 lab robots": Table 2

| **Primary Templates** | **Antibiotic Resistance** | | **Color** | **Figure Reference** |
| --- | --- | --- | --- | --- |
| aeBlue | Kanamycin | Blue | | 3, 5 |
| Yukon | Kanamycin | Orange | | 3, 5 |
| Sample_1_pWL87_8A_ARF19_01 | Ampicillin | N/A | | 3, 5 |
| fuGFP | Kanamycin | Green | | 3, 5 |
| tsPurple | Kanamycin | Purple | | 3, 5 |
| eforCP | Kanamycin | Pink | | 3, 5 |
| pWL242_pGP8A_ARF7pdar | Ampicillin | N/A | | 4 |
| pWL241_pGP8A_ARF5pdar | Ampicillin | N/A | | 4 |
| pGP8A_ARF7 | Ampicillin | N/A | | 2 |

**Table 2. List of templates used for j5 assembly designs.**
