## Supplementary material for "AssemblyTron: Flexible automation of DNA assembly with Opentrons OT-2 lab robots": Table 1

| **Name** | **Length** | **Tm** | **Tm (3')** | **Cost (USD)** | **Sequence** | **Sequence (3' only)** |
| --- | --- | --- | --- | --- | --- | --- |
| oWL00375_(backbone)_forward | 49 | 65.3 | 64.7 | 4.9 | CACACCAGGTCTCAAAAAGCTTTATAGATTACAGTCGACAGATCAAAGG | AAGCTTTATAGATTACAGTCGACAGATCAAAGG |
| oWL00376_(backbone)_reverse | 48 | 67.0 | 65.3 | 4.8 | CACACCAGGTCTCAATGGGACTCTTTCTCCTCTTTAATCTCTAGTAGC | GGGACTCTTTCTCCTCTTTAATCTCTAGTAGC |
| oWL00377_(Yukon)_forward | 38 | 73.9 | 61.5 | 3.8 | CACACCAGGTCTCACCATGGCTTCCCTGTCAAAACAAG | ATGGCTTCCCTGTCAAAACAAG |
| oWL00378_(Yukon)_reverse | 43 | 70.6 | 63.4 | 4.3 | CACACCAGGTCTCACTCTTATCAATGGTGATGGTGGTGATGTG | TTATCAATGGTGATGGTGGTGATGTG |
| oWL00379_(bb2)_forward | 38 | 74.5 | 65.5 | 3.8 | CACACCAGGTCTCAAGAGCTCAAAAAAAAACCCCGCCC | GAGCTCAAAAAAAAACCCCGCCC |
| oWL00380_(bb2)_reverse | 47 | 63.9 | 61.6 | 4.7 | CACACCAGGTCTCAGTCGACTGTAATCTATAAAGCTTTTAGAAAAAC | GTCGACTGTAATCTATAAAGCTTTTAGAAAAAC |
| oWL00381_(AmpR)_forward | 45 | 71.1 | 63.5 | 4.5 | CACACCAGGTCTCACGACCGCGGAACCCCTATTTGTTTATTTTTC | CGCGGAACCCCTATTTGTTTATTTTTC |
| oWL00382_(AmpR)_reverse | 43 | 70.0 | 65.3 | 4.3 | CACACCAGGTCTCATTTTACCAATGCTTAATCAGTGAGGCACC | TTACCAATGCTTAATCAGTGAGGCACC |
| oWL00383_(fuGFP)_forward | 34 | 76.4 | 61.8 | 3.4 | CACACCAGGTCTCACCATGGTGAGCAGTGGCGAA | ATGGTGAGCAGTGGCGAA |
| oWL00384_(fuGFP)_reverse | 47 | 67.8 | 63.8 | 4.7 | CACACCAGGTCTCACTCTTATTAGCTCTTATACAGCTCGTCCATACC | TTATTAGCTCTTATACAGCTCGTCCATACC |
| oWL00385_(tsPurple)_forward | 40 | 73.2 | 63.1 | 4 | CACACCAGGTCTCACCATGGCGAGCTTAGTCAAGAAAGAC | ATGGCGAGCTTAGTCAAGAAAGAC |
| oWL00386_(tsPurple)_reverse | 41 | 70.8 | 61.7 | 4.1 | CACACCAGGTCTCACTCTCATTAAGTTGCTTTCTCAGGCAC | TCATTAAGTTGCTTTCTCAGGCAC |
| oWL00387_(eforCP)_forward | 46 | 70.9 | 66.8 | 4.6 | CACACCAGGTCTCACCATGGCAGTGATTAAGCAGGTAATGAAGACC | ATGGCAGTGATTAAGCAGGTAATGAAGACC |
| oWL00388_(eforCP)_reverse | 44 | 74.2 | 70.1 | 4.4 | CACACCAGGTCTCACTCTTATTATGGGAGAGCCTTCGGCAGCGG | TTATTATGGGAGAGCCTTCGGCAGCGG |
| oWL00357_(backbone)_forward | 35 | 65.9 | 58.6 | 3.5 | GCATTGGTAAAAGCTTTATAGATTACAGTCGACAG | AAGCTTTATAGATTACAGTCGACAG |
| oWL00358_(backbone)_reverse | 29 | 64.4 | 60.3 | 2.9 | CCATGGGACTCTTTCTCCTCTTTAATCTC | GGGACTCTTTCTCCTCTTTAATCTC |
| oWL00359_(Yukon)_forward | 34 | 71.1 | 61.5 | 3.4 | AGAAAGAGTCCCATGGCTTCCCTGTCAAAACAAG | ATGGCTTCCCTGTCAAAACAAG |
| oWL00360_(Yukon)_reverse | 40 | 67.8 | 58.1 | 4 | GGGGTTTTTTTTTGAGCTCTTATCAATGGTGATGGTGGTG | TTATCAATGGTGATGGTGGTG |
| oWL00361_(bb2)_forward | 22 | 62.6 | 61.3 | 2.2 | AGAGCTCAAAAAAAAACCCCGC | GAGCTCAAAAAAAAACCCCGC |
| oWL00362_(bb2)_reverse | 35 | 68.3 | 58.1 | 3.5 | GTTCCGCGGTCGACTGTAATCTATAAAGCTTTTAG | GTCGACTGTAATCTATAAAGCTTTTAG |
| oWL00363_(AmpR)_forward | 26 | 70.4 | 59.9 | 2.6 | CAGTCGACCGCGGAACCCCTATTTGT | CGCGGAACCCCTATTTGT |
| oWL00364_(AmpR)_reverse | 36 | 66.8 | 61.3 | 3.6 | TCTATAAAGCTTTTACCAATGCTTAATCAGTGAGGC | TTACCAATGCTTAATCAGTGAGGC |
| oWL00365_(fuGFP)_forward | 30 | 72.5 | 61.8 | 3 | AGAAAGAGTCCCATGGTGAGCAGTGGCGAA | ATGGTGAGCAGTGGCGAA |
| oWL00366_(fuGFP)_reverse | 45 | 68.2 | 61.3 | 4.5 | GGGGTTTTTTTTTGAGCTCTTATTAGCTCTTATACAGCTCGTCCA | TTATTAGCTCTTATACAGCTCGTCCA |
| oWL00367_(tsPurple)_forward | 34 | 70.5 | 60.8 | 3.4 | AGAAAGAGTCCCATGGCGAGCTTAGTCAAGAAAG | ATGGCGAGCTTAGTCAAGAAAG |
| oWL00368_(tsPurple)_reverse | 43 | 69.0 | 61.7 | 4.3 | GGGGTTTTTTTTTGAGCTCTCATTAAGTTGCTTTCTCAGGCAC | TCATTAAGTTGCTTTCTCAGGCAC |
| oWL00369_(eforCP)_forward | 36 | 70.4 | 61.4 | 3.6 | AGAAAGAGTCCCATGGCAGTGATTAAGCAGGTAATG | ATGGCAGTGATTAAGCAGGTAATG |
| oWL00370_(eforCP)_reverse | 40 | 68.4 | 59.2 | 4 | GGGGTTTTTTTTTGAGCTCTTATTATGGGAGAGCCTTCGG | TTATTATGGGAGAGCCTTCGG |
| oWL00236_(pdar_2)_forward | 29 | 76.9 | 69.6 | 0 | CGACGACGATGGCCTGACTCCGCTGCACC | ATGGCCTGACTCCGCTGCACC |
| oWL00241_(backbone)_reverse | 35 | 71.9 | 65.1 | 3.5 | GCTCTGTCATCTTTGCCTTCGTTTATCTTGCCTGC | CTTTGCCTTCGTTTATCTTGCCTGC |
| oWL00242_(ARF5)_forward | 39 | 72.5 | 68.2 | 3.9 | GAAGGCAAAGATGACAGAGCAGAAAGCCCTAGTAAAGCG | ATGACAGAGCAGAAAGCCCTAGTAAAGCG |
| oWL00235_(IAA12-pdar)_(A_to_G)_reverse | 31 | 73.9 | 64.6 | 0 | CAGGCCATCGTCGTCGGTTGCGTTAACATCG | GTCGTCGGTTGCGTTAACATCG |
| oWL00194_(ARF7_0-10th)_forward | 35 | 69.8 | 59.4 | 3.5 | GCAGGCTTCAAAATGAAAGCTCCTTCATCAAATGG | ATGAAAGCTCCTTCATCAAATGG |
| oWL00193_(ARF7_0-10th)_reverse | 29 | 62.9 | 62.9 | 2.9 | AGGACATATGTAGAAAGGAGTTAAAACCG | AGGACATATGTAGAAAGGAGTTAAAACCG |
| oWL00192_(ARF7_51-end)_forward | 41 | 67.8 | 59.8 | 4.1 | CCTTTCTACATATGTCCTCCTTCGAGTACTATCTTTCCTGG | CCTTCGAGTACTATCTTTCCTGG |
| oWL00191_(ARF7_51-end)_reverse | 29 | 72.8 | 63.3 | 2.9 | GCTGGGTGTCACCGGTTAAACGAAGTGGC | TCACCGGTTAAACGAAGTGGC |
| oWL00190_(backbone)_forward | 33 | 72.7 | 64.3 | 3.3 | ACCGGTGACACCCAGCTTTCTTGTACAAAGTGG | CACCCAGCTTTCTTGTACAAAGTGG |
| oWL00189_(backbone)_reverse | 43 | 68.5 | 68.4 | 4.3 | GCTTTCATTTTGAAGCCTGCTTTTTTGTACAAACTTGTGATGG | TTTGAAGCCTGCTTTTTTGTACAAACTTGTGATGG |
| oWL00188_(ARF7_0-570th)_reverse | 24 | 60.2 | 60.2 | 2.4 | TCCATCTAAACCGTAAGAAGTTCC | TCCATCTAAACCGTAAGAAGTTCC |
| oWL00187_(ARF7_571-end)_forward | 35 | 69.614 | 56.869 | 3.5 | CTTACGGTTTAGATGGATCGAGGGGATATGACTCC | TCGAGGGGATATGACTCC |

**Table 1. List of primers designed by j5 for the assemblies.**
