## Supplementary material for "AssemblyTron: Flexible automation of DNA assembly with Opentrons OT-2 lab robots": File 9

```
Mon Sep 12, 2022 11:01 EDT
-ARF5-pdar-pmas00001.gb from 1 to 9308to_B10_IVA_rxn1_C5+206_TCYC1_Seq_R.ab1--  Matches:1277; Mismatches:5;
Gaps:8181; Unattempted:0_C10_IVA_rxn1_C6+206_TCYC1_Seq_R.ab1--  Matches:1130; Mismatches:2; Gaps:3635;
Unattempted:4687_A10_IVA_rxn1_C4+206_TCYC1_Seq_R.ab1--  Matches:1127; Mismatches:2; Gaps:3632;
Unattempted:4693_E03_IVA_rxn1_C3+206_TCYC1_Seq_R.ab1--  Matches:1106; Mismatches:3; Gaps:3641;
Unattempted:4708_C03_IVA_rxn1_C1+206_TCYC1_Seq_R.ab1--  Matches:1012; Mismatches:0; Gaps:3688;
Unattempted:4810_D03_IVA_rxn1_C2+206_TCYC1_Seq_R.ab1--  Matches:999; Mismatches:4; Gaps:3685; Unattempted:4823

                  *         *         *         *         *         *         *         *         *         *
     1>tcgcgcgtttcggtgatgacggtgaaaacctctgacacatgcagctcccggagacggtcacagcttgtctgtaagcggatgccgggagcagacaagcccg>100
  1438<~~~~~~~~~~~~~~~~~~~~~~~~~~~~~~~~~~~~~~~~~~~~~~~~~~~~~~~~~~~~~~~~~~~~~~~~~~~~~~~~~~~~~~~~~~~~~~~~<1438
  1468<~~~~~~~~~~~~~~~~~~~~~~~~~~~~~~~~~~~~~~~~~~~~~~~~~~~~~~~~~~~~~~~~~~~~~~~~~~~~~~~~~~~~~~~~~~~~~~~~<1468
  1464<~~~~~~~~~~~~~~~~~~~~~~~~~~~~~~~~~~~~~~~~~~~~~~~~~~~~~~~~~~~~~~~~~~~~~~~~~~~~~~~~~~~~~~~~~~~~~~~~<1464
  1407<~~~~~~~~~~~~~~~~~~~~~~~~~~~~~~~~~~~~~~~~~~~~~~~~~~~~~~~~~~~~~~~~~~~~~~~~~~~~~~~~~~~~~~~~~~~~~~~~<1407
  1379<~~~~~~~~~~~~~~~~~~~~~~~~~~~~~~~~~~~~~~~~~~~~~~~~~~~~~~~~~~~~~~~~~~~~~~~~~~~~~~~~~~~~~~~~~~~~~~~~<1379
  1410<~~~~~~~~~~~~~~~~~~~~~~~~~~~~~~~~~~~~~~~~~~~~~~~~~~~~~~~~~~~~~~~~~~~~~~~~~~~~~~~~~~~~~~~~~~~~~~~~<1410

                  *         *         *         *         *         *         *         *         *         *
   101>tcagggcgcgtcagcgggtgttggcgggtgtcggggctggcttaactatgcggcatcagagcagattgtactgagagtgcaccataattcgtttaaaccg>200
  1438<~~~~~~~~~~~~~~~~~~~~~~~~~~~~~~~~~~~~~~~~~~~~~~~~~~~~~~~~~~~~~~~~~~~~~~~~~~~~~~~~~~~~~~~~~~~~~~~~<1438
  1468<~~~~~~~~~~~~~~~~~~~~~~~~~~~~~~~~~~~~~~~~~~~~~~~~~~~~~~~~~~~~~~~~~~~~~~~~~~~~~~~~~~~~~~~~~~~~~~~~<1468
  1464<~~~~~~~~~~~~~~~~~~~~~~~~~~~~~~~~~~~~~~~~~~~~~~~~~~~~~~~~~~~~~~~~~~~~~~~~~~~~~~~~~~~~~~~~~~~~~~~~<1464
  1407<~~~~~~~~~~~~~~~~~~~~~~~~~~~~~~~~~~~~~~~~~~~~~~~~~~~~~~~~~~~~~~~~~~~~~~~~~~~~~~~~~~~~~~~~~~~~~~~~<1407
  1379<~~~~~~~~~~~~~~~~~~~~~~~~~~~~~~~~~~~~~~~~~~~~~~~~~~~~~~~~~~~~~~~~~~~~~~~~~~~~~~~~~~~~~~~~~~~~~~~~<1379
  1410<~~~~~~~~~~~~~~~~~~~~~~~~~~~~~~~~~~~~~~~~~~~~~~~~~~~~~~~~~~~~~~~~~~~~~~~~~~~~~~~~~~~~~~~~~~~~~~~~<1410

                  *         *         *         *         *         *         *         *         *         *
   201>ttttaagagcttggtgagcgctaggagtcactgccaggtatcgtttgaacacggcattagtcagggaagtcataacacagtcctttcccgcaattttctt>300
  1438<~~~~~~~~~~~~~~~~~~~~~~~~~~~~~~~~~~~~~~~~~~~~~~~~~~~~~~~~~~~~~~~~~~~~~~~~~~~~~~~~~~~~~~~~~~~~~~~~<1438
  1468<~~~~~~~~~~~~~~~~~~~~~~~~~~~~~~~~~~~~~~~~~~~~~~~~~~~~~~~~~~~~~~~~~~~~~~~~~~~~~~~~~~~~~~~~~~~~~~~~<1468
  1464<~~~~~~~~~~~~~~~~~~~~~~~~~~~~~~~~~~~~~~~~~~~~~~~~~~~~~~~~~~~~~~~~~~~~~~~~~~~~~~~~~~~~~~~~~~~~~~~~<1464
  1407<~~~~~~~~~~~~~~~~~~~~~~~~~~~~~~~~~~~~~~~~~~~~~~~~~~~~~~~~~~~~~~~~~~~~~~~~~~~~~~~~~~~~~~~~~~~~~~~~<1407
  1379<~~~~~~~~~~~~~~~~~~~~~~~~~~~~~~~~~~~~~~~~~~~~~~~~~~~~~~~~~~~~~~~~~~~~~~~~~~~~~~~~~~~~~~~~~~~~~~~~<1379
  1410<~~~~~~~~~~~~~~~~~~~~~~~~~~~~~~~~~~~~~~~~~~~~~~~~~~~~~~~~~~~~~~~~~~~~~~~~~~~~~~~~~~~~~~~~~~~~~~~~<1410

                  *         *         *         *         *         *         *         *         *         *
   301>tttctattactcttggcctcctctagtacactctatattttttttatgcctcggtaatgattttcatttttttttttccacctagcggatgactctttttt>400
  1438<~~~~~~~~~~~~~~~~~~~~~~~~~~~~~~~~~~~~~~~~~~~~~~~~~~~~~~~~~~~~~~~~~~~~~~~~~~~~~~~~~~~~~~~~~~~~~~~~<1438
  1468<~~~~~~~~~~~~~~~~~~~~~~~~~~~~~~~~~~~~~~~~~~~~~~~~~~~~~~~~~~~~~~~~~~~~~~~~~~~~~~~~~~~~~~~~~~~~~~~~<1468
  1464<~~~~~~~~~~~~~~~~~~~~~~~~~~~~~~~~~~~~~~~~~~~~~~~~~~~~~~~~~~~~~~~~~~~~~~~~~~~~~~~~~~~~~~~~~~~~~~~~<1464
```

```
1407<~~~~~~~~~~~~~~~~~~~~~~~~~~~~~~~~~~~~~~~~~~~~~~~~~~~~~~~~~~~~~~~~~~~~~~~~~~~~~~~~~~~~~~~~~~~~~~~~~<1407
1379<~~~~~~~~~~~~~~~~~~~~~~~~~~~~~~~~~~~~~~~~~~~~~~~~~~~~~~~~~~~~~~~~~~~~~~~~~~~~~~~~~~~~~~~~~~~~~~~~<1379
1410<~~~~~~~~~~~~~~~~~~~~~~~~~~~~~~~~~~~~~~~~~~~~~~~~~~~~~~~~~~~~~~~~~~~~~~~~~~~~~~~~~~~~~~~~~~~~~~~~<1410

              *         *         *         *         *         *         *         *         *         *
 401>tttcttagcgattggcattatcacataatgaattatacattatataaagtaatgtgatttcttcgaagaatatactaaaaaatgagcaggcaagataaac>500
1438<~~~~~~~~~~~~~~~~~~~~~~~~~~~~~~~~~~~~~~~~~~~~~~~~~~~~~~~~~~~~~~~~~~~~~~~~~~~~~~~~~~~~~~~~~~~~~~~~<1438
1468<~~~~~~~~~~~~~~~~~~~~~~~~~~~~~~~~~~~~~~~~~~~~~~~~~~~~~~~~~~~~~~~~~~~~~~~~~~~~~~~~~~~~~~~~~~~~~~~~<1468
1464<~~~~~~~~~~~~~~~~~~~~~~~~~~~~~~~~~~~~~~~~~~~~~~~~~~~~~~~~~~~~~~~~~~~~~~~~~~~~~~~~~~~~~~~~~~~~~~~~<1464
1407<~~~~~~~~~~~~~~~~~~~~~~~~~~~~~~~~~~~~~~~~~~~~~~~~~~~~~~~~~~~~~~~~~~~~~~~~~~~~~~~~~~~~~~~~~~~~~~~~<1407
1379<~~~~~~~~~~~~~~~~~~~~~~~~~~~~~~~~~~~~~~~~~~~~~~~~~~~~~~~~~~~~~~~~~~~~~~~~~~~~~~~~~~~~~~~~~~~~~~~~<1379
1410<~~~~~~~~~~~~~~~~~~~~~~~~~~~~~~~~~~~~~~~~~~~~~~~~~~~~~~~~~~~~~~~~~~~~~~~~~~~~~~~~~~~~~~~~~~~~~~~~<1410

              *         *         *         *         *         *         *         *         *         *
 501>gaaggcaaagatgacagagcagaaagccctagtaaagcgtattacaaatgaaaccaagattcagattgcgatctctttaaagggtggtcccctagcgata>600
1438<~~~~~~~~~~~~~~~~~~~~~~~~~~~~~~~~~~~~~~~~~~~~~~~~~~~~~~~~~~~~~~~~~~~~~~~~~~~~~~~~~~~~~~~~~~~~~~~~<1438
1468<~~~~~~~~~~~~~~~~~~~~~~~~~~~~~~~~~~~~~~~~~~~~~~~~~~~~~~~~~~~~~~~~~~~~~~~~~~~~~~~~~~~~~~~~~~~~~~~~<1468
1464<~~~~~~~~~~~~~~~~~~~~~~~~~~~~~~~~~~~~~~~~~~~~~~~~~~~~~~~~~~~~~~~~~~~~~~~~~~~~~~~~~~~~~~~~~~~~~~~~<1464
1407<~~~~~~~~~~~~~~~~~~~~~~~~~~~~~~~~~~~~~~~~~~~~~~~~~~~~~~~~~~~~~~~~~~~~~~~~~~~~~~~~~~~~~~~~~~~~~~~~<1407
1379<~~~~~~~~~~~~~~~~~~~~~~~~~~~~~~~~~~~~~~~~~~~~~~~~~~~~~~~~~~~~~~~~~~~~~~~~~~~~~~~~~~~~~~~~~~~~~~~~<1379
1410<~~~~~~~~~~~~~~~~~~~~~~~~~~~~~~~~~~~~~~~~~~~~~~~~~~~~~~~~~~~~~~~~~~~~~~~~~~~~~~~~~~~~~~~~~~~~~~~~<1410

              *         *         *         *         *         *         *         *         *         *
 601>gagcactcgatcttcccagaaaaagaggcagaagcagtagcagaacaggccacacaatcgcaagtgattaacgtccacacaggtatagggtttctggacc>700
1438<~~~~~~~~~~~~~~~~~~~~~~~~~~~~~~~~~~~~~~~~~~~~~~~~~~~~~~~~~~~~~~~~~~~~~~~~~~~~~~~~~~~~~~~~~~~~~~~~<1438
1468<~~~~~~~~~~~~~~~~~~~~~~~~~~~~~~~~~~~~~~~~~~~~~~~~~~~~~~~~~~~~~~~~~~~~~~~~~~~~~~~~~~~~~~~~~~~~~~~~<1468
1464<~~~~~~~~~~~~~~~~~~~~~~~~~~~~~~~~~~~~~~~~~~~~~~~~~~~~~~~~~~~~~~~~~~~~~~~~~~~~~~~~~~~~~~~~~~~~~~~~<1464
1407<~~~~~~~~~~~~~~~~~~~~~~~~~~~~~~~~~~~~~~~~~~~~~~~~~~~~~~~~~~~~~~~~~~~~~~~~~~~~~~~~~~~~~~~~~~~~~~~~<1407
1379<~~~~~~~~~~~~~~~~~~~~~~~~~~~~~~~~~~~~~~~~~~~~~~~~~~~~~~~~~~~~~~~~~~~~~~~~~~~~~~~~~~~~~~~~~~~~~~~~<1379
1410<~~~~~~~~~~~~~~~~~~~~~~~~~~~~~~~~~~~~~~~~~~~~~~~~~~~~~~~~~~~~~~~~~~~~~~~~~~~~~~~~~~~~~~~~~~~~~~~~<1410

              *         *         *         *         *         *         *         *         *         *
 701>atatgatacatgctctggccaagcattccggctggtcgctaatcgttgagtgcattggtgacttacacatagacgaccatcacaccactgaagactgcgg>800
1438<~~~~~~~~~~~~~~~~~~~~~~~~~~~~~~~~~~~~~~~~~~~~~~~~~~~~~~~~~~~~~~~~~~~~~~~~~~~~~~~~~~~~~~~~~~~~~~~~<1438
1468<~~~~~~~~~~~~~~~~~~~~~~~~~~~~~~~~~~~~~~~~~~~~~~~~~~~~~~~~~~~~~~~~~~~~~~~~~~~~~~~~~~~~~~~~~~~~~~~~<1468
1464<~~~~~~~~~~~~~~~~~~~~~~~~~~~~~~~~~~~~~~~~~~~~~~~~~~~~~~~~~~~~~~~~~~~~~~~~~~~~~~~~~~~~~~~~~~~~~~~~<1464
1407<~~~~~~~~~~~~~~~~~~~~~~~~~~~~~~~~~~~~~~~~~~~~~~~~~~~~~~~~~~~~~~~~~~~~~~~~~~~~~~~~~~~~~~~~~~~~~~~~<1407
1379<~~~~~~~~~~~~~~~~~~~~~~~~~~~~~~~~~~~~~~~~~~~~~~~~~~~~~~~~~~~~~~~~~~~~~~~~~~~~~~~~~~~~~~~~~~~~~~~~<1379
1410<~~~~~~~~~~~~~~~~~~~~~~~~~~~~~~~~~~~~~~~~~~~~~~~~~~~~~~~~~~~~~~~~~~~~~~~~~~~~~~~~~~~~~~~~~~~~~~~~<1410
```

```
              *         *         *         *         *         *         *         *         *         *
 801>gattgctctcggtcaagcttttaaagaggccctaggggccgtgcgtggagtaaaaaggtttggatcaggatttgcgcctttggatgaggcactttccaga>900
1438<~~~~~~~~~~~~~~~~~~~~~~~~~~~~~~~~~~~~~~~~~~~~~~~~~~~~~~~~~~~~~~~~~~~~~~~~~~~~~~~~~~~~~~~~~~~~~~~~~~~<1438
1468<~~~~~~~~~~~~~~~~~~~~~~~~~~~~~~~~~~~~~~~~~~~~~~~~~~~~~~~~~~~~~~~~~~~~~~~~~~~~~~~~~~~~~~~~~~~~~~~~~~~<1468
1464<~~~~~~~~~~~~~~~~~~~~~~~~~~~~~~~~~~~~~~~~~~~~~~~~~~~~~~~~~~~~~~~~~~~~~~~~~~~~~~~~~~~~~~~~~~~~~~~~~~~<1464
1407<~~~~~~~~~~~~~~~~~~~~~~~~~~~~~~~~~~~~~~~~~~~~~~~~~~~~~~~~~~~~~~~~~~~~~~~~~~~~~~~~~~~~~~~~~~~~~~~~~~~<1407
1379<~~~~~~~~~~~~~~~~~~~~~~~~~~~~~~~~~~~~~~~~~~~~~~~~~~~~~~~~~~~~~~~~~~~~~~~~~~~~~~~~~~~~~~~~~~~~~~~~~~~<1379
1410<~~~~~~~~~~~~~~~~~~~~~~~~~~~~~~~~~~~~~~~~~~~~~~~~~~~~~~~~~~~~~~~~~~~~~~~~~~~~~~~~~~~~~~~~~~~~~~~~~~~<1410

              *         *         *         *         *         *         *         *         *         *
 901>gcggtggtagatctttcgaacaggccgtacgcagttgtcgaacttggtttgcaaagggagaaagtaggagatctctcttgcgagatgatcccgcattttc>1000
1438<~~~~~~~~~~~~~~~~~~~~~~~~~~~~~~~~~~~~~~~~~~~~~~~~~~~~~~~~~~~~~~~~~~~~~~~~~~~~~~~~~~~~~~~~~~~~~~~~~~~<1438
1468<~~~~~~~~~~~~~~~~~~~~~~~~~~~~~~~~~~~~~~~~~~~~~~~~~~~~~~~~~~~~~~~~~~~~~~~~~~~~~~~~~~~~~~~~~~~~~~~~~~~<1468
1464<~~~~~~~~~~~~~~~~~~~~~~~~~~~~~~~~~~~~~~~~~~~~~~~~~~~~~~~~~~~~~~~~~~~~~~~~~~~~~~~~~~~~~~~~~~~~~~~~~~~<1464
1407<~~~~~~~~~~~~~~~~~~~~~~~~~~~~~~~~~~~~~~~~~~~~~~~~~~~~~~~~~~~~~~~~~~~~~~~~~~~~~~~~~~~~~~~~~~~~~~~~~~~<1407
1379<~~~~~~~~~~~~~~~~~~~~~~~~~~~~~~~~~~~~~~~~~~~~~~~~~~~~~~~~~~~~~~~~~~~~~~~~~~~~~~~~~~~~~~~~~~~~~~~~~~~<1379
1410<~~~~~~~~~~~~~~~~~~~~~~~~~~~~~~~~~~~~~~~~~~~~~~~~~~~~~~~~~~~~~~~~~~~~~~~~~~~~~~~~~~~~~~~~~~~~~~~~~~~<1410

              *         *         *         *         *         *         *         *         *         *
1001>ttgaaagctttgcagaggctagcagaattaccctccacgttgattgtctgcgaggcaagaatgatcatcaccgtagtgagagtgcgttcaaggctcttgc>1100
1438<~~~~~~~~~~~~~~~~~~~~~~~~~~~~~~~~~~~~~~~~~~~~~~~~~~~~~~~~~~~~~~~~~~~~~~~~~~~~~~~~~~~~~~~~~~~~~~~~~~~<1438
1468<~~~~~~~~~~~~~~~~~~~~~~~~~~~~~~~~~~~~~~~~~~~~~~~~~~~~~~~~~~~~~~~~~~~~~~~~~~~~~~~~~~~~~~~~~~~~~~~~~~~<1468
1464<~~~~~~~~~~~~~~~~~~~~~~~~~~~~~~~~~~~~~~~~~~~~~~~~~~~~~~~~~~~~~~~~~~~~~~~~~~~~~~~~~~~~~~~~~~~~~~~~~~~<1464
1407<~~~~~~~~~~~~~~~~~~~~~~~~~~~~~~~~~~~~~~~~~~~~~~~~~~~~~~~~~~~~~~~~~~~~~~~~~~~~~~~~~~~~~~~~~~~~~~~~~~~<1407
1379<~~~~~~~~~~~~~~~~~~~~~~~~~~~~~~~~~~~~~~~~~~~~~~~~~~~~~~~~~~~~~~~~~~~~~~~~~~~~~~~~~~~~~~~~~~~~~~~~~~~<1379
1410<~~~~~~~~~~~~~~~~~~~~~~~~~~~~~~~~~~~~~~~~~~~~~~~~~~~~~~~~~~~~~~~~~~~~~~~~~~~~~~~~~~~~~~~~~~~~~~~~~~~<1410

              *         *         *         *         *         *         *         *         *         *
1101>ggttgccataagagaagccacctcgcccaatggtaccaacgatgttccctccaccaaaggtgttcttatgtaggcgaatttcttatgatttatgattttt>1200
1438<~~~~~~~~~~~~~~~~~~~~~~~~~~~~~~~~~~~~~~~~~~~~~~~~~~~~~~~~~~~~~~~~~~~~~~~~~~~~~~~~~~~~~~~~~~~~~~~~~~~<1438
1468<~~~~~~~~~~~~~~~~~~~~~~~~~~~~~~~~~~~~~~~~~~~~~~~~~~~~~~~~~~~~~~~~~~~~~~~~~~~~~~~~~~~~~~~~~~~~~~~~~~~<1468
1464<~~~~~~~~~~~~~~~~~~~~~~~~~~~~~~~~~~~~~~~~~~~~~~~~~~~~~~~~~~~~~~~~~~~~~~~~~~~~~~~~~~~~~~~~~~~~~~~~~~~<1464
1407<~~~~~~~~~~~~~~~~~~~~~~~~~~~~~~~~~~~~~~~~~~~~~~~~~~~~~~~~~~~~~~~~~~~~~~~~~~~~~~~~~~~~~~~~~~~~~~~~~~~<1407
1379<~~~~~~~~~~~~~~~~~~~~~~~~~~~~~~~~~~~~~~~~~~~~~~~~~~~~~~~~~~~~~~~~~~~~~~~~~~~~~~~~~~~~~~~~~~~~~~~~~~~<1379
1410<~~~~~~~~~~~~~~~~~~~~~~~~~~~~~~~~~~~~~~~~~~~~~~~~~~~~~~~~~~~~~~~~~~~~~~~~~~~~~~~~~~~~~~~~~~~~~~~~~~~<1410

              *         *         *         *         *         *         *         *         *         *
```

1201>attattaaataagttataaaaaaaataagtgtatacaaattttaaagtgactcttaggttttaaaacgaaaattcttattcttgagtaactctttcctgt>1300
1438<~~~~~~~~~~~~~~~~~~~~~~~~~~~~~~~~~~~~~~~~~~~~~~~~~~~~~~~~~~~~~~~~~~~~~~~~~~~~~~~~~~~~~~~~~~~~~~~~<1438
1468<~~~~~~~~~~~~~~~~~~~~~~~~~~~~~~~~~~~~~~~~~~~~~~~~~~~~~~~~~~~~~~~~~~~~~~~~~~~~~~~~~~~~~~~~~~~~~~~~<1468
1464<~~~~~~~~~~~~~~~~~~~~~~~~~~~~~~~~~~~~~~~~~~~~~~~~~~~~~~~~~~~~~~~~~~~~~~~~~~~~~~~~~~~~~~~~~~~~~~~~<1464
1407<~~~~~~~~~~~~~~~~~~~~~~~~~~~~~~~~~~~~~~~~~~~~~~~~~~~~~~~~~~~~~~~~~~~~~~~~~~~~~~~~~~~~~~~~~~~~~~~~<1407
1379<~~~~~~~~~~~~~~~~~~~~~~~~~~~~~~~~~~~~~~~~~~~~~~~~~~~~~~~~~~~~~~~~~~~~~~~~~~~~~~~~~~~~~~~~~~~~~~~~<1379
1410<~~~~~~~~~~~~~~~~~~~~~~~~~~~~~~~~~~~~~~~~~~~~~~~~~~~~~~~~~~~~~~~~~~~~~~~~~~~~~~~~~~~~~~~~~~~~~~~~<1410

* * * * * * * * * *

1301>aggtcaggttgctttctcaggtatagcatgaggtcgctcttattgaccacacctcaagaaatgatggtaaatgaaataggaaatcaaggagcatgaaggc>1400
1438<~~~~~~~~~~~~~~~~~~~~~~~~~~~~~~~~~~~~~~~~~~~~~~~~~~~~~~~~~~~~~~~~~~~~~~~~~~~~~~~~~~~~~~~~~~~~~~~~<1438
1468<~~~~~~~~~~~~~~~~~~~~~~~~~~~~~~~~~~~~~~~~~~~~~~~~~~~~~~~~~~~~~~~~~~~~~~~~~~~~~~~~~~~~~~~~~~~~~~~~<1468
1464<~~~~~~~~~~~~~~~~~~~~~~~~~~~~~~~~~~~~~~~~~~~~~~~~~~~~~~~~~~~~~~~~~~~~~~~~~~~~~~~~~~~~~~~~~~~~~~~~<1464
1407<~~~~~~~~~~~~~~~~~~~~~~~~~~~~~~~~~~~~~~~~~~~~~~~~~~~~~~~~~~~~~~~~~~~~~~~~~~~~~~~~~~~~~~~~~~~~~~~~<1407
1379<~~~~~~~~~~~~~~~~~~~~~~~~~~~~~~~~~~~~~~~~~~~~~~~~~~~~~~~~~~~~~~~~~~~~~~~~~~~~~~~~~~~~~~~~~~~~~~~~<1379
1410<~~~~~~~~~~~~~~~~~~~~~~~~~~~~~~~~~~~~~~~~~~~~~~~~~~~~~~~~~~~~~~~~~~~~~~~~~~~~~~~~~~~~~~~~~~~~~~~~<1410

* * * * * * * * * *

1401>aaaagacaaatataagggtcgaacgaaaaataaagtgaaaagtgttgatatgatgtatttggctttgcggcgccgaaaaaacgagtttacgcaattgcac>1500
1438<~~~~~~~~~~~~~~~~~~~~~~~~~~~~~~~~~~~~~~~~~~~~~~~~~~~~~~~~~~~~~~~~~~~~~~~~~~~~~~~~~~~~~~~~~~~~~~~~<1438
1468<~~~~~~~~~~~~~~~~~~~~~~~~~~~~~~~~~~~~~~~~~~~~~~~~~~~~~~~~~~~~~~~~~~~~~~~~~~~~~~~~~~~~~~~~~~~~~~~~<1468
1464<~~~~~~~~~~~~~~~~~~~~~~~~~~~~~~~~~~~~~~~~~~~~~~~~~~~~~~~~~~~~~~~~~~~~~~~~~~~~~~~~~~~~~~~~~~~~~~~~<1464
1407<~~~~~~~~~~~~~~~~~~~~~~~~~~~~~~~~~~~~~~~~~~~~~~~~~~~~~~~~~~~~~~~~~~~~~~~~~~~~~~~~~~~~~~~~~~~~~~~~<1407
1379<~~~~~~~~~~~~~~~~~~~~~~~~~~~~~~~~~~~~~~~~~~~~~~~~~~~~~~~~~~~~~~~~~~~~~~~~~~~~~~~~~~~~~~~~~~~~~~~~<1379
1410<~~~~~~~~~~~~~~~~~~~~~~~~~~~~~~~~~~~~~~~~~~~~~~~~~~~~~~~~~~~~~~~~~~~~~~~~~~~~~~~~~~~~~~~~~~~~~~~~<1410

* * * * * * * * * *

1501>aatcatgctgactctgtggcggacccgcgctcttgccggcccggcgataacgctgggcgtgaggctgtgcccggcggagttttttgcgcctgcattttcc>1600
1438<~~~~~~~~~~~~~~~~~~~~~~~~~~~~~~~~~~~~~~~~~~~~~~~~~~~~~~~~~~~~~~~~~~~~~~~~~~~~~~~~~~~~~~~~~~~~~~~~<1438
1468<~~~~~~~~~~~~~~~~~~~~~~~~~~~~~~~~~~~~~~~~~~~~~~~~~~~~~~~~~~~~~~~~~~~~~~~~~~~~~~~~~~~~~~~~~~~~~~~~<1468
1464<~~~~~~~~~~~~~~~~~~~~~~~~~~~~~~~~~~~~~~~~~~~~~~~~~~~~~~~~~~~~~~~~~~~~~~~~~~~~~~~~~~~~~~~~~~~~~~~~<1464
1407<~~~~~~~~~~~~~~~~~~~~~~~~~~~~~~~~~~~~~~~~~~~~~~~~~~~~~~~~~~~~~~~~~~~~~~~~~~~~~~~~~~~~~~~~~~~~~~~~<1407
1379<~~~~~~~~~~~~~~~~~~~~~~~~~~~~~~~~~~~~~~~~~~~~~~~~~~~~~~~~~~~~~~~~~~~~~~~~~~~~~~~~~~~~~~~~~~~~~~~~<1379
1410<~~~~~~~~~~~~~~~~~~~~~~~~~~~~~~~~~~~~~~~~~~~~~~~~~~~~~~~~~~~~~~~~~~~~~~~~~~~~~~~~~~~~~~~~~~~~~~~~<1410

* * * * * * * * * *

1601>aaggtttaccctgcgctaaggggcgagattggagaagcaataagaatgccggttggggttgcgatgatgacgaccacgacaactggtgtcattatttaag>1700
1438<~~~~~~~~~~~~~~~~~~~~~~~~~~~~~~~~~~~~~~~~~~~~~~~~~~~~~~~~~~~~~~~~~~~~~~~~~~~~~~~~~~~~~~~~~~~~~~~~<1438
1468<~~~~~~~~~~~~~~~~~~~~~~~~~~~~~~~~~~~~~~~~~~~~~~~~~~~~~~~~~~~~~~~~~~~~~~~~~~~~~~~~~~~~~~~~~~~~~~~~<1468

```
1464<~~~~~~~~~~~~~~~~~~~~~~~~~~~~~~~~~~~~~~~~~~~~~~~~~~~~~~~~~~~~~~~~~~~~~~~~~~~~~~~~~~~~~~~~~~~~~~~~~<1464
1407<~~~~~~~~~~~~~~~~~~~~~~~~~~~~~~~~~~~~~~~~~~~~~~~~~~~~~~~~~~~~~~~~~~~~~~~~~~~~~~~~~~~~~~~~~~~~~~~~<1407
1379<~~~~~~~~~~~~~~~~~~~~~~~~~~~~~~~~~~~~~~~~~~~~~~~~~~~~~~~~~~~~~~~~~~~~~~~~~~~~~~~~~~~~~~~~~~~~~~~~<1379
1410<~~~~~~~~~~~~~~~~~~~~~~~~~~~~~~~~~~~~~~~~~~~~~~~~~~~~~~~~~~~~~~~~~~~~~~~~~~~~~~~~~~~~~~~~~~~~~~~~<1410

              *         *         *         *         *         *         *         *         *         *
1701>ttgccgaaagaacctgagtgcatttgcaacatgagtatactagaagaatgagccaagacttgcgagacgcgagtttgccggtggtgcgaacaatagagcg>1800
1438<~~~~~~~~~~~~~~~~~~~~~~~~~~~~~~~~~~~~~~~~~~~~~~~~~~~~~~~~~~~~~~~~~~~~~~~~~~~~~~~~~~~~~~~~~~~~~~~~<1438
1468<~~~~~~~~~~~~~~~~~~~~~~~~~~~~~~~~~~~~~~~~~~~~~~~~~~~~~~~~~~~~~~~~~~~~~~~~~~~~~~~~~~~~~~~~~~~~~~~~<1468
1464<~~~~~~~~~~~~~~~~~~~~~~~~~~~~~~~~~~~~~~~~~~~~~~~~~~~~~~~~~~~~~~~~~~~~~~~~~~~~~~~~~~~~~~~~~~~~~~~~<1464
1407<~~~~~~~~~~~~~~~~~~~~~~~~~~~~~~~~~~~~~~~~~~~~~~~~~~~~~~~~~~~~~~~~~~~~~~~~~~~~~~~~~~~~~~~~~~~~~~~~<1407
1379<~~~~~~~~~~~~~~~~~~~~~~~~~~~~~~~~~~~~~~~~~~~~~~~~~~~~~~~~~~~~~~~~~~~~~~~~~~~~~~~~~~~~~~~~~~~~~~~~<1379
1410<~~~~~~~~~~~~~~~~~~~~~~~~~~~~~~~~~~~~~~~~~~~~~~~~~~~~~~~~~~~~~~~~~~~~~~~~~~~~~~~~~~~~~~~~~~~~~~~~<1410

              *         *         *         *         *         *         *         *         *         *
1801>accatgaccttgaaggtgagacgcgcataaccgctagagtactttgaagaggaaacagcaatagggttgctaccagtataaatagacaggtacatacaac>1900
1438<~~~~~~~~~~~~~~~~~~~~~~~~~~~~~~~~~~~~~~~~~~~~~~~~~~~~~~~~~~~~~~~~~~~~~~~~~~~~~~~~~~~~~~~~~~~~~~~~<1438
1468<~~~~~~~~~~~~~~~~~~~~~~~~~~~~~~~~~~~~~~~~~~~~~~~~~~~~~~~~~~~~~~~~~~~~~~~~~~~~~~~~~~~~~~~~~~~~~~~~<1468
1464<~~~~~~~~~~~~~~~~~~~~~~~~~~~~~~~~~~~~~~~~~~~~~~~~~~~~~~~~~~~~~~~~~~~~~~~~~~~~~~~~~~~~~~~~~~~~~~~~<1464
1407<~~~~~~~~~~~~~~~~~~~~~~~~~~~~~~~~~~~~~~~~~~~~~~~~~~~~~~~~~~~~~~~~~~~~~~~~~~~~~~~~~~~~~~~~~~~~~~~~<1407
1379<~~~~~~~~~~~~~~~~~~~~~~~~~~~~~~~~~~~~~~~~~~~~~~~~~~~~~~~~~~~~~~~~~~~~~~~~~~~~~~~~~~~~~~~~~~~~~~~~<1379
1410<~~~~~~~~~~~~~~~~~~~~~~~~~~~~~~~~~~~~~~~~~~~~~~~~~~~~~~~~~~~~~~~~~~~~~~~~~~~~~~~~~~~~~~~~~~~~~~~~<1410

              *         *         *         *         *         *         *         *         *         *
1901>actggaaatggttgtctgtttgagtacgctttcaattcatttgggtgtgcactttattatgttacaatatggaagggaactttacacttctcctatgcac>2000
1438<~~~~~~~~~~~~~~~~~~~~~~~~~~~~~~~~~~~~~~~~~~~~~~~~~~~~~~~~~~~~~~~~~~~~~~~~~~~~~~~~~~~~~~~~~~~~~~~~<1438
1468<~~~~~~~~~~~~~~~~~~~~~~~~~~~~~~~~~~~~~~~~~~~~~~~~~~~~~~~~~~~~~~~~~~~~~~~~~~~~~~~~~~~~~~~~~~~~~~~~<1468
1464<~~~~~~~~~~~~~~~~~~~~~~~~~~~~~~~~~~~~~~~~~~~~~~~~~~~~~~~~~~~~~~~~~~~~~~~~~~~~~~~~~~~~~~~~~~~~~~~~<1464
1407<~~~~~~~~~~~~~~~~~~~~~~~~~~~~~~~~~~~~~~~~~~~~~~~~~~~~~~~~~~~~~~~~~~~~~~~~~~~~~~~~~~~~~~~~~~~~~~~~<1407
1379<~~~~~~~~~~~~~~~~~~~~~~~~~~~~~~~~~~~~~~~~~~~~~~~~~~~~~~~~~~~~~~~~~~~~~~~~~~~~~~~~~~~~~~~~~~~~~~~~<1379
1410<~~~~~~~~~~~~~~~~~~~~~~~~~~~~~~~~~~~~~~~~~~~~~~~~~~~~~~~~~~~~~~~~~~~~~~~~~~~~~~~~~~~~~~~~~~~~~~~~<1410

              *         *         *         *         *         *         *         *         *         *
2001>atatattaattaaagtccaatgctagtagagaagggggggtaacacccctccgcgctcttttccgatttttttctaaaccgtggaatatttcggatatcct>2100
1438<~~~~~~~~~~~~~~~~~~~~~~~~~~~~~~~~~~~~~~~~~~~~~~~~~~~~~~~~~~~~~~~~~~~~~~~~~~~~~~~~~~~~~~~~~~~~~~~~<1438
1468<~~~~~~~~~~~~~~~~~~~~~~~~~~~~~~~~~~~~~~~~~~~~~~~~~~~~~~~~~~~~~~~~~~~~~~~~~~~~~~~~~~~~~~~~~~~~~~~~<1468
1464<~~~~~~~~~~~~~~~~~~~~~~~~~~~~~~~~~~~~~~~~~~~~~~~~~~~~~~~~~~~~~~~~~~~~~~~~~~~~~~~~~~~~~~~~~~~~~~~~<1464
1407<~~~~~~~~~~~~~~~~~~~~~~~~~~~~~~~~~~~~~~~~~~~~~~~~~~~~~~~~~~~~~~~~~~~~~~~~~~~~~~~~~~~~~~~~~~~~~~~~<1407
1379<~~~~~~~~~~~~~~~~~~~~~~~~~~~~~~~~~~~~~~~~~~~~~~~~~~~~~~~~~~~~~~~~~~~~~~~~~~~~~~~~~~~~~~~~~~~~~~~~<1379
```

```
1410<~~~~~~~~~~~~~~~~~~~~~~~~~~~~~~~~~~~~~~~~~~~~~~~~~~~~~~~~~~~~~~~~~~~~~~~~~~~~~~~~~~~~~~~~~~~~~~~~<1410

              *         *         *         *         *         *         *         *         *         *
2101>tttgttgtttccgggtgtacaatatggacttcctcttttctggcaaccaaacccatacatcgggattcctataataccttcgttggtctccctaacatgt>2200
1438<~~~~~~~~~~~~~~~~~~~~~~~~~~~~~~~~~~~~~~~~~~~~~~~~~~~~~~~~~~~~~~~~~~~~~~~~~~~~~~~~~~~~~~~~~~~~~~~~<1438
1468<~~~~~~~~~~~~~~~~~~~~~~~~~~~~~~~~~~~~~~~~~~~~~~~~~~~~~~~~~~~~~~~~~~~~~~~~~~~~~~~~~~~~~~~~~~~~~~~~<1468
1464<~~~~~~~~~~~~~~~~~~~~~~~~~~~~~~~~~~~~~~~~~~~~~~~~~~~~~~~~~~~~~~~~~~~~~~~~~~~~~~~~~~~~~~~~~~~~~~~~<1464
1407<~~~~~~~~~~~~~~~~~~~~~~~~~~~~~~~~~~~~~~~~~~~~~~~~~~~~~~~~~~~~~~~~~~~~~~~~~~~~~~~~~~~~~~~~~~~~~~~~<1407
1379<~~~~~~~~~~~~~~~~~~~~~~~~~~~~~~~~~~~~~~~~~~~~~~~~~~~~~~~~~~~~~~~~~~~~~~~~~~~~~~~~~~~~~~~~~~~~~~~~<1379
1410<~~~~~~~~~~~~~~~~~~~~~~~~~~~~~~~~~~~~~~~~~~~~~~~~~~~~~~~~~~~~~~~~~~~~~~~~~~~~~~~~~~~~~~~~~~~~~~~~<1410

              *         *         *         *         *         *         *         *         *         *
2201>aggtggcggaggggagatatacaatagaacagataccagacaagacataatgggctaaacaagactacaccaattacactgcctcattgatggtggtaca>2300
1438<~~~~~~~~~~~~~~~~~~~~~~~~~~~~~~~~~~~~~~~~~~~~~~~~~~~~~~~~~~~~~~~~~~~~~~~~~~~~~~~~~~~~~~~~~~~~~~~~<1438
1468<~~~~~~~~~~~~~~~~~~~~~~~~~~~~~~~~~~~~~~~~~~~~~~~~~~~~~~~~~~~~~~~~~~~~~~~~~~~~~~~~~~~~~~~~~~~~~~~~<1468
1464<~~~~~~~~~~~~~~~~~~~~~~~~~~~~~~~~~~~~~~~~~~~~~~~~~~~~~~~~~~~~~~~~~~~~~~~~~~~~~~~~~~~~~~~~~~~~~~~~<1464
1407<~~~~~~~~~~~~~~~~~~~~~~~~~~~~~~~~~~~~~~~~~~~~~~~~~~~~~~~~~~~~~~~~~~~~~~~~~~~~~~~~~~~~~~~~~~~~~~~~<1407
1379<~~~~~~~~~~~~~~~~~~~~~~~~~~~~~~~~~~~~~~~~~~~~~~~~~~~~~~~~~~~~~~~~~~~~~~~~~~~~~~~~~~~~~~~~~~~~~~~~<1379
1410<~~~~~~~~~~~~~~~~~~~~~~~~~~~~~~~~~~~~~~~~~~~~~~~~~~~~~~~~~~~~~~~~~~~~~~~~~~~~~~~~~~~~~~~~~~~~~~~~<1410

              *         *         *         *         *         *         *         *         *         *
2301>taacgaactaatactgtagccctagacttgatagccatcatcatatcgaagtttcactacccttttttccatttgccatctattgaagtaataataggcgc>2400
1438<~~~~~~~~~~~~~~~~~~~~~~~~~~~~~~~~~~~~~~~~~~~~~~~~~~~~~~~~~~~~~~~~~~~~~~~~~~~~~~~~~~~~~~~~~~~~~~~~<1438
1468<~~~~~~~~~~~~~~~~~~~~~~~~~~~~~~~~~~~~~~~~~~~~~~~~~~~~~~~~~~~~~~~~~~~~~~~~~~~~~~~~~~~~~~~~~~~~~~~~<1468
1464<~~~~~~~~~~~~~~~~~~~~~~~~~~~~~~~~~~~~~~~~~~~~~~~~~~~~~~~~~~~~~~~~~~~~~~~~~~~~~~~~~~~~~~~~~~~~~~~~<1464
1407<~~~~~~~~~~~~~~~~~~~~~~~~~~~~~~~~~~~~~~~~~~~~~~~~~~~~~~~~~~~~~~~~~~~~~~~~~~~~~~~~~~~~~~~~~~~~~~~~<1407
1379<~~~~~~~~~~~~~~~~~~~~~~~~~~~~~~~~~~~~~~~~~~~~~~~~~~~~~~~~~~~~~~~~~~~~~~~~~~~~~~~~~~~~~~~~~~~~~~~~<1379
1410<~~~~~~~~~~~~~~~~~~~~~~~~~~~~~~~~~~~~~~~~~~~~~~~~~~~~~~~~~~~~~~~~~~~~~~~~~~~~~~~~~~~~~~~~~~~~~~~~<1410

              *         *         *         *         *         *         *         *         *         *
2401>atgcaacttcttttcttttttttttcttttctctctcccccgttgttgtctcaccatatccgcaatgacaaaaaaatgatggaagacactaaaggaaaaaa>2500
1438<~~~~~~~~~~~~~~~~~~~~~~~~~~~~~~~~~~~~~~~~~~~~~~~~~~~~~~~~~~~~~~~~~~~~~~~~~~~~~~~~~~~~~~~~~~~~~~~~<1438
1468<~~~~~~~~~~~~~~~~~~~~~~~~~~~~~~~~~~~~~~~~~~~~~~~~~~~~~~~~~~~~~~~~~~~~~~~~~~~~~~~~~~~~~~~~~~~~~~~~<1468
1464<~~~~~~~~~~~~~~~~~~~~~~~~~~~~~~~~~~~~~~~~~~~~~~~~~~~~~~~~~~~~~~~~~~~~~~~~~~~~~~~~~~~~~~~~~~~~~~~~<1464
1407<~~~~~~~~~~~~~~~~~~~~~~~~~~~~~~~~~~~~~~~~~~~~~~~~~~~~~~~~~~~~~~~~~~~~~~~~~~~~~~~~~~~~~~~~~~~~~~~~<1407
1379<~~~~~~~~~~~~~~~~~~~~~~~~~~~~~~~~~~~~~~~~~~~~~~~~~~~~~~~~~~~~~~~~~~~~~~~~~~~~~~~~~~~~~~~~~~~~~~~~<1379
1410<~~~~~~~~~~~~~~~~~~~~~~~~~~~~~~~~~~~~~~~~~~~~~~~~~~~~~~~~~~~~~~~~~~~~~~~~~~~~~~~~~~~~~~~~~~~~~~~~<1410
```

```
              *         *         *         *         *         *         *         *         *         *
2501>ttaacgacaaagacagcaccaacagatgtcgttgttccagagctgatgaggggtatctcgaagcacacgaaactttttccttccttcattcacgcacact>2600
1438<~~~~~~~~~~~~~~~~~~~~~~~~~~~~~~~~~~~~~~~~~~~~~~~~~~~~~~~~~~~~~~~~~~~~~~~~~~~~~~~~~~~~~~~~~~~~~~~~~~~~<1438
1468<~~~~~~~~~~~~~~~~~~~~~~~~~~~~~~~~~~~~~~~~~~~~~~~~~~~~~~~~~~~~~~~~~~~~~~~~~~~~~~~~~~~~~~~~~~~~~~~~~~~~<1468
1464<~~~~~~~~~~~~~~~~~~~~~~~~~~~~~~~~~~~~~~~~~~~~~~~~~~~~~~~~~~~~~~~~~~~~~~~~~~~~~~~~~~~~~~~~~~~~~~~~~~~~<1464
1407<~~~~~~~~~~~~~~~~~~~~~~~~~~~~~~~~~~~~~~~~~~~~~~~~~~~~~~~~~~~~~~~~~~~~~~~~~~~~~~~~~~~~~~~~~~~~~~~~~~~~<1407
1379<~~~~~~~~~~~~~~~~~~~~~~~~~~~~~~~~~~~~~~~~~~~~~~~~~~~~~~~~~~~~~~~~~~~~~~~~~~~~~~~~~~~~~~~~~~~~~~~~~~~~<1379
1410<~~~~~~~~~~~~~~~~~~~~~~~~~~~~~~~~~~~~~~~~~~~~~~~~~~~~~~~~~~~~~~~~~~~~~~~~~~~~~~~~~~~~~~~~~~~~~~~~~~~~<1410

              *         *         *         *         *         *         *         *         *         *
2601>actctctaatgagcaacggtatacggccttccttccagttacttgaatttgaaataaaaaaaagtttgctgtcttgctatcaagtataaatagacctgca>2700
1438<~~~~~~~~~~~~~~~~~~~~~~~~~~~~~~~~~~~~~~~~~~~~~~~~~~~~~~~~~~~~~~~~~~~~~~~~~~~~~~~~~~~~~~~~~~~~~~~~~~~~<1438
1468<~~~~~~~~~~~~~~~~~~~~~~~~~~~~~~~~~~~~~~~~~~~~~~~~~~~~~~~~~~~~~~~~~~~~~~~~~~~~~~~~~~~~~~~~~~~~~~~~~~~~<1468
1464<~~~~~~~~~~~~~~~~~~~~~~~~~~~~~~~~~~~~~~~~~~~~~~~~~~~~~~~~~~~~~~~~~~~~~~~~~~~~~~~~~~~~~~~~~~~~~~~~~~~~<1464
1407<~~~~~~~~~~~~~~~~~~~~~~~~~~~~~~~~~~~~~~~~~~~~~~~~~~~~~~~~~~~~~~~~~~~~~~~~~~~~~~~~~~~~~~~~~~~~~~~~~~~~<1407
1379<~~~~~~~~~~~~~~~~~~~~~~~~~~~~~~~~~~~~~~~~~~~~~~~~~~~~~~~~~~~~~~~~~~~~~~~~~~~~~~~~~~~~~~~~~~~~~~~~~~~~<1379
1410<~~~~~~~~~~~~~~~~~~~~~~~~~~~~~~~~~~~~~~~~~~~~~~~~~~~~~~~~~~~~~~~~~~~~~~~~~~~~~~~~~~~~~~~~~~~~~~~~~~~~<1410

              *         *         *         *         *         *         *         *         *         *
2701>attattaatcttttgtttcctcgtcattgttctcgttccctttcttccttgtttctttttctgcacaatatttcaagctataccaagcatacaatcaact>2800
1438<~~~~~~~~~~~~~~~~~~~~~~~~~~~~~~~~~~~~~~~~~~~~~~~~~~~~~~~~~~~~~~~~~~~~~~~~~~~~~~~~~~~~~~~~~~~~~~~~~~~~<1438
1468<~~~~~~~~~~~~~~~~~~~~~~~~~~~~~~~~~~~~~~~~~~~~~~~~~~~~~~~~~~~~~~~~~~~~~~~~~~~~~~~~~~~~~~~~~~~~~~~~~~~~<1468
1464<~~~~~~~~~~~~~~~~~~~~~~~~~~~~~~~~~~~~~~~~~~~~~~~~~~~~~~~~~~~~~~~~~~~~~~~~~~~~~~~~~~~~~~~~~~~~~~~~~~~~<1464
1407<~~~~~~~~~~~~~~~~~~~~~~~~~~~~~~~~~~~~~~~~~~~~~~~~~~~~~~~~~~~~~~~~~~~~~~~~~~~~~~~~~~~~~~~~~~~~~~~~~~~~<1407
1379<~~~~~~~~~~~~~~~~~~~~~~~~~~~~~~~~~~~~~~~~~~~~~~~~~~~~~~~~~~~~~~~~~~~~~~~~~~~~~~~~~~~~~~~~~~~~~~~~~~~~<1379
1410<~~~~~~~~~~~~~~~~~~~~~~~~~~~~~~~~~~~~~~~~~~~~~~~~~~~~~~~~~~~~~~~~~~~~~~~~~~~~~~~~~~~~~~~~~~~~~~~~~~~~<1410

              *         *         *         *         *         *         *         *         *         *
2801>atctcatatacatctagaactagtggatccccatcacaagtttgtacaaaaaagcaggcttcaaaatgatggcttcattgtcttgtgttgaagacaaga>2900
1438<~~~~~~~~~~~~~~~~~~~~~~~~~~~~~~~~~~~~~~~~~~~~~~~~~~~~~~~~~~~~~~~~~~~~~~~~~~~~~~~~~~~~~~~~~~~~~~~~~~~~<1438
1468<~~~~~~~~~~~~~~~~~~~~~~~~~~~~~~~~~~~~~~~~~~~~~~~~~~~~~~~~~~~~~~~~~~~~~~~~~~~~~~~~~~~~~~~~~~~~~~~~~~~~<1468
1464<~~~~~~~~~~~~~~~~~~~~~~~~~~~~~~~~~~~~~~~~~~~~~~~~~~~~~~~~~~~~~~~~~~~~~~~~~~~~~~~~~~~~~~~~~~~~~~~~~~~~<1464
1407<~~~~~~~~~~~~~~~~~~~~~~~~~~~~~~~~~~~~~~~~~~~~~~~~~~~~~~~~~~~~~~~~~~~~~~~~~~~~~~~~~~~~~~~~~~~~~~~~~~~~<1407
1379<~~~~~~~~~~~~~~~~~~~~~~~~~~~~~~~~~~~~~~~~~~~~~~~~~~~~~~~~~~~~~~~~~~~~~~~~~~~~~~~~~~~~~~~~~~~~~~~~~~~~<1379
1410<~~~~~~~~~~~~~~~~~~~~~~~~~~~~~~~~~~~~~~~~~~~~~~~~~~~~~~~~~~~~~~~~~~~~~~~~~~~~~~~~~~~~~~~~~~~~~~~~~~~~<1410

              *         *         *         *         *         *         *         *         *         *
2901>tgaaaacaagttgtttggttaatggtggaggaactataacaacaacaacatctcaatctaccttgcttgaagagatgaagctgttgaaagaccagtcagg>3000
1438<~~~~~~~~~~~~~~~~~~~~~~~~~~~~~~~~~~~~~~~~~~~~~~~~~~~~~~~~~~~~~~~~~~~~~~~~~~~~~~~~~~~~~~~~~~~~~~~~~~~~<1438
```

```
1468<~~~~~~~~~~~~~~~~~~~~~~~~~~~~~~~~~~~~~~~~~~~~~~~~~~~~~~~~~~~~~~~~~~~~~~~~~~~~~~~~~~~~~~~~~~~~~~~~~<1468
1464<~~~~~~~~~~~~~~~~~~~~~~~~~~~~~~~~~~~~~~~~~~~~~~~~~~~~~~~~~~~~~~~~~~~~~~~~~~~~~~~~~~~~~~~~~~~~~~~~<1464
1407<~~~~~~~~~~~~~~~~~~~~~~~~~~~~~~~~~~~~~~~~~~~~~~~~~~~~~~~~~~~~~~~~~~~~~~~~~~~~~~~~~~~~~~~~~~~~~~~~<1407
1379<~~~~~~~~~~~~~~~~~~~~~~~~~~~~~~~~~~~~~~~~~~~~~~~~~~~~~~~~~~~~~~~~~~~~~~~~~~~~~~~~~~~~~~~~~~~~~~~~<1379
1410<~~~~~~~~~~~~~~~~~~~~~~~~~~~~~~~~~~~~~~~~~~~~~~~~~~~~~~~~~~~~~~~~~~~~~~~~~~~~~~~~~~~~~~~~~~~~~~~~<1410

                  *         *         *         *         *         *         *         *         *         *
3001>tacaagaaagccggtaataaactcggagctatggcacgcttgtgcaggccctttggtgtgtctccctcaagttgggagcttagtgtattacttctcacaa>3100
1438<~~~~~~~~~~~~~~~~~~~~~~~~~~~~~~~~~~~~~~~~~~~~~~~~~~~~~~~~~~~~~~~~~~~~~~~~~~~~~~~~~~~~~~~~~~~~~~~~<1438
1468<~~~~~~~~~~~~~~~~~~~~~~~~~~~~~~~~~~~~~~~~~~~~~~~~~~~~~~~~~~~~~~~~~~~~~~~~~~~~~~~~~~~~~~~~~~~~~~~~<1468
1464<~~~~~~~~~~~~~~~~~~~~~~~~~~~~~~~~~~~~~~~~~~~~~~~~~~~~~~~~~~~~~~~~~~~~~~~~~~~~~~~~~~~~~~~~~~~~~~~~<1464
1407<~~~~~~~~~~~~~~~~~~~~~~~~~~~~~~~~~~~~~~~~~~~~~~~~~~~~~~~~~~~~~~~~~~~~~~~~~~~~~~~~~~~~~~~~~~~~~~~~<1407
1379<~~~~~~~~~~~~~~~~~~~~~~~~~~~~~~~~~~~~~~~~~~~~~~~~~~~~~~~~~~~~~~~~~~~~~~~~~~~~~~~~~~~~~~~~~~~~~~~~<1379
1410<~~~~~~~~~~~~~~~~~~~~~~~~~~~~~~~~~~~~~~~~~~~~~~~~~~~~~~~~~~~~~~~~~~~~~~~~~~~~~~~~~~~~~~~~~~~~~~~~<1410

                  *         *         *         *         *         *         *         *         *         *
3101>ggtcatagcgagcaggttgctgtttcaaccagaagatcagcaacaacacaagttcctaattatccgaaccttccatctcagttgatgtgtcaagtccata>3200
1438<~~~~~~~~~~~~~~~~~~~~~~~~~~~~~~~~~~~~~~~~~~~~~~~~~~~~~~~~~~~~~~~~~~~~~~~~~~~~~~~~~~~~~~~~~~~~~~~~<1438
1468<~~~~~~~~~~~~~~~~~~~~~~~~~~~~~~~~~~~~~~~~~~~~~~~~~~~~~~~~~~~~~~~~~~~~~~~~~~~~~~~~~~~~~~~~~~~~~~~~<1468
1464<~~~~~~~~~~~~~~~~~~~~~~~~~~~~~~~~~~~~~~~~~~~~~~~~~~~~~~~~~~~~~~~~~~~~~~~~~~~~~~~~~~~~~~~~~~~~~~~~<1464
1407<~~~~~~~~~~~~~~~~~~~~~~~~~~~~~~~~~~~~~~~~~~~~~~~~~~~~~~~~~~~~~~~~~~~~~~~~~~~~~~~~~~~~~~~~~~~~~~~~<1407
1379<~~~~~~~~~~~~~~~~~~~~~~~~~~~~~~~~~~~~~~~~~~~~~~~~~~~~~~~~~~~~~~~~~~~~~~~~~~~~~~~~~~~~~~~~~~~~~~~~<1379
1410<~~~~~~~~~~~~~~~~~~~~~~~~~~~~~~~~~~~~~~~~~~~~~~~~~~~~~~~~~~~~~~~~~~~~~~~~~~~~~~~~~~~~~~~~~~~~~~~~<1410

                  *         *         *         *         *         *         *         *         *         *
3201>atgttactcttcatgctgacaaagacagtgacgaaatctatgctcagatgagtcttcaacctgttcactctgagagagatgtgttccctgtaccagactt>3300
1438<~~~~~~~~~~~~~~~~~~~~~~~~~~~~~~~~~~~~~~~~~~~~~~~~~~~~~~~~~~~~~~~~~~~~~~~~~~~~~~~~~~~~~~~~~~~~~~~~<1438
1468<~~~~~~~~~~~~~~~~~~~~~~~~~~~~~~~~~~~~~~~~~~~~~~~~~~~~~~~~~~~~~~~~~~~~~~~~~~~~~~~~~~~~~~~~~~~~~~~~<1468
1464<~~~~~~~~~~~~~~~~~~~~~~~~~~~~~~~~~~~~~~~~~~~~~~~~~~~~~~~~~~~~~~~~~~~~~~~~~~~~~~~~~~~~~~~~~~~~~~~~<1464
1407<~~~~~~~~~~~~~~~~~~~~~~~~~~~~~~~~~~~~~~~~~~~~~~~~~~~~~~~~~~~~~~~~~~~~~~~~~~~~~~~~~~~~~~~~~~~~~~~~<1407
1379<~~~~~~~~~~~~~~~~~~~~~~~~~~~~~~~~~~~~~~~~~~~~~~~~~~~~~~~~~~~~~~~~~~~~~~~~~~~~~~~~~~~~~~~~~~~~~~~~<1379
1410<~~~~~~~~~~~~~~~~~~~~~~~~~~~~~~~~~~~~~~~~~~~~~~~~~~~~~~~~~~~~~~~~~~~~~~~~~~~~~~~~~~~~~~~~~~~~~~~~<1410

                  *         *         *         *         *         *         *         *         *         *
3301>tggaatgctgagaggaagtaagcacccgactgagtttttctgcaaaacacttactgcaagtgacacaagcacacatggaggtttctcagtgccacgtaga>3400
1438<~~~~~~~~~~~~~~~~~~~~~~~~~~~~~~~~~~~~~~~~~~~~~~~~~~~~~~~~~~~~~~~~~~~~~~~~~~~~~~~~~~~~~~~~~~~~~~~~<1438
1468<~~~~~~~~~~~~~~~~~~~~~~~~~~~~~~~~~~~~~~~~~~~~~~~~~~~~~~~~~~~~~~~~~~~~~~~~~~~~~~~~~~~~~~~~~~~~~~~~<1468
1464<~~~~~~~~~~~~~~~~~~~~~~~~~~~~~~~~~~~~~~~~~~~~~~~~~~~~~~~~~~~~~~~~~~~~~~~~~~~~~~~~~~~~~~~~~~~~~~~~<1464
1407<~~~~~~~~~~~~~~~~~~~~~~~~~~~~~~~~~~~~~~~~~~~~~~~~~~~~~~~~~~~~~~~~~~~~~~~~~~~~~~~~~~~~~~~~~~~~~~~~<1407
```

```
1379<~~~~~~~~~~~~~~~~~~~~~~~~~~~~~~~~~~~~~~~~~~~~~~~~~~~~~~~~~~~~~~~~~~~~~~~~~~~~~~~~~~~~~~~~~~~~~~~~~<1379
1410<~~~~~~~~~~~~~~~~~~~~~~~~~~~~~~~~~~~~~~~~~~~~~~~~~~~~~~~~~~~~~~~~~~~~~~~~~~~~~~~~~~~~~~~~~~~~~~~~<1410

                  *         *         *         *         *         *         *         *         *         *
3401>gctgcagagaagctatttccaccattggactactcagcacagccgccaacgcaagagcttgtagttcgagatcttcatgagaatacttggacatttcgcc>3500
1438<~~~~~~~~~~~~~~~~~~~~~~~~~~~~~~~~~~~~~~~~~~~~~~~~~~~~~~~~~~~~~~~~~~~~~~~~~~~~~~~~~~~~~~~~~~~~~~~~<1438
1468<~~~~~~~~~~~~~~~~~~~~~~~~~~~~~~~~~~~~~~~~~~~~~~~~~~~~~~~~~~~~~~~~~~~~~~~~~~~~~~~~~~~~~~~~~~~~~~~~<1468
1464<~~~~~~~~~~~~~~~~~~~~~~~~~~~~~~~~~~~~~~~~~~~~~~~~~~~~~~~~~~~~~~~~~~~~~~~~~~~~~~~~~~~~~~~~~~~~~~~~<1464
1407<~~~~~~~~~~~~~~~~~~~~~~~~~~~~~~~~~~~~~~~~~~~~~~~~~~~~~~~~~~~~~~~~~~~~~~~~~~~~~~~~~~~~~~~~~~~~~~~~<1407
1379<~~~~~~~~~~~~~~~~~~~~~~~~~~~~~~~~~~~~~~~~~~~~~~~~~~~~~~~~~~~~~~~~~~~~~~~~~~~~~~~~~~~~~~~~~~~~~~~~<1379
1410<~~~~~~~~~~~~~~~~~~~~~~~~~~~~~~~~~~~~~~~~~~~~~~~~~~~~~~~~~~~~~~~~~~~~~~~~~~~~~~~~~~~~~~~~~~~~~~~~<1410

                  *         *         *         *         *         *         *         *         *         *
3501>atatctaccgagggcaaccaaagagacatctcctaactacaggatggagtttgttcgttggatcgaagagattgagagctggggattctgttttgttcat>3600
1438<~~~~~~~~~~~~~~~~~~~~~~~~~~~~~~~~~~~~~~~~~~~~~~~~~~~~~~~~~~~~~~~~~~~~~~~~~~~~~~~~~~~~~~~~~~~~~~~~<1438
1468<~~~~~~~~~~~~~~~~~~~~~~~~~~~~~~~~~~~~~~~~~~~~~~~~~~~~~~~~~~~~~~~~~~~~~~~~~~~~~~~~~~~~~~~~~~~~~~~~<1468
1464<~~~~~~~~~~~~~~~~~~~~~~~~~~~~~~~~~~~~~~~~~~~~~~~~~~~~~~~~~~~~~~~~~~~~~~~~~~~~~~~~~~~~~~~~~~~~~~~~<1464
1407<~~~~~~~~~~~~~~~~~~~~~~~~~~~~~~~~~~~~~~~~~~~~~~~~~~~~~~~~~~~~~~~~~~~~~~~~~~~~~~~~~~~~~~~~~~~~~~~~<1407
1379<~~~~~~~~~~~~~~~~~~~~~~~~~~~~~~~~~~~~~~~~~~~~~~~~~~~~~~~~~~~~~~~~~~~~~~~~~~~~~~~~~~~~~~~~~~~~~~~~<1379
1410<~~~~~~~~~~~~~~~~~~~~~~~~~~~~~~~~~~~~~~~~~~~~~~~~~~~~~~~~~~~~~~~~~~~~~~~~~~~~~~~~~~~~~~~~~~~~~~~~<1410

                  *         *         *         *         *         *         *         *         *         *
3601>cagggatgagaagtcacaacttatggtcggtgttaggcgtgccaatcgccaacaaacagcacttccttcatcagttctctcagcggatagtatgcacatc>3700
1438<~~~~~~~~~~~~~~~~~~~~~~~~~~~~~~~~~~~~~~~~~~~~~~~~~~~~~~~~~~~~~~~~~~~~~~~~~~~~~~~~~~~~~~~~~~~~~~~~<1438
1468<~~~~~~~~~~~~~~~~~~~~~~~~~~~~~~~~~~~~~~~~~~~~~~~~~~~~~~~~~~~~~~~~~~~~~~~~~~~~~~~~~~~~~~~~~~~~~~~~<1468
1464<~~~~~~~~~~~~~~~~~~~~~~~~~~~~~~~~~~~~~~~~~~~~~~~~~~~~~~~~~~~~~~~~~~~~~~~~~~~~~~~~~~~~~~~~~~~~~~~~<1464
1407<~~~~~~~~~~~~~~~~~~~~~~~~~~~~~~~~~~~~~~~~~~~~~~~~~~~~~~~~~~~~~~~~~~~~~~~~~~~~~~~~~~~~~~~~~~~~~~~~<1407
1379<~~~~~~~~~~~~~~~~~~~~~~~~~~~~~~~~~~~~~~~~~~~~~~~~~~~~~~~~~~~~~~~~~~~~~~~~~~~~~~~~~~~~~~~~~~~~~~~~<1379
1410<~~~~~~~~~~~~~~~~~~~~~~~~~~~~~~~~~~~~~~~~~~~~~~~~~~~~~~~~~~~~~~~~~~~~~~~~~~~~~~~~~~~~~~~~~~~~~~~~<1410

                  *         *         *         *         *         *         *         *         *         *
3701>ggtgttcttgctgctgctgctcacgcaaccgccaaccgtactcctttttttgatattctataatccaagagcttgtccagcagagttcgtgatccctctag>3800
1438<~~~~~~~~~~~~~~~~~~~~~~~~~~~~~~~~~~~~~~~~~~~~~~~~~~~~~~~~~~~~~~~~~~~~~~~~~~~~~~~~~~~~~~~~~~~~~~~~<1438
1468<~~~~~~~~~~~~~~~~~~~~~~~~~~~~~~~~~~~~~~~~~~~~~~~~~~~~~~~~~~~~~~~~~~~~~~~~~~~~~~~~~~~~~~~~~~~~~~~~<1468
1464<~~~~~~~~~~~~~~~~~~~~~~~~~~~~~~~~~~~~~~~~~~~~~~~~~~~~~~~~~~~~~~~~~~~~~~~~~~~~~~~~~~~~~~~~~~~~~~~~<1464
1407<~~~~~~~~~~~~~~~~~~~~~~~~~~~~~~~~~~~~~~~~~~~~~~~~~~~~~~~~~~~~~~~~~~~~~~~~~~~~~~~~~~~~~~~~~~~~~~~~<1407
1379<~~~~~~~~~~~~~~~~~~~~~~~~~~~~~~~~~~~~~~~~~~~~~~~~~~~~~~~~~~~~~~~~~~~~~~~~~~~~~~~~~~~~~~~~~~~~~~~~<1379
1410<~~~~~~~~~~~~~~~~~~~~~~~~~~~~~~~~~~~~~~~~~~~~~~~~~~~~~~~~~~~~~~~~~~~~~~~~~~~~~~~~~~~~~~~~~~~~~~~~<1410
```

```
              *         *         *         *         *         *         *         *         *         *
3801>ctaagtaccgtaaggcgatatgcgggtctcagctctcagttggtatgagatttggaatgatgtttgaaactgaagattccgggaaacgaaggtacatggg>3900
1438<~~~~~~~~~~~~~~~~~~~~~~~~~~~~~~~~~~~~~~~~~~~~~~~~~~~~~~~~~~~~~~~~~~~~~~~~~~~~~~~~~~~~~~~~~~~~~~~~~~<1438
1468<~~~~~~~~~~~~~~~~~~~~~~~~~~~~~~~~~~~~~~~~~~~~~~~~~~~~~~~~~~~~~~~~~~~~~~~~~~~~~~~~~~~~~~~~~~~~~~~~~~<1468
1464<~~~~~~~~~~~~~~~~~~~~~~~~~~~~~~~~~~~~~~~~~~~~~~~~~~~~~~~~~~~~~~~~~~~~~~~~~~~~~~~~~~~~~~~~~~~~~~~~~~<1464
1407<~~~~~~~~~~~~~~~~~~~~~~~~~~~~~~~~~~~~~~~~~~~~~~~~~~~~~~~~~~~~~~~~~~~~~~~~~~~~~~~~~~~~~~~~~~~~~~~~~~<1407
1379<~~~~~~~~~~~~~~~~~~~~~~~~~~~~~~~~~~~~~~~~~~~~~~~~~~~~~~~~~~~~~~~~~~~~~~~~~~~~~~~~~~~~~~~~~~~~~~~~~~<1379
1410<~~~~~~~~~~~~~~~~~~~~~~~~~~~~~~~~~~~~~~~~~~~~~~~~~~~~~~~~~~~~~~~~~~~~~~~~~~~~~~~~~~~~~~~~~~~~~~~~~~<1410

              *         *         *         *         *         *         *         *         *         *
3901>aactattgttggaatcagcgatttggatccgttgagatggcctggttctaagtggcgtaaccttcaggtagaatgggatgagcctggatgtaatgataaa>4000
1438<~~~~~~~~~~~~~~~~~~~~~~~~~~~~~~~~~~~~~~~~~~~~~~~~~~~~~~~~~~~~~~~~~~~~~~~~~~~~~~~~~~~~~~~~~~~~~~~~~~<1438
1468<~~~~~~~~~~~~~~~~~~~~~~~~~~~~~~~~~~~~~~~~~~~~~~~~~~~~~~~~~~~~~~~~~~~~~~~~~~~~~~~~~~~~~~~~~~~~~~~~~~<1468
1464<~~~~~~~~~~~~~~~~~~~~~~~~~~~~~~~~~~~~~~~~~~~~~~~~~~~~~~~~~~~~~~~~~~~~~~~~~~~~~~~~~~~~~~~~~~~~~~~~~~<1464
1407<~~~~~~~~~~~~~~~~~~~~~~~~~~~~~~~~~~~~~~~~~~~~~~~~~~~~~~~~~~~~~~~~~~~~~~~~~~~~~~~~~~~~~~~~~~~~~~~~~~<1407
1379<~~~~~~~~~~~~~~~~~~~~~~~~~~~~~~~~~~~~~~~~~~~~~~~~~~~~~~~~~~~~~~~~~~~~~~~~~~~~~~~~~~~~~~~~~~~~~~~~~~<1379
1410<~~~~~~~~~~~~~~~~~~~~~~~~~~~~~~~~~~~~~~~~~~~~~~~~~~~~~~~~~~~~~~~~~~~~~~~~~~~~~~~~~~~~~~~~~~~~~~~~~~<1410

              *         *         *         *         *         *         *         *         *         *
4001>cctactcgggtcagtccatgggatatcgaaacacctgaaagtctcttcatttttccttctctgacctcaggactcaaacgtcagctccatccatcttact>4100
1438<~~~~~~~~~~~~~~~~~~~~~~~~~~~~~~~~~~~~~~~~~~~~~~~~~~~~~~~~~~~~~~~~~~~~~~~~~~~~~~~~~~~~~~~~~~~~~~~~~~<1438
1468<~~~~~~~~~~~~~~~~~~~~~~~~~~~~~~~~~~~~~~~~~~~~~~~~~~~~~~~~~~~~~~~~~~~~~~~~~~~~~~~~~~~~~~~~~~~~~~~~~~<1468
1464<~~~~~~~~~~~~~~~~~~~~~~~~~~~~~~~~~~~~~~~~~~~~~~~~~~~~~~~~~~~~~~~~~~~~~~~~~~~~~~~~~~~~~~~~~~~~~~~~~~<1464
1407<~~~~~~~~~~~~~~~~~~~~~~~~~~~~~~~~~~~~~~~~~~~~~~~~~~~~~~~~~~~~~~~~~~~~~~~~~~~~~~~~~~~~~~~~~~~~~~~~~~<1407
1379<~~~~~~~~~~~~~~~~~~~~~~~~~~~~~~~~~~~~~~~~~~~~~~~~~~~~~~~~~~~~~~~~~~~~~~~~~~~~~~~~~~~~~~~~~~~~~~~~~~<1379
1410<~~~~~~~~~~~~~~~~~~~~~~~~~~~~~~~~~~~~~~~~~~~~~~~~~~~~~~~~~~~~~~~~~~~~~~~~~~~~~~~~~~~~~~~~~~~~~~~~~~<1410

              *         *         *         *         *         *         *         *         *         *
4101>ttgctggtgaaactgaatggggtagcttgataaaacggccacttatacgtgttcctgattccgcgaatgggattatgccatatgcatctttccctagtat>4200
1438<~~~~~~~~~~~~~~~~~~~~~~~~~~~~~~~~~~~~~~~~~~~~~~~~~~~~~~~~~~~~~~~~~~~~~~~~~~~~~~~~~~~~~~~~~~~~~~~~~~<1438
1468<~~~~~~~~~~~~~~~~~~~~~~~~~~~~~~~~~~~~~~~~~~~~~~~~~~~~~~~~~~~~~~~~~~~~~~~~~~~~~~~~~~~~~~~~~~~~~~~~~~<1468
1464<~~~~~~~~~~~~~~~~~~~~~~~~~~~~~~~~~~~~~~~~~~~~~~~~~~~~~~~~~~~~~~~~~~~~~~~~~~~~~~~~~~~~~~~~~~~~~~~~~~<1464
1407<~~~~~~~~~~~~~~~~~~~~~~~~~~~~~~~~~~~~~~~~~~~~~~~~~~~~~~~~~~~~~~~~~~~~~~~~~~~~~~~~~~~~~~~~~~~~~~~~~~<1407
1379<~~~~~~~~~~~~~~~~~~~~~~~~~~~~~~~~~~~~~~~~~~~~~~~~~~~~~~~~~~~~~~~~~~~~~~~~~~~~~~~~~~~~~~~~~~~~~~~~~~<1379
1410<~~~~~~~~~~~~~~~~~~~~~~~~~~~~~~~~~~~~~~~~~~~~~~~~~~~~~~~~~~~~~~~~~~~~~~~~~~~~~~~~~~~~~~~~~~~~~~~~~~<1410

              *         *         *         *         *         *         *         *         *         *
4201>ggcttcggagcagcttatgaaaatgatgatgaggcctcacaacaaccaaaatgtaccatctttcatgtctgagatgcagcagaatattgtaatggggaat>4300
```

```
1438<~~~~~~~~~~~~~~~~~~~~~~~~~~~~~~~~~~~~~~~~~~~~~~~~~~~~~~~~~~~~~~~~~~~~~~~~~~~~~~~~~~~~~~~~~~~~~~~~~~~~<1438
1468<~~~~~~~~~~~~~~~~~~~~~~~~~~~~~~~~~~~~~~~~~~~~~~~~~~~~~~~~~~~~~~~~~~~~~~~~~~~~~~~~~~~~~~~~~~~~~~~~~~~~<1468
1464<~~~~~~~~~~~~~~~~~~~~~~~~~~~~~~~~~~~~~~~~~~~~~~~~~~~~~~~~~~~~~~~~~~~~~~~~~~~~~~~~~~~~~~~~~~~~~~~~~~~~<1464
1407<~~~~~~~~~~~~~~~~~~~~~~~~~~~~~~~~~~~~~~~~~~~~~~~~~~~~~~~~~~~~~~~~~~~~~~~~~~~~~~~~~~~~~~~~~~~~~~~~~~~~<1407
1379<~~~~~~~~~~~~~~~~~~~~~~~~~~~~~~~~~~~~~~~~~~~~~~~~~~~~~~~~~~~~~~~~~~~~~~~~~~~~~~~~~~~~~~~~~~~~~~~~~~~~<1379
1410<~~~~~~~~~~~~~~~~~~~~~~~~~~~~~~~~~~~~~~~~~~~~~~~~~~~~~~~~~~~~~~~~~~~~~~~~~~~~~~~~~~~~~~~~~~~~~~~~~~~~<1410
```

```
              *         *         *         *         *         *         *         *         *         *
4301>ggaggtttgctaggagatatgaagatgcagcaacccctgatgatgaaccagaaatctgagatggtgcagccacaaaacaagctaacagtgaacccatctg>4400
1438<~~~~~~~~~~~~~~~~~~~~~~~~~~~~~~~~~~~~~~~~~~~~~~~~~~~~~~~~~~~~~~~~~~~~~~~~~~~~~~~~~~~~~~~~~~~~~~~~~~~~<1438
1468<~~~~~~~~~~~~~~~~~~~~~~~~~~~~~~~~~~~~~~~~~~~~~~~~~~~~~~~~~~~~~~~~~~~~~~~~~~~~~~~~~~~~~~~~~~~~~~~~~~~~<1468
1464<~~~~~~~~~~~~~~~~~~~~~~~~~~~~~~~~~~~~~~~~~~~~~~~~~~~~~~~~~~~~~~~~~~~~~~~~~~~~~~~~~~~~~~~~~~~~~~~~~~~~<1464
1407<~~~~~~~~~~~~~~~~~~~~~~~~~~~~~~~~~~~~~~~~~~~~~~~~~~~~~~~~~~~~~~~~~~~~~~~~~~~~~~~~~~~~~~~~~~~~~~~~~~~~<1407
1379<~~~~~~~~~~~~~~~~~~~~~~~~~~~~~~~~~~~~~~~~~~~~~~~~~~~~~~~~~~~~~~~~~~~~~~~~~~~~~~~~~~~~~~~~~~~~~~~~~~~~<1379
1410<~~~~~~~~~~~~~~~~~~~~~~~~~~~~~~~~~~~~~~~~~~~~~~~~~~~~~~~~~~~~~~~~~~~~~~~~~~~~~~~~~~~~~~~~~~~~~~~~~~~~<1410
```

```
              *         *         *         *         *         *         *         *         *         *
4401>cttctaatacgagtggccaagaacagaatctttcacagagtatgagtgctcctgctaaacctgagaactctacactctctggttgcagctctggtagagt>4500
1438<~~~~~~~~~~~~~~~~~~~~~~~~~~~~~~~~~~~~~~~~~~~~~~~~~~~~~~~~~~~~~~~~~~~~~~~~~~~~~~~~~~~~~~~~~~~~~~~~~~~~<1438
1467<~~~~~~~~~~~~~~~~~~~~~~~~~~~~~~~~~~~~~~~~~~~~~~~~~~~~~~~~~~~~~~~~~~~~~~~~~~~~~~~~~~~~~~~~~~~~~~~~~~CT<1466
1464<~~~~~~~~~~~~~~~~~~~~~~~~~~~~~~~~~~~~~~~~~~~~~~~~~~~~~~~~~~~~~~~~~~~~~~~~~~~~~~~~~~~~~~~~~~~~~~~~~~~~<1464
1407<~~~~~~~~~~~~~~~~~~~~~~~~~~~~~~~~~~~~~~~~~~~~~~~~~~~~~~~~~~~~~~~~~~~~~~~~~~~~~~~~~~~~~~~~~~~~~~~~~~~~<1407
1379<~~~~~~~~~~~~~~~~~~~~~~~~~~~~~~~~~~~~~~~~~~~~~~~~~~~~~~~~~~~~~~~~~~~~~~~~~~~~~~~~~~~~~~~~~~~~~~~~~~~~<1379
1410<~~~~~~~~~~~~~~~~~~~~~~~~~~~~~~~~~~~~~~~~~~~~~~~~~~~~~~~~~~~~~~~~~~~~~~~~~~~~~~~~~~~~~~~~~~~~~~~~~~~~<1410
```

```
              *         *         *         *             *         *           *         *          *
4501>ccaacatggacttgagcagtcaatggaacaggcaagccaggttact-ac-a--tccacagtgtgta-atgagga-aaaggttaatcagctactt-cagaa>4593
1437<~~~~~~~~~~~~~~~~~~~~~~~~~~~~~~GGCAAGCCAGGTTACTCACTATCTCCACAGTGTGTATATGA-GAGAAAGGTTATTCAGCTACTTCCAG-A<1370
1465<TGAGAGCACGTTCAATATGGAAACACGGCCAAGCCCAGTTATACTA-CA-T--CCCACAGTGTGTA-TATGAAG-AAAGGTTATTCAGCTATCT-CAGAA<1373
1463<~~~~~CAAATCATATGGACTTGAGTGACGTCTAATGGAAACAGCAG-CC-A--GTATACACTCCAC-GTGGTAA-TGAGGAATAGTTATTTCAG-TACTC<1376
1406<~~~~~~~~~~~~~~~~~~~~~~~~~~~~~~~~~~~~~~~~~~~~~~~~~~~~~~~~~~~~~~~~~C-AAGTGGT-GTAATGAGGAAAGTATCAC-GTACT<1375
1379<~~~~~~~~~~~~~~~~~~~~~~~~~~~~~~~~~~~~~~~~~~~~~~~~~~~~~~~~~~~~~~~~~~~~~~~~~~~~~~~~~~~~~~~~~~~~~~~~~~~~<1379
1410<~~~~~~~~~~~~~~~~~~~~~~~~~~~~~~~~~~~~~~~~~~~~~~~~~~~~~~~~~~~~~~~~~~~~~~~~~~~~~~~~~~~~~~~~~~~~~~~~~~~~<1410
```

```
           *         *         *          *         *         *         *          *         *         *
4594>accgggtgcttcgtcgcctgtacaagctgat-caatgtcttgacattactcatcagatttaccaaccacag-tctgatccaataaatggattctctttcc>4691
1369<ACC-GGTGC-TCGTCGCCTGTACAAGCTGATTC-ATGTCTTGACA-TACTCATCAGATTTACC-ACCACAGTTCTG-TTCAATAAAT-GATTCTCTTT-C<1278
1372<CCCGGTTGCTTCTGTTCGCCTTACAAGCTGA-TTCAATTCTTGACATTATCATCAGATTTACCACCACACA-GTTCTGTTCCAATAATGATTCTCCTTCC<1275
1375<AGAACCCGGTGCTCTCGTTCGCCTGTACAAG-CTGATCATGTCTTGACAATACTTCATCAGATTTACACAC-AGTCTGTTCAATAATGGATTCTCTCTTT<1278
```

```
1374<CAGAAACGGTTGGCTTTCGTGCCTGTAACAA-GGTGGATTCAATGTCTTGACATACTCATCAGATTACACA-CAGTCTGATCAATAATGGATCTTCTTTG<1277
1378<~~~~~~~~~~~~~~~~~~~~~~~~~~~~~~~~~~~~~~~~~~~~~~~~~~~~~~CATGTGTCTGTGACATA-TTCATCAGCTACACACAGTCGATTCATA<1334
1409<~~~~~~~~~~~~~~~~~~~~~~~~~~~ACGA-TCTACAGTGACCTGTCAGCTGTACAGTCCTGAACATATC-CTCAGGTACATCCAGGTTCGATCCAATA<1339

             *         *         *          *          *         *            *         *         *
4692>tggaaactgatgagctgacatcacaagtct-cttccttccagtc-tcttgccggatcatacaa-gcaac-cattcattctatcctcccaggattcttcag>4787
1277<TGGAAACTGATGAGCTGACATCACAAGTCT-CTTC--TGCAGTC-TCTTGCC-GATCATACAA-GCA--GCATTCATTCT-TTCTCCCAGGATTCTTCAG<1187
1274<TGGAAACTGATGAGCTGACATCACAAGTCT-CT-CCT--CAGTC-TC-TGCCGGATCATACAA-GCAAC-CATTCATTCTATC-TCCCAGGATTC-TCAG<1185
1277<CTGAAACTGATGAGCTGACATCACAAGTCTTCTTCC-TCCAGTC-TC-TGC--GATCATACAA-GCAAC-CATTCATTCTATCCTCC-AGGA-TC-TCAG<1188
1276<CTGAACTGATGAGCTGAACATCACAAGTCT-CTTC--TGCAGTCTTCTTGCGGGATCATACAAGGCA--GCA-TCATTCTAT-CTCCCAGGA-TC-TCAG<1186
1333<TGATTTCTCTTACCTTGACTGATGAGCTGA-CATCCCAAGTTCT-CTTCTGCAGTCTCTTGCG-ATCAT-ACCAGCATCAATCATCTATTCCTCCAGATT<1238
1338<ATTGATTCTCTTGCTGACTGATGAGGCTGA-CCATCACAAGTCT-CTCCTGCAGTCTCTTGCG-ATCAT-ACAAGCACCATCATTCTTATCCTCCCAGAT<1243

       *         *          *         *         *         *         *         *         *         *
4788>ctgttgtgttaccggattc-cacaaactcaccgctgtttcatgatgtgtgggacactcagttgaacggtctcaagtttgaccagttcagtcccttgatgc>4886
1186<CTGTTGTGTTACCGGATTC-CACAAACTCACCGCTGTTTCATGATGTGTGGGACACTCAGTTGAACGGTCTCAAGTTTGACCAGTTCAGTCCCTTGATGC<1088
1184<CTG-TGTG-TACCGGA-TC-CACAAACTCACCGCTGTTTCATGATGTGTGGGACACTCAGTTGAACGGTCTCAAGTTTGACCAGTTCAGTCCCTTGATGC<1089
1187<CTGTTGTGTTACCGGATTC-CACAAACTCACCGCTGTTTCATGATGTGTGGGACACTCAGTTGAACGGTCTCAAGTTTGACCAGTTCAGTCCCTTGATGC<1089
1185<CTGTTGTGTTA-CGGAT--GCACAAACTCACCGCTGTTTCATGATGTGTGGGACACTCAGTTGAACGGTCTCAAGTTTGACCAGTTCAGTCCCTTGATGC<1089
1237<CTTCAGCTGTGGTACTGAT-CCACAACTCACCGCTGTTTCATGATGTGTGGGACA-TCAGTTGAAC-GTCTCAAGTTTGA-CAG-TCAGTCCCTTGATGC<1143
1242<CTCAGCTGTGTGTACGATC-CACCAACTCCACCGCTGTTCATGATGTGTGGGACACTCAGT-GAACGGTCTCAAGTTTGATCAGTTCAGTCCCTTGATGC<1145

        *         *         *         *         *         *         *         *         *         *
4887>agcaggacctttatgctagtcagaatatctgtatgagtaatagcacaaccagtaacattctagatcctccactctcaaacacagtccttgatgacttctg>4986
1087<AGCAGGACCTTTATGCTAGTCAGAATATCTGTATGAGTAATAGCACAACCAGTAACATTCTAGATCCTCCACTCTCAAACACAGTCCTTGATGACTTCTG<988
1088<AGCAGGACCTTTATGCTAGTCAGAATATCTGTATGAGTAATAGCACAACCAGTAACATTCTAGATCCTCCACTCTCAAACACAGTCCTTGATGACTTCTG<989
1088<AGCAGGACCTTTATGCTAGTCAGAATATCTGTATGAGTAATAGCACAACCAGTAACATTCTAGATCCTCCACTCTCAAACACAGTCCTTGATGACTTCTG<989
1088<AGCAGGACCTT-ATGCTAGTCAGAATATCTGTATGAGTAATAGCACAACCAGTAACATTCTAGATCCTCCACTCTCAAACACAGTCCTTGATGACTTCTG<990
1142<AGCAGGACCTTTATGCTAGTCAGA-TATCTGTATGAGT-ATAGCACAACCAGTAACATTCTAGATCCTCCACTCTCAAACACAGTCCTTGATGACTTCTG<1045
1144<AGCAGGACCTTTATGCTAGTCAGA-TATCTGTATGAGTAATAGCAC-ACCAGTAACATTCTAGATCCTCCACTCTCAAACACAGTCCTTGATGACTTCTG<1047

        *         *         *         *          *         *         *         *         *         *
4987>tgccatcaaagacactgatttccagaaccacccctt-ctggttgtttggttggaaacaacaacactagctttgctcaagatgtccagtcgcagatcacatc>5085
 987<TGCCATCAAAGACACTGATTTCCAGAACCACCCCTT-CTGGTTGTTTGGTTGGAAACAACAACACTAGCTTTGCTCAAGATGTCCAGTCGCAGATCACATC<889
 988<TGCCATCAAAGACACTGATTTCCAGAACCACCCCTTTCTGGTTGTTTGGTTGGAAACAACAACACTAGCTTTGCTCAAGATGTCCAGTCGCAGATCACATC<889
 988<TGCCATCAAAGACACTGATTTCCAGAACCACCCCTT-CTGGTTGTTTGGTTGGAAACAACAACACTAGCTTTGCTCAAGATGTCCAGTCGCAGATCACATC<890
 989<TGCCATCAAAGACACTGATTTCCAGAACCACCCCTTTCTGGTTGTTTGGTTGGAAACAACAACACTAGCTTTGCTCAAGATGTCCAGTCGCAGATCACATC<890
1044<TGCCATCAAAGACACTGATTTCCAGAACCACCCCTT-CTGGTTGTTTGGTTGGAAACAACAACACTAGCTTTGCTCAAGATGTCCAGTCGCAGATCACATC<946
1046<TGCCATCAAAGACACTGATTTCCAGAACCACCTTT-CTGGTTGTTTGGTTGGAAACAACAACACTAGCTTTGCTCAAGATGTCCAGTCGCAGATCACATC<948
```

```
              *         *         *         *         *         *         *         *         *         *
5086>agctagctttgcagactcacaggccttctctcgccaagattttccagataattctggaggcactggtacatcttcaagcaatgttgattttgatgattgt>5185
 888<AGCTAGCTTTGCAGACTCACAGGCCTTCTCTCGCCAAGATTTTCCAGATAATTCTGGAGGCACTGGTACATCTTCAAGCAATGTTGATTTTGATGATTGT<789
 888<AGCTAGCTTTGCAGACTCACAGGCCTTCTCTCGCCAAGATTTTCCAGATAATTCTGGAGGCACTGGTACATCTTCAAGCAATGTTGATTTTGATGATTGT<789
 889<AGCTAGCTTTGCAGACTCACAGGCCTTCTCTCGCCAAGATTTTCCAGATAATTCTGGAGGCACTGGTACATCTTCAAGCAATGTTGATTTTGATGATTGT<790
 889<AGCTAGCTTTGCAGACTCACAGGCCTTCTCTCGCCAAGATTTTCCAGATAATTCTGGAGGCACTGGTACATCTTCAAGCAATGTTGATTTTGATGATTGT<790
 945<AGCTAGCTTTGCAGACTCACAGGCCTTCTCTCGCCAAGATTTTCCAGATAATTCTGGAGGCACTGGTACATCTTCAAGCAATGTTGATTTTGATGATTGT<846
 947<AGCTAGCTTTGCATACTCACAGGCCTTCTCTCGCCAAGATTTTCCAGATAATTCTGGAGGCACTGGTACATCTTCAAGCAATGTTGATTTTGATGATTGT<848

              *         *         *         *         *         *         *         *         *         *
5186>agtctgcggcaaaatagtaaaggctcatcatggcagaaaattgcgacaccccgcgtccgaaccggcagcgatctgggtaaaaagctgctggaagcagccg>5285
 788<AGTCTGCGGCAAAATAGTAAAGGCTCATCATGGCAGAAAATTGCGACACCCCGCGTCCGAACCGGCAGCGATCTGGGTAAAAAGCTGCTGGAAGCAGCCG<689
 788<AGTCTGCGGCAAAATAGTAAAGGCTCATCATGGCAGAAAATTGCGACACCCCGCGTCCGAACCGGCAGCGATCTGGGTAAAAAGCTGCTGGAAGCAGCCG<689
 789<AGTCTGCGGCAAAATAGTAAAGGCTCATCATGGCAGAAAATTGCGACACCCCGCGTCCGAACCGGCAGCGATCTGGGTAAAAAGCTGCTGGAAGCAGCCG<690
 789<AGTCTGCGGCAAAATAGTAAAGGCTCATCATGGCAGAAAATTGCGACACCCCGCGTCCGAACCGGCAGCGATCTGGGTAAAAAGCTGCTGGAAGCAGCCG<690
 845<AGTCTGCGGCAAAATAGTAAAGGCTCATCATGGCAGAAAATTGCGACACCCCGCGTCCGAACCGGCAGCGATCTGGGTAAAAAGCTGCTGGAAGCAGCCG<746
 847<AGTCTGCGGCAAAATAGTAAAGGCTCATCATGGCAGAAAATTGCGACACCCCGCGTCCGAACCGGCAGCGATCTGGGTAAAAAGCTGCTGGAAGCAGCCG<748

              *         *         *         *         *         *         *         *         *
5286>cggccggccaagatgatgaggtgcgtattctgatggcgaatggggccgatgttaacgcaaccgacgacgatggcctgactccgct--------------->5370
 688<CGGCCGGCCAAGATGATGAGGTGCGTATTCTGATGGCGAATGGGGCCGATGTTAACGCAACCGACGACGATGGCCTGACTCCGCT---------------<604
 688<CGGCCGGCCAAGATGATGAGGTGCGTATTCTGATGGCGAATGGGGCCGATGTTAACGCAACCGACGACAATGGCCTGACTCCGCT---------------<604
 689<CGGCCGGCCAAGATGATGAGGTGCGTATTCTGATGGCGAATGGGGCCGATGTTAACGCAACCGACGACAATGGCCTGACTCCGCT---------------<605
 689<CGGCCGGCCAAGATGATGAGGTGCGTATTCTGATGGCGAATGGGGCCGATGTTAACGCAACCGACGACGATGGCCTGACTCCGCT---------------<605
 745<CGGCCGGCCAAGATGATGAGGTGCGTATTCTGATGGCGAATGGGGCCGATGTTAACGCAACCGACGACGATGGCCTGACTCCGCTGCACCTGGCGAATGG<646
 747<CGGCCGGCCAAGATGATGAGGTGCGTATTCTGATGGCGAATGGGGCCGATGTTAACGCAACCGACGACGATGGCCTGACTCCGCTGCACCTGGCGGAATG<648

                                                          *         *         *         *         *
5371>-------------------------------------------gcacctggcggctgcaaacgggcaactggaaatcgtagaggtactgctgaaaaatgg>5427
 603<-------------------------------------------GCACCTGGCGGCTGCAAACGGGCAACTGGAAATCGTAGAGGTACTGCTGAAAAATGG<547
 603<-------------------------------------------GCACCTGGCGGCTGCAAACGGGCAACTGGAAATCGTAGAGGTACTGCTGAAAAATGG<547
 604<-------------------------------------------GCACCTGGCGGCTGCAAACGGGCAACTGGAAATCGTAGAGGTACTGCTGAAAAATGG<548
 604<-------------------------------------------GCACCTGGCGGCTGCAAACGGGCAACTGGAAATCGTAGAGGTACTGCTGAAAAATGG<548
 645<GGCCGATGTTAACGCAACCGACGACGATGGCCTGACTCCGCT-GCACCTGGCGGCTGCAAACGGGCAACTGGAAATCGTAGAGGTACTGCTGAAAAATGG<547
 647<GGGCCGATGTTAACGCAACCGACGACGATGGCCTGACTCCGCTGCACCTGGCGGCTGCAAACGGGCAACTGGAAATCGTAGAGGTACTGCTGAAAAATGG<548

        *         *         *         *         *         *         *         *         *         *
```

```
5428>cgccgatgtgaacgcttctgatagtgcgggtattactccgctgcacctggccgcttatgacggccatctggagattgtcgaagtcctgctgaagcacggg>5527
 546<CGCCGATGTGAACGCTTCTGATAGTGCGGGTATTACTCCGCTGCACCTGGCCGCTTATGACGGCCATCTGGAGATTGTCGAAGTCCTGCTGAAGCACGGG<447
 546<CGCCGATGTGAACGCTTCTGATAGTGCGGGTATTACTCCGCTGCACCTGGCCGCTTATGACGGCCATCTGGAGATTGTCGAAGTCCTGCTGAAGCACGGG<447
 547<CGCCGATGTGAACGCTTCTGATAGTGCGGGTATTACTCCGCTGCACCTGGCCGCTTATGACGGCCATCTGGAGATTGTCGAAGTCCTGCTGAAGCACGGG<448
 547<CGCCGATGTGAACGCTTCTGATAGTGCGGGTATTACTCCGCTGCACCTGGCCGCTTATGACGGCCATCTGGAGATTGTCGAAGTCCTGCTGAAGCACGGG<448
 546<CGCCGATGTGAACGCTTCTGATAGTGCGGGTATTACTCCGCTGCACCTGGCCGCTTATGACGGCCATCTGGAGATTGTCGAAGTCCTGCTGAAGCACGGG<447
 547<CGCCGATGTGAACGCTTCTGATAGTGCGGGTATTACTCCGCTGCACCTGGCCGCTTATGACGGCCATCTGGAGATTGTCGAAGTCCTGCTGAAGCACGGG<448

       *         *         *         *         *         *         *         *         *         *
5528>gctgacgttaatgcgtacgaccgcgccgggtggacaccgctgcacctagcagcgctgagtggccaactggagattgtggaagttctgctgaaacacggcg>5627
 446<GCTGACGTTAATGCGTACGACCGCGCCGGGTGGACACCGCTGCACCTAGCAGCGCTGAGTGGCCAACTGGAGATTGTGGAAGTTCTGCTGAAACACGGCG<347
 446<GCTGACGTTAATGCGTACGACCGCGCCGGGTGGACACCGCTGCACCTAGCAGCGCTGAGTGGCCAACTGGAGATTGTGGAAGTTCTGCTGAAACACGGCG<347
 447<GCTGACGTTAATGCGTACGACCGCGCCGGGTGGACACCGCTGCACCTAGCAGCGCTGAGTGGCCAACTGGAGATTGTGGAAGTTCTGCTGAAACACGGCG<348
 447<GCTGACGTTAATGCGTACGACCGCGCCGGGTGGACACCGCTGCACCTAGCAGCGCTGAGTGGCCAACTGGAGATTGTGGAAGTTCTGCTGAAACACGGCG<348
 446<GCTGACGTTAATGCGTACGACCGCGCCGGGTGGACACCGCTGCACCTAGCAGCGCTGAGTGGCCAACTGGAGATTGTGGAAGTTCTGCTGAAACACGGCG<347
 447<GCTGACGTTAATGCGTACGACCGCGCCGGGTGGACACCGCTGCACCTAGCAGCGCTGAGTGGCCAACTGGAGATTGTGGAAGTTCTGCTGAAACACGGCG<348

       *         *         *         *         *         *         *         *         *         *
                                                                                                     H  H
                                                                                                    CACca
5628>cagatgtcaacgcccaagacgcactgggcctgaccgcgtttgatatctcgattaatcaaggtcaggaagatctggcagagatcctgcaactcgagcacca>5727
 346<CAGATGTCAACGCCCAAGACGCACTGGGCCTGACCGCGTTTGATATCTCGATTAATCAAGGTCAGGAAGATCTGGCAGAGATCCTGCAACTCGAGCACCA<247
 346<CAGATGTCAACGCCCAAGACGCACTGGGCCTGACCGCGTTTGATATCTCGATTAATCAAGGTCAGGAAGATCTGGCAGAGATCCTGCAACTCGAGCACCA<247
 347<CAGATGTCAACGCCCAAGACGCACTGGGCCTGACCGCGTTTGATATCTCGATTAATCAAGGTCAGGAAGATCTGGCAGAGATCCTGCAACTCGAGCACCA<248
 347<CAGATGTCAACGCCCAAGACGCACTGGGCCTGACCGCGTTTGATATCTCGATTAATCAAGGTCAGGAAGATCTGGCAGAGATCCTGCAACTCGAGCACCA<248
 346<CAGATGTCAACGCCCAAGACGCACTGGGCCTGACCGCGTTTGATATCTCGATTAATCAAGGTCAGGAAGATCTGGCAGAGATCCTGCAACTCGAGCACCA<247
 347<CAGATGTCAACGCCCAAGACGCACTGGGCCTGACCGCGTTTGATATCTCGATTAATCAAGGTCAGGAAGATCTGGCAGAGATCCTGCAACTCGAGCACCA<248

       *         *
       H  H  H  H      ----------------------------------------------------------------------------------
     cCACcacCACcac     ----------------------------------------------------------------------------------
5728>ccaccaccaccactgaca---------------------------------------------------------------------------------->5745
 246<CCACCACCACCACTGACACCCAGCTTTCTTGTCCACCACCACTGACACCCAGCTTTCTTGTCCACCACCACTGACACCCAGCTTTCTTGTCCACCACCAC<147
 246<CCACCACCACCACTGACACCCAGCTTTCTTGTCCACCACCACTGACACCCAGCTTTCTTGTCCACCACCACTGACACCCAGCTTTCTTGTCCACCACCAC<147
 247<CCACCACCACCACTGACACCCAGCTTTCTTGTCCACCACCACTGACACCCAGCTTTCTTGTCCACCACCACTGACACCCAGCTTTCTTGTCCACCACCAC<148
 247<CCACCACCACCACTGACACCCAGCTTTCTTGTCCACCACCACTGACACCCAGCTTTCTTGTCCACCACCACTGACACCCAGCTTTCTTGTCCACCACCAC<148
 246<CCACCACCACCACTGACACCCAGCTTTCTTGTCCACCACCACTGACACCCAGCTTTCTTGTCCACCACCACTGACACCCAGCTTTCTTGTCCACCACCAC<147
 247<CCACCACCACCACTGACACCCAGCTTTCTTGTCCACCACCACTGACACCCAGCTTTCTTGTCCACCACCACTGACACCCAGCTTTCTTGTCCACCACCAC<148
```

```
                                                                                    *         *         *         *
5746>---------------------------------------------------------------cccagctttcttgtacaaagtggttcttgtacaaagt>5782
 146<TGACACCCAGCTTTCTTGTCCACCACCACTGACACCCAGCTTTCTTGTCCACCACCACTGACACCCAGCTTTCTTGT----------------ACAAAGT<63
 146<TGACACCCAGCTTTCTTGTCCACCACCACTGACACCCAGCTTTCTTGTCCACCACCACTGACACCCAGCTTTCTTGT----------------ACAAAGT<63
 147<TGACACCCAGCTTTCTTGTCCACCACCACTGACACCCAGCTTTCTTGTCCACCACCACTGACACCCAGCTTTCTTGT----------------ACAAAGT<64
 147<TGACACCCAGCTTTCTTGTCCACCACCACTGACACCCAGCTTTCTTGTCCACCACCACTGACACCCAGCTTTCTTGT----------------ACAAAGT<64
 146<TGACACCCAGCTTTCTTGTCCACCACCACTGACACCCAGCTTTCTTGTCCACCACCACTGACACCCAGCTTTCTTGT----------------ACAAAGT<63
 147<TGACACCCAGCTTTCTTGTCCACCACCACTGACACCCAGCTTTCTTGTCCACCACCACTGACACCCAGCTTTCTTGT----------------ACAAAGT<64

              *         *         *         *         *         *         *         *         *         *
5783>ggtgatgggctgcaggaattcgatatcaagcttatcgataccgtcgacctcgagtcatgtaattagttatgtcacgcttacattcacgccctccccccac>5882
  62<GGTGATGGGCTGCAGGAATTCGATTTCAAGCTTATCGATACCGTCGACCTCGAGTCA-------------------------------------------<6
  62<GGTGATGGGCTGCAGGAATTCGATTTCAAGCTTATCGATACCGTCGACCTCGAGTCA-------------------------------------------<6
  63<GGTGATGGGCTGCAGGAATTCGATTTCAAGCTTATCGATACCGTCGACCTCGAGTCA-------------------------------------------<7
  63<GGTGATGGGCTGCAGGAATTCGATTTCAAGCTTATCGATACCGTCGACCTCGAGTCA-------------------------------------------<7
  62<GGTGATGGGCTGCAGGAATTCGAT-TCAAGCTTATCGATACCGTCGACCTCGAGTCA-------------------------------------------<7
  63<GGTGATGGGCTGCAGGAATTCGATTTCAAGCTTATCGATACCGTCGACCTCGAGTCA-------------------------------------------<7

              *         *         *         *         *         *         *         *         *         *
5883>atccgctctaaccgaaaaggaaggagttagacaacctgaagtctaggtccctatttattttttatagttatgttagtattaagaacgttatttatattt>5982
   6<----------------------------------------------------------------------------------------------------<6
   6<----------------------------------------------------------------------------------------------------<6
   7<----------------------------------------------------------------------------------------------------<7
   7<----------------------------------------------------------------------------------------------------<7
   7<----------------------------------------------------------------------------------------------------<7
   7<----------------------------------------------------------------------------------------------------<7

              *         *         *         *         *         *         *         *         *         *
5983>caaattttcttttttttttctgtacagacgcgtgtacgcatgtaacattatactgaaaaccttgcttgagaaggttttgggacgctcgaaggctttaattt>6082
   6<----------------------------------------------------------------------------------------------------<6
   6<----------------------------------------------------------------------------------------------------<6
   7<----------------------------------------------------------------------------------------------------<7
   7<----------------------------------------------------------------------------------------------------<7
   7<----------------------------------------------------------------------------------------------------<7
   7<----------------------------------------------------------------------------------------------------<7

              *         *         *         *         *         *         *         *         *         *
6083>gtgacaccgattatttaaagctgcagcatacgatatatatacatgtgtatatatgtatacctatgaatgtcagtaagtatgtatacgaacagtatgatac>6182
   5<-------------------------------------------------------------------------------------------GTATG~~~~<1
```

```
   6<----------------------------------------------------------------------------------------------------<6
   7<----------------------------------------------------------------------------------------------------<7
   7<----------------------------------------------------------------------------------------------------<7
   6<------------------------------------------------------------------------GTAAGT~~~~~~~~~~~~~~~~~~~~~~<1
   7<----------------------------------------------------------------------------------------------------<7

            *         *         *         *         *         *         *         *         *         *
6183>tgaagatgacaaggtaatgcatcattctatacgtgtcattctgaacgaggcgcgctttccttttttctttttgctttttcttttttttttctcttgaactc>6282
   1<~~~~~~~~~~~~~~~~~~~~~~~~~~~~~~~~~~~~~~~~~~~~~~~~~~~~~~~~~~~~~~~~~~~~~~~~~~~~~~~~~~~~~~~~~~~~~~~~~~~<1
   6<----------------------------------------------------------------------------------------------------<6
   7<----------------------------------------------------------------------------------------------------<7
   7<----------------------------------------------------------------------------------------------------<7
   1<~~~~~~~~~~~~~~~~~~~~~~~~~~~~~~~~~~~~~~~~~~~~~~~~~~~~~~~~~~~~~~~~~~~~~~~~~~~~~~~~~~~~~~~~~~~~~~~~~~~<1
   7<----------------------------------------------------------------------------------------------------<7

            *         *         *         *         *         *         *         *         *         *
6283>gagaaaaaaaatataaaagagatggaggaacgggaaaaagttagttgtggtgataggtggcaagtggtattccgtaagaacaacaagaaaagcatttcat>6382
   1<~~~~~~~~~~~~~~~~~~~~~~~~~~~~~~~~~~~~~~~~~~~~~~~~~~~~~~~~~~~~~~~~~~~~~~~~~~~~~~~~~~~~~~~~~~~~~~~~~~~<1
   5<-------------------------------------------------------------------------GTAAG~~~~~~~~~~~~~~~~~~~~~~<1
   7<----------------------------------------------------------------------------------------------------<7
   7<----------------------------------------------------------------------------------------------------<7
   1<~~~~~~~~~~~~~~~~~~~~~~~~~~~~~~~~~~~~~~~~~~~~~~~~~~~~~~~~~~~~~~~~~~~~~~~~~~~~~~~~~~~~~~~~~~~~~~~~~~~<1
   7<----------------------------------------------------------------------------------------------------<7

            *         *         *         *         *         *         *         *         *         *
6383>attatggctgaactgagcgaacaagtgcaaaatttaagcatcaacgacaacaacgagaatggttatgttcctcctcacttaagaggaaaaccaagaagtg>6482
   1<~~~~~~~~~~~~~~~~~~~~~~~~~~~~~~~~~~~~~~~~~~~~~~~~~~~~~~~~~~~~~~~~~~~~~~~~~~~~~~~~~~~~~~~~~~~~~~~~~~~<1
   1<~~~~~~~~~~~~~~~~~~~~~~~~~~~~~~~~~~~~~~~~~~~~~~~~~~~~~~~~~~~~~~~~~~~~~~~~~~~~~~~~~~~~~~~~~~~~~~~~~~~<1
   7<----------------------------------------------------------------------------------------------------<7
   7<----------------------------------------------------------------------------------------------------<7
   1<~~~~~~~~~~~~~~~~~~~~~~~~~~~~~~~~~~~~~~~~~~~~~~~~~~~~~~~~~~~~~~~~~~~~~~~~~~~~~~~~~~~~~~~~~~~~~~~~~~~<1
   7<----------------------------------------------------------------------------------------------------<7

            *         *         *         *         *         *         *         *         *         *
6483>ccagaaataacagtagcaactacaataacaacaacggcggctacaacggtggccgtggcggtggcagcttctttagcaacaaccgtcgtggtggttacgg>6582
   1<~~~~~~~~~~~~~~~~~~~~~~~~~~~~~~~~~~~~~~~~~~~~~~~~~~~~~~~~~~~~~~~~~~~~~~~~~~~~~~~~~~~~~~~~~~~~~~~~~~~<1
   1<~~~~~~~~~~~~~~~~~~~~~~~~~~~~~~~~~~~~~~~~~~~~~~~~~~~~~~~~~~~~~~~~~~~~~~~~~~~~~~~~~~~~~~~~~~~~~~~~~~~<1
   7<----------------------------------------------------------------------------------------------------<7
   7<----------------------------------------------------------------------------------------------------<7
```

```
   1<~~~~~~~~~~~~~~~~~~~~~~~~~~~~~~~~~~~~~~~~~~~~~~~~~~~~~~~~~~~~~~~~~~~~~~~~~~~~~~~~~~~~~~~~~~~~~~~~~~~~<1
   7<----------------------------------------------------------------------------------------------------<7

              *         *         *         *         *         *         *         *         *         *
6583>caacggtggtttcttcggtggaaacaacggtggcagcagatctaacggccgttctggtggtagatggatcgatggcaaacatgtcccagctccaagaaac>6682
   1<~~~~~~~~~~~~~~~~~~~~~~~~~~~~~~~~~~~~~~~~~~~~~~~~~~~~~~~~~~~~~~~~~~~~~~~~~~~~~~~~~~~~~~~~~~~~~~~~~~~~<1
   1<~~~~~~~~~~~~~~~~~~~~~~~~~~~~~~~~~~~~~~~~~~~~~~~~~~~~~~~~~~~~~~~~~~~~~~~~~~~~~~~~~~~~~~~~~~~~~~~~~~~~<1
   7<----------------------------------------------------------------------------------------------------<7
   7<----------------------------------------------------------------------------------------------------<7
   1<~~~~~~~~~~~~~~~~~~~~~~~~~~~~~~~~~~~~~~~~~~~~~~~~~~~~~~~~~~~~~~~~~~~~~~~~~~~~~~~~~~~~~~~~~~~~~~~~~~~~<1
   7<----------------------------------------------------------------------------------------------------<7

              *         *         *         *         *         *         *         *         *         *
6683>gaaaaggccgagatcgccatatttggtgtccccgaggatccaaatttccaatcttctggtattaacttcgataactacgatgatattccagtggacgcct>6782
   1<~~~~~~~~~~~~~~~~~~~~~~~~~~~~~~~~~~~~~~~~~~~~~~~~~~~~~~~~~~~~~~~~~~~~~~~~~~~~~~~~~~~~~~~~~~~~~~~~~~~~<1
   1<~~~~~~~~~~~~~~~~~~~~~~~~~~~~~~~~~~~~~~~~~~~~~~~~~~~~~~~~~~~~~~~~~~~~~~~~~~~~~~~~~~~~~~~~~~~~~~~~~~~~<1
   7<----------------------------------------------------------------------------------------------------<7
   7<----------------------------------------------------------------------------------------------------<7
   1<~~~~~~~~~~~~~~~~~~~~~~~~~~~~~~~~~~~~~~~~~~~~~~~~~~~~~~~~~~~~~~~~~~~~~~~~~~~~~~~~~~~~~~~~~~~~~~~~~~~~<1
   7<----------------------------------------------------------------------------------------------------<7

              *         *         *         *         *         *         *         *         *         *
6783>ctggtaaggatgttcctgaaccaatcacagaatttacctcacctccattggacggattgttattggaaaacatcaaattggcccgtttcaccaagccaac>6882
   1<~~~~~~~~~~~~~~~~~~~~~~~~~~~~~~~~~~~~~~~~~~~~~~~~~~~~~~~~~~~~~~~~~~~~~~~~~~~~~~~~~~~~~~~~~~~~~~~~~~~~<1
   1<~~~~~~~~~~~~~~~~~~~~~~~~~~~~~~~~~~~~~~~~~~~~~~~~~~~~~~~~~~~~~~~~~~~~~~~~~~~~~~~~~~~~~~~~~~~~~~~~~~~~<1
   6<---GTAAGG~~~~~~~~~~~~~~~~~~~~~~~~~~~~~~~~~~~~~~~~~~~~~~~~~~~~~~~~~~~~~~~~~~~~~~~~~~~~~~~~~~~~~~~~~~~<1
   7<----------------------------------------------------------------------------------------------------<7
   1<~~~~~~~~~~~~~~~~~~~~~~~~~~~~~~~~~~~~~~~~~~~~~~~~~~~~~~~~~~~~~~~~~~~~~~~~~~~~~~~~~~~~~~~~~~~~~~~~~~~~<1
   7<----------------------------------------------------------------------------------------------------<7

              *         *         *         *         *         *         *         *         *         *
6883>acctgtgcaaaaatactccgtccctatcgttgccaacggcagagatttgatggcctgtgcgcagaccggttctggtaagactggtgggtttttattccca>6982
   1<~~~~~~~~~~~~~~~~~~~~~~~~~~~~~~~~~~~~~~~~~~~~~~~~~~~~~~~~~~~~~~~~~~~~~~~~~~~~~~~~~~~~~~~~~~~~~~~~~~~~<1
   1<~~~~~~~~~~~~~~~~~~~~~~~~~~~~~~~~~~~~~~~~~~~~~~~~~~~~~~~~~~~~~~~~~~~~~~~~~~~~~~~~~~~~~~~~~~~~~~~~~~~~<1
   1<~~~~~~~~~~~~~~~~~~~~~~~~~~~~~~~~~~~~~~~~~~~~~~~~~~~~~~~~~~~~~~~~~~~~~~~~~~~~~~~~~~~~~~~~~~~~~~~~~~~~<1
   7<----------------------------------------------------------------------------------------------------<7
   1<~~~~~~~~~~~~~~~~~~~~~~~~~~~~~~~~~~~~~~~~~~~~~~~~~~~~~~~~~~~~~~~~~~~~~~~~~~~~~~~~~~~~~~~~~~~~~~~~~~~~<1
   7<----------------------------------------------------------------------------------------------------<7
```

```
              *         *         *         *         *         *         *         *         *         *
6983>gtgttgtccgaatcatttaagactggaccatctcctcaaccagagtctcaaggctccttttaccaaagaaaggcctacccaactgctgtcattatggctc>7082
   1<~~~~~~~~~~~~~~~~~~~~~~~~~~~~~~~~~~~~~~~~~~~~~~~~~~~~~~~~~~~~~~~~~~~~~~~~~~~~~~~~~~~~~~~~~~~~~~~~~~~~<1
   1<~~~~~~~~~~~~~~~~~~~~~~~~~~~~~~~~~~~~~~~~~~~~~~~~~~~~~~~~~~~~~~~~~~~~~~~~~~~~~~~~~~~~~~~~~~~~~~~~~~~~<1
   1<~~~~~~~~~~~~~~~~~~~~~~~~~~~~~~~~~~~~~~~~~~~~~~~~~~~~~~~~~~~~~~~~~~~~~~~~~~~~~~~~~~~~~~~~~~~~~~~~~~~~<1
   7<----------------------------------------------------------------------------------------------------<7
   1<~~~~~~~~~~~~~~~~~~~~~~~~~~~~~~~~~~~~~~~~~~~~~~~~~~~~~~~~~~~~~~~~~~~~~~~~~~~~~~~~~~~~~~~~~~~~~~~~~~~~<1
   7<----------------------------------------------------------------------------------------------------<7

              *         *         *         *         *         *         *         *         *         *
7083>cagtttaaaccatggtcatagctgtttcctgtgtgaaattgttatccgctcacaattccacacaacataggagccggaagcataaagtgtaaagcctggg>7182
   1<~~~~~~~~~~~~~~~~~~~~~~~~~~~~~~~~~~~~~~~~~~~~~~~~~~~~~~~~~~~~~~~~~~~~~~~~~~~~~~~~~~~~~~~~~~~~~~~~~~~~<1
   1<~~~~~~~~~~~~~~~~~~~~~~~~~~~~~~~~~~~~~~~~~~~~~~~~~~~~~~~~~~~~~~~~~~~~~~~~~~~~~~~~~~~~~~~~~~~~~~~~~~~~<1
   1<~~~~~~~~~~~~~~~~~~~~~~~~~~~~~~~~~~~~~~~~~~~~~~~~~~~~~~~~~~~~~~~~~~~~~~~~~~~~~~~~~~~~~~~~~~~~~~~~~~~~<1
   7<----------------------------------------------------------------------------------------------------<7
   1<~~~~~~~~~~~~~~~~~~~~~~~~~~~~~~~~~~~~~~~~~~~~~~~~~~~~~~~~~~~~~~~~~~~~~~~~~~~~~~~~~~~~~~~~~~~~~~~~~~~~<1
   7<----------------------------------------------------------------------------------------------------<7

              *         *         *         *         *         *         *         *         *         *
7183>gtgcctaatgagtgaggtaactcacattaattgcgttgcgctcactgcccgctttccagtcgggaaacctgtcgtgccagctgcattaatgaatcggcca>7282
   1<~~~~~~~~~~~~~~~~~~~~~~~~~~~~~~~~~~~~~~~~~~~~~~~~~~~~~~~~~~~~~~~~~~~~~~~~~~~~~~~~~~~~~~~~~~~~~~~~~~~~<1
   1<~~~~~~~~~~~~~~~~~~~~~~~~~~~~~~~~~~~~~~~~~~~~~~~~~~~~~~~~~~~~~~~~~~~~~~~~~~~~~~~~~~~~~~~~~~~~~~~~~~~~<1
   1<~~~~~~~~~~~~~~~~~~~~~~~~~~~~~~~~~~~~~~~~~~~~~~~~~~~~~~~~~~~~~~~~~~~~~~~~~~~~~~~~~~~~~~~~~~~~~~~~~~~~<1
   7<----------------------------------------------------------------------------------------------------<7
   1<~~~~~~~~~~~~~~~~~~~~~~~~~~~~~~~~~~~~~~~~~~~~~~~~~~~~~~~~~~~~~~~~~~~~~~~~~~~~~~~~~~~~~~~~~~~~~~~~~~~~<1
   7<----------------------------------------------------------------------------------------------------<7

              *         *         *         *         *         *         *         *         *         *
7283>acgcgcggggagaggcggtttgcgtattgggcgctcttccgcttcctcgctcactgactcgctgcgctcggtcgttcggctgcggcgagcggtatcagct>7382
   1<~~~~~~~~~~~~~~~~~~~~~~~~~~~~~~~~~~~~~~~~~~~~~~~~~~~~~~~~~~~~~~~~~~~~~~~~~~~~~~~~~~~~~~~~~~~~~~~~~~~~<1
   1<~~~~~~~~~~~~~~~~~~~~~~~~~~~~~~~~~~~~~~~~~~~~~~~~~~~~~~~~~~~~~~~~~~~~~~~~~~~~~~~~~~~~~~~~~~~~~~~~~~~~<1
   1<~~~~~~~~~~~~~~~~~~~~~~~~~~~~~~~~~~~~~~~~~~~~~~~~~~~~~~~~~~~~~~~~~~~~~~~~~~~~~~~~~~~~~~~~~~~~~~~~~~~~<1
   7<----------------------------------------------------------------------------------------------------<7
   1<~~~~~~~~~~~~~~~~~~~~~~~~~~~~~~~~~~~~~~~~~~~~~~~~~~~~~~~~~~~~~~~~~~~~~~~~~~~~~~~~~~~~~~~~~~~~~~~~~~~~<1
   7<----------------------------------------------------------------------------------------------------<7

              *         *         *         *         *         *         *         *         *         *
7383>cactcaaaggcggtaatacggttatccacagaatcaggggataacgcaggaaagaacatgtgagcaaaaggccagcaaaaggccaggaaccgtaaaaagg>7482
```

```
   1<~~~~~~~~~~~~~~~~~~~~~~~~~~~~~~~~~~~~~~~~~~~~~~~~~~~~~~~~~~~~~~~~~~~~~~~~~~~~~~~~~~~~~~~~~~~~~~~~<1
   1<~~~~~~~~~~~~~~~~~~~~~~~~~~~~~~~~~~~~~~~~~~~~~~~~~~~~~~~~~~~~~~~~~~~~~~~~~~~~~~~~~~~~~~~~~~~~~~~~<1
   1<~~~~~~~~~~~~~~~~~~~~~~~~~~~~~~~~~~~~~~~~~~~~~~~~~~~~~~~~~~~~~~~~~~~~~~~~~~~~~~~~~~~~~~~~~~~~~~~~<1
   7<----------------------------------------------------------------------------------------------------<7
   1<~~~~~~~~~~~~~~~~~~~~~~~~~~~~~~~~~~~~~~~~~~~~~~~~~~~~~~~~~~~~~~~~~~~~~~~~~~~~~~~~~~~~~~~~~~~~~~~~<1
   7<----------------------------------------------------------------------------------------------------<7

              *         *         *         *         *         *         *         *         *         *
7483>ccgcgttgctggcgtttttccataggctcggcccccctgacgagcatcacaaaaatcgacgctcaagtcagaggtggcgaaacccgacaggactataaag>7582
   1<~~~~~~~~~~~~~~~~~~~~~~~~~~~~~~~~~~~~~~~~~~~~~~~~~~~~~~~~~~~~~~~~~~~~~~~~~~~~~~~~~~~~~~~~~~~~~~~~<1
   1<~~~~~~~~~~~~~~~~~~~~~~~~~~~~~~~~~~~~~~~~~~~~~~~~~~~~~~~~~~~~~~~~~~~~~~~~~~~~~~~~~~~~~~~~~~~~~~~~<1
   1<~~~~~~~~~~~~~~~~~~~~~~~~~~~~~~~~~~~~~~~~~~~~~~~~~~~~~~~~~~~~~~~~~~~~~~~~~~~~~~~~~~~~~~~~~~~~~~~~<1
   7<----------------------------------------------------------------------------------------------------<7
   1<~~~~~~~~~~~~~~~~~~~~~~~~~~~~~~~~~~~~~~~~~~~~~~~~~~~~~~~~~~~~~~~~~~~~~~~~~~~~~~~~~~~~~~~~~~~~~~~~<1
   7<----------------------------------------------------------------------------------------------------<7

              *         *         *         *         *         *         *         *         *         *
7583>ataccaggcgttcccccctggaagctccctcgtgcgctctcctgttccgaccctgccgcttaccggatacctgtccgcctttctcccttcgggaagcgtg>7682
   1<~~~~~~~~~~~~~~~~~~~~~~~~~~~~~~~~~~~~~~~~~~~~~~~~~~~~~~~~~~~~~~~~~~~~~~~~~~~~~~~~~~~~~~~~~~~~~~~~<1
   1<~~~~~~~~~~~~~~~~~~~~~~~~~~~~~~~~~~~~~~~~~~~~~~~~~~~~~~~~~~~~~~~~~~~~~~~~~~~~~~~~~~~~~~~~~~~~~~~~<1
   1<~~~~~~~~~~~~~~~~~~~~~~~~~~~~~~~~~~~~~~~~~~~~~~~~~~~~~~~~~~~~~~~~~~~~~~~~~~~~~~~~~~~~~~~~~~~~~~~~<1
   7<----------------------------------------------------------------------------------------------------<7
   1<~~~~~~~~~~~~~~~~~~~~~~~~~~~~~~~~~~~~~~~~~~~~~~~~~~~~~~~~~~~~~~~~~~~~~~~~~~~~~~~~~~~~~~~~~~~~~~~~<1
   7<----------------------------------------------------------------------------------------------------<7

              *         *         *         *         *         *         *         *         *         *
7683>gcgctttctcaatgctcacgctgtaggtatctcagttcggtgtaggtcgttcgctccaagctgggctgtgtgcacgaaccccccgttcagcccgaccgct>7782
   1<~~~~~~~~~~~~~~~~~~~~~~~~~~~~~~~~~~~~~~~~~~~~~~~~~~~~~~~~~~~~~~~~~~~~~~~~~~~~~~~~~~~~~~~~~~~~~~~~<1
   1<~~~~~~~~~~~~~~~~~~~~~~~~~~~~~~~~~~~~~~~~~~~~~~~~~~~~~~~~~~~~~~~~~~~~~~~~~~~~~~~~~~~~~~~~~~~~~~~~<1
   1<~~~~~~~~~~~~~~~~~~~~~~~~~~~~~~~~~~~~~~~~~~~~~~~~~~~~~~~~~~~~~~~~~~~~~~~~~~~~~~~~~~~~~~~~~~~~~~~~<1
   7<----------------------------------------------------------------------------------------------------<7
   1<~~~~~~~~~~~~~~~~~~~~~~~~~~~~~~~~~~~~~~~~~~~~~~~~~~~~~~~~~~~~~~~~~~~~~~~~~~~~~~~~~~~~~~~~~~~~~~~~<1
   7<----------------------------------------------------------------------------------------------------<7

              *         *         *         *         *         *         *         *         *         *
7783>gcgccttatccggtaactatcgtcttgagtccaacccggtaagacacgacttatcgccactggcagcagccactggtaacaggattagcagagcgaggta>7882
   1<~~~~~~~~~~~~~~~~~~~~~~~~~~~~~~~~~~~~~~~~~~~~~~~~~~~~~~~~~~~~~~~~~~~~~~~~~~~~~~~~~~~~~~~~~~~~~~~~<1
   1<~~~~~~~~~~~~~~~~~~~~~~~~~~~~~~~~~~~~~~~~~~~~~~~~~~~~~~~~~~~~~~~~~~~~~~~~~~~~~~~~~~~~~~~~~~~~~~~~<1
   1<~~~~~~~~~~~~~~~~~~~~~~~~~~~~~~~~~~~~~~~~~~~~~~~~~~~~~~~~~~~~~~~~~~~~~~~~~~~~~~~~~~~~~~~~~~~~~~~~<1
```

```
    7<----------------------------------------------------------------------------------------------------<7
    1<~~~~~~~~~~~~~~~~~~~~~~~~~~~~~~~~~~~~~~~~~~~~~~~~~~~~~~~~~~~~~~~~~~~~~~~~~~~~~~~~~~~~~~~~~~~~~~~~~~~~<1
    7<----------------------------------------------------------------------------------------------------<7

              *         *         *         *         *         *         *         *         *         *
 7883>tgtaggcggtgctacagagttcttgaagtggtggcctaactacggctacactagaaggacagtatttggtatctgcgctctgctgaagccagttaccttc>7982
    1<~~~~~~~~~~~~~~~~~~~~~~~~~~~~~~~~~~~~~~~~~~~~~~~~~~~~~~~~~~~~~~~~~~~~~~~~~~~~~~~~~~~~~~~~~~~~~~~~~~~~<1
    1<~~~~~~~~~~~~~~~~~~~~~~~~~~~~~~~~~~~~~~~~~~~~~~~~~~~~~~~~~~~~~~~~~~~~~~~~~~~~~~~~~~~~~~~~~~~~~~~~~~~~<1
    1<~~~~~~~~~~~~~~~~~~~~~~~~~~~~~~~~~~~~~~~~~~~~~~~~~~~~~~~~~~~~~~~~~~~~~~~~~~~~~~~~~~~~~~~~~~~~~~~~~~~~<1
    7<----------------------------------------------------------------------------------------------------<7
    1<~~~~~~~~~~~~~~~~~~~~~~~~~~~~~~~~~~~~~~~~~~~~~~~~~~~~~~~~~~~~~~~~~~~~~~~~~~~~~~~~~~~~~~~~~~~~~~~~~~~~<1
    7<----------------------------------------------------------------------------------------------------<7

              *         *         *         *         *         *         *         *         *         *
 7983>ggaaaaagagttggtagctcttgatccggcaaacaaaccaccgctggtagcggtggtttttttgtttgcaagcagcagattacgcgcagaaaaaaaggat>8082
    1<~~~~~~~~~~~~~~~~~~~~~~~~~~~~~~~~~~~~~~~~~~~~~~~~~~~~~~~~~~~~~~~~~~~~~~~~~~~~~~~~~~~~~~~~~~~~~~~~~~~~<1
    1<~~~~~~~~~~~~~~~~~~~~~~~~~~~~~~~~~~~~~~~~~~~~~~~~~~~~~~~~~~~~~~~~~~~~~~~~~~~~~~~~~~~~~~~~~~~~~~~~~~~~<1
    1<~~~~~~~~~~~~~~~~~~~~~~~~~~~~~~~~~~~~~~~~~~~~~~~~~~~~~~~~~~~~~~~~~~~~~~~~~~~~~~~~~~~~~~~~~~~~~~~~~~~~<1
    7<----------------------------------------------------------------------------------------------------<7
    1<~~~~~~~~~~~~~~~~~~~~~~~~~~~~~~~~~~~~~~~~~~~~~~~~~~~~~~~~~~~~~~~~~~~~~~~~~~~~~~~~~~~~~~~~~~~~~~~~~~~~<1
    7<----------------------------------------------------------------------------------------------------<7

              *         *         *         *         *         *         *         *         *         *
 8083>ctcaagaagatcctttgatcttttctacggggtctgacgctcagtggaacgaaaactcacgttaagggattttggtcatgagattatcaaaaaggatctt>8182
    1<~~~~~~~~~~~~~~~~~~~~~~~~~~~~~~~~~~~~~~~~~~~~~~~~~~~~~~~~~~~~~~~~~~~~~~~~~~~~~~~~~~~~~~~~~~~~~~~~~~~~<1
    1<~~~~~~~~~~~~~~~~~~~~~~~~~~~~~~~~~~~~~~~~~~~~~~~~~~~~~~~~~~~~~~~~~~~~~~~~~~~~~~~~~~~~~~~~~~~~~~~~~~~~<1
    1<~~~~~~~~~~~~~~~~~~~~~~~~~~~~~~~~~~~~~~~~~~~~~~~~~~~~~~~~~~~~~~~~~~~~~~~~~~~~~~~~~~~~~~~~~~~~~~~~~~~~<1
    7<----------------------------------------------------------------------------------------------------<7
    1<~~~~~~~~~~~~~~~~~~~~~~~~~~~~~~~~~~~~~~~~~~~~~~~~~~~~~~~~~~~~~~~~~~~~~~~~~~~~~~~~~~~~~~~~~~~~~~~~~~~~<1
    7<----------------------------------------------------------------------------------------------------<7

              *         *         *         *         *         *         *         *         *         *
                                                                                   *  W  H  K  I  L  S  A
                                                                                  TTAccaATGcttAATcagTGAggcA
 8183>cacctagatccttttaaattaaaaatgaagttttaaatcaatctaaagtatatatgagtaaacttggtctgacagttaccaatgcttaatcagtgaggca>8282
    1<~~~~~~~~~~~~~~~~~~~~~~~~~~~~~~~~~~~~~~~~~~~~~~~~~~~~~~~~~~~~~~~~~~~~~~~~~~~~~~~~~~~~~~~~~~~~~~~~~~~~<1
    1<~~~~~~~~~~~~~~~~~~~~~~~~~~~~~~~~~~~~~~~~~~~~~~~~~~~~~~~~~~~~~~~~~~~~~~~~~~~~~~~~~~~~~~~~~~~~~~~~~~~~<1
    1<~~~~~~~~~~~~~~~~~~~~~~~~~~~~~~~~~~~~~~~~~~~~~~~~~~~~~~~~~~~~~~~~~~~~~~~~~~~~~~~~~~~~~~~~~~~~~~~~~~~~<1
    7<----------------------------------------------------------------------------------------------------<7
```

```
   1<~~~~~~~~~~~~~~~~~~~~~~~~~~~~~~~~~~~~~~~~~~~~~~~~~~~~~~~~~~~~~~~~~~~~~~~~~~~~~~~~~~~~~~~~~~~~~~~~<1
   7<----------------------------------------------------------------------------------------------------<7

            *         *         *         *         *         *         *         *         *         *
     G  I  E  A  I  Q  R  N  R  E  D  M  T  A  Q  S  G  T  T  Y  I  V  V  I  R  S  P  K  G  D  P  G  L  A
     CCtatCTCagcGATctgTCTattTCGttcATCcatAGTtgcCTGactGCCcgtCGTgtaGATaacTACgatACGggaGGGcttACCatcTGGcccCAGtg
8283>cctatctcagcgatctgtctatttcgttcatccatagttgcctgactgcccgtcgtgtagataactacgatacgggagggcttaccatctggccccagtg>8382
   1<~~~~~~~~~~~~~~~~~~~~~~~~~~~~~~~~~~~~~~~~~~~~~~~~~~~~~~~~~~~~~~~~~~~~~~~~~~~~~~~~~~~~~~~~~~~~~~~~<1
   1<~~~~~~~~~~~~~~~~~~~~~~~~~~~~~~~~~~~~~~~~~~~~~~~~~~~~~~~~~~~~~~~~~~~~~~~~~~~~~~~~~~~~~~~~~~~~~~~~<1
   1<~~~~~~~~~~~~~~~~~~~~~~~~~~~~~~~~~~~~~~~~~~~~~~~~~~~~~~~~~~~~~~~~~~~~~~~~~~~~~~~~~~~~~~~~~~~~~~~~<1
   7<----------------------------------------------------------------------------------------------------<7
   1<~~~~~~~~~~~~~~~~~~~~~~~~~~~~~~~~~~~~~~~~~~~~~~~~~~~~~~~~~~~~~~~~~~~~~~~~~~~~~~~~~~~~~~~~~~~~~~~~<1
   7<----------------------------------------------------------------------------------------------------<7

            *         *         *         *         *         *         *         *         *         *
       A  I  I  G  R  S  G  R  E  G  A  G  S  K  D  A  I  F  W  G  A  P  L  A  S  R  L  L  P  G  A  V  K
     cTGCaatGATaccGCGagaCCCacgCTCaccGGCtccAGAtttATCagcAATaaaCCAgccAGCcggAAGggcCGAgcgCAGaagTGGtccTGCaacTTT
8383>ctgcaatgataccgcgagacccacgctcaccggctccagatttatcagcaataaaccagccagccggaagggccgagcgcagaagtggtcctgcaacttt>8482
   1<~~~~~~~~~~~~~~~~~~~~~~~~~~~~~~~~~~~~~~~~~~~~~~~~~~~~~~~~~~~~~~~~~~~~~~~~~~~~~~~~~~~~~~~~~~~~~~~~<1
   1<~~~~~~~~~~~~~~~~~~~~~~~~~~~~~~~~~~~~~~~~~~~~~~~~~~~~~~~~~~~~~~~~~~~~~~~~~~~~~~~~~~~~~~~~~~~~~~~~<1
   1<~~~~~~~~~~~~~~~~~~~~~~~~~~~~~~~~~~~~~~~~~~~~~~~~~~~~~~~~~~~~~~~~~~~~~~~~~~~~~~~~~~~~~~~~~~~~~~~~<1
   7<----------------------------------------------------------------------------------------------------<7
   1<~~~~~~~~~~~~~~~~~~~~~~~~~~~~~~~~~~~~~~~~~~~~~~~~~~~~~~~~~~~~~~~~~~~~~~~~~~~~~~~~~~~~~~~~~~~~~~~~<1
   7<----------------------------------------------------------------------------------------------------<7

            *         *         *         *         *         *         *         *         *         *
      D  A  E  M  W  D  I  L  Q  Q  R  S  A  L  T  L  L  E  G  T  L  L  K  R  L  T  T  A  M  A  V  P  M
     atcCGCctcCATccaGTCtatTAAttgTTGccgGGAagcTAGagtAAGtagTTCgccAGTtaaTAGtttGCGcaaCGTtgtTGCcatTGCtacAGGcatC
8483>atccgcctccatccagtctattaattgttgccgggaagctagagtaagtagttcgccagttaatagtttgcgcaacgttgttgccattgctacaggcatc>8582
   1<~~~~~~~~~~~~~~~~~~~~~~~~~~~~~~~~~~~~~~~~~~~~~~~~~~~~~~~~~~~~~~~~~~~~~~~~~~~~~~~~~~~~~~~~~~~~~~~~<1
   1<~~~~~~~~~~~~~~~~~~~~~~~~~~~~~~~~~~~~~~~~~~~~~~~~~~~~~~~~~~~~~~~~~~~~~~~~~~~~~~~~~~~~~~~~~~~~~~~~<1
   1<~~~~~~~~~~~~~~~~~~~~~~~~~~~~~~~~~~~~~~~~~~~~~~~~~~~~~~~~~~~~~~~~~~~~~~~~~~~~~~~~~~~~~~~~~~~~~~~~<1
   7<----------------------------------------------------------------------------------------------------<7
   1<~~~~~~~~~~~~~~~~~~~~~~~~~~~~~~~~~~~~~~~~~~~~~~~~~~~~~~~~~~~~~~~~~~~~~~~~~~~~~~~~~~~~~~~~~~~~~~~~<1
   7<----------------------------------------------------------------------------------------------------<7

            *         *         *         *         *         *         *         *         *         *
     T  T  D  R  E  D  N  P  I  A  E  N  L  E  P  E  W  R  D  L  R  T  V  H  D  G  M  N  H  F  F  A  T  L
     GTggtGTCacgCTCgtcGTTtggTATggcTTCattCAGctcCGGttcCCAacgATCaagGCGagtTACatgATCcccCATgttGTGaaaAAAagcGGTta
```

```
8583>gtggtgtcacgctcgtcgtttggtatggcttcattcagctccggttcccaacgatcaaggcgagttacatgatcccccatgttgtgaaaaaaagcggtta>8682
   1<~~~~~~~~~~~~~~~~~~~~~~~~~~~~~~~~~~~~~~~~~~~~~~~~~~~~~~~~~~~~~~~~~~~~~~~~~~~~~~~~~~~~~~~~~~~~~~~~~~~<1
   1<~~~~~~~~~~~~~~~~~~~~~~~~~~~~~~~~~~~~~~~~~~~~~~~~~~~~~~~~~~~~~~~~~~~~~~~~~~~~~~~~~~~~~~~~~~~~~~~~~~~<1
   1<~~~~~~~~~~~~~~~~~~~~~~~~~~~~~~~~~~~~~~~~~~~~~~~~~~~~~~~~~~~~~~~~~~~~~~~~~~~~~~~~~~~~~~~~~~~~~~~~~~~<1
   7<----------------------------------------------------------------------------------------------------<7
   1<~~~~~~~~~~~~~~~~~~~~~~~~~~~~~~~~~~~~~~~~~~~~~~~~~~~~~~~~~~~~~~~~~~~~~~~~~~~~~~~~~~~~~~~~~~~~~~~~~~~<1
   7<----------------------------------------------------------------------------------------------------<7

              *         *         *         *         *         *         *         *         *         *
       E  K  P  G  G  I  T  T  L  L  L  N  A  A  T  N  D  S  M  T  I  A  A  S  C  L  E  R  V  T  M  G  D
     gCTCcttCGGtccTCCgatCGTtgtCAGaagTAAgttGGCcgcAGTgttATCactCATggtTATggcAGCactGCAtaaTTCtctTACtgtCATgccATC
8683>gctccttcggtcctccgatcgttgtcagaagtaagttggccgcagtgttatcactcatggttatggcagcactgcataattctcttactgtcatgccatc>8782
   1<~~~~~~~~~~~~~~~~~~~~~~~~~~~~~~~~~~~~~~~~~~~~~~~~~~~~~~~~~~~~~~~~~~~~~~~~~~~~~~~~~~~~~~~~~~~~~~~~~~~<1
   1<~~~~~~~~~~~~~~~~~~~~~~~~~~~~~~~~~~~~~~~~~~~~~~~~~~~~~~~~~~~~~~~~~~~~~~~~~~~~~~~~~~~~~~~~~~~~~~~~~~~<1
   1<~~~~~~~~~~~~~~~~~~~~~~~~~~~~~~~~~~~~~~~~~~~~~~~~~~~~~~~~~~~~~~~~~~~~~~~~~~~~~~~~~~~~~~~~~~~~~~~~~~~<1
   7<----------------------------------------------------------------------------------------------------<7
   1<~~~~~~~~~~~~~~~~~~~~~~~~~~~~~~~~~~~~~~~~~~~~~~~~~~~~~~~~~~~~~~~~~~~~~~~~~~~~~~~~~~~~~~~~~~~~~~~~~~~<1
   7<----------------------------------------------------------------------------------------------------<7

              *         *         *         *         *         *         *         *         *         *
      T  L  H  K  E  T  V  P  S  Y  E  V  L  D  N  Q  S  Y  H  I  R  R  G  L  Q  E  Q  G  A  D  I  R  S
     cgtAAGatgCTTttcTGTgacTGGtgaGTActcAACcaaGTCattCTGagaATAgtgTATgcgGCGaccGAGttgCTCttgCCCggcGTCaatACGggaT
8783>cgtaagatgcttttctgtgactggtgagtactcaaccaagtcattctgagaatagtgtatgcggcgaccgagttgctcttgcccggcgtcaatacgggat>8882
   1<~~~~~~~~~~~~~~~~~~~~~~~~~~~~~~~~~~~~~~~~~~~~~~~~~~~~~~~~~~~~~~~~~~~~~~~~~~~~~~~~~~~~~~~~~~~~~~~~~~~<1
   1<~~~~~~~~~~~~~~~~~~~~~~~~~~~~~~~~~~~~~~~~~~~~~~~~~~~~~~~~~~~~~~~~~~~~~~~~~~~~~~~~~~~~~~~~~~~~~~~~~~~<1
   1<~~~~~~~~~~~~~~~~~~~~~~~~~~~~~~~~~~~~~~~~~~~~~~~~~~~~~~~~~~~~~~~~~~~~~~~~~~~~~~~~~~~~~~~~~~~~~~~~~~~<1
   7<----------------------------------------------------------------------------------------------------<7
   1<~~~~~~~~~~~~~~~~~~~~~~~~~~~~~~~~~~~~~~~~~~~~~~~~~~~~~~~~~~~~~~~~~~~~~~~~~~~~~~~~~~~~~~~~~~~~~~~~~~~<1
   7<----------------------------------------------------------------------------------------------------<7

              *         *         *         *         *         *         *         *         *         *
     L  V  A  G  C  L  L  V  K  F  T  S  M  M  P  F  R  E  E  P  R  F  S  E  L  I  K  G  S  N  L  D  L  E
     AAtacCGCgccACAtagCAGaacTTTaaaAGTgctCATcatTGGaaaACGttcTTCgggGCGaaaACTctcAAGgatCTTaccGCTgttGAGatcCAGtt
8883>aataccgcgccacatagcagaactttaaaagtgctcatcattggaaaacgttcttcggggcgaaaactctcaaggatcttaccgctgttgagatccagtt>8982
   1<~~~~~~~~~~~~~~~~~~~~~~~~~~~~~~~~~~~~~~~~~~~~~~~~~~~~~~~~~~~~~~~~~~~~~~~~~~~~~~~~~~~~~~~~~~~~~~~~~~~<1
   1<~~~~~~~~~~~~~~~~~~~~~~~~~~~~~~~~~~~~~~~~~~~~~~~~~~~~~~~~~~~~~~~~~~~~~~~~~~~~~~~~~~~~~~~~~~~~~~~~~~~<1
   1<~~~~~~~~~~~~~~~~~~~~~~~~~~~~~~~~~~~~~~~~~~~~~~~~~~~~~~~~~~~~~~~~~~~~~~~~~~~~~~~~~~~~~~~~~~~~~~~~~~~<1
   7<----------------------------------------------------------------------------------------------------<7
   1<~~~~~~~~~~~~~~~~~~~~~~~~~~~~~~~~~~~~~~~~~~~~~~~~~~~~~~~~~~~~~~~~~~~~~~~~~~~~~~~~~~~~~~~~~~~~~~~~~~~<1
   7<----------------------------------------------------------------------------------------------------<7
```

```
              *         *         *         *         *         *         *         *         *         *
        I  Y  G  V  R  A  G  L  Q  D  E  A  D  K  V  K  V  L  T  E  P  H  A  F  V  P  L  C  F  A  A  F  F
     cGATgtaACCcacTCGtgcACCcaaCTGatcTTCagcATCtttTACtttCACcagCGTttcTGGgtgAGCaaaAACaggAAGgcaAAAtgcCGCaaaAAA
8983>cgatgtaacccactcgtgcacccaactgatcttcagcatcttttactttcaccagcgtttctgggtgagcaaaaacaggaaggcaaaatgccgcaaaaaa>9082
   1<~~~~~~~~~~~~~~~~~~~~~~~~~~~~~~~~~~~~~~~~~~~~~~~~~~~~~~~~~~~~~~~~~~~~~~~~~~~~~~~~~~~~~~~~~~~~~~~~~<1
   1<~~~~~~~~~~~~~~~~~~~~~~~~~~~~~~~~~~~~~~~~~~~~~~~~~~~~~~~~~~~~~~~~~~~~~~~~~~~~~~~~~~~~~~~~~~~~~~~~<1
   1<~~~~~~~~~~~~~~~~~~~~~~~~~~~~~~~~~~~~~~~~~~~~~~~~~~~~~~~~~~~~~~~~~~~~~~~~~~~~~~~~~~~~~~~~~~~~~~~~<1
   7<----------------------------------------------------------------------------------------------------<7
   1<~~~~~~~~~~~~~~~~~~~~~~~~~~~~~~~~~~~~~~~~~~~~~~~~~~~~~~~~~~~~~~~~~~~~~~~~~~~~~~~~~~~~~~~~~~~~~~~~<1
   7<----------------------------------------------------------------------------------------------------<7

              *         *         *         *         *         *         *         *         *         *
      P  I  L  A  V  R  F  H  Q  I  S  M
     gggAATaagGGCgacACGgaaATGttgAATactCAT
9083>gggaataagggcgacacggaaatgttgaatactcatactcttcctttttcaatattattgaagcatttatcagggttattgtctcatgagcggatacata>9182
   1<~~~~~~~~~~~~~~~~~~~~~~~~~~~~~~~~~~~~~~~~~~~~~~~~~~~~~~~~~~~~~~~~~~~~~~~~~~~~~~~~~~~~~~~~~~~~~~~~<1
   1<~~~~~~~~~~~~~~~~~~~~~~~~~~~~~~~~~~~~~~~~~~~~~~~~~~~~~~~~~~~~~~~~~~~~~~~~~~~~~~~~~~~~~~~~~~~~~~~~<1
   1<~~~~~~~~~~~~~~~~~~~~~~~~~~~~~~~~~~~~~~~~~~~~~~~~~~~~~~~~~~~~~~~~~~~~~~~~~~~~~~~~~~~~~~~~~~~~~~~~<1
   6<-------------------------------------------------------------------------------------------GGAT-----<3
   1<~~~~~~~~~~~~~~~~~~~~~~~~~~~~~~~~~~~~~~~~~~~~~~~~~~~~~~~~~~~~~~~~~~~~~~~~~~~~~~~~~~~~~~~~~~~~~~~~<1
   7<----------------------------------------------------------------------------------------------------<7

              *         *         *         *         *         *         *         *         *         *
9183>tttgaatgtatttagaaaaataaacaaataggggttccgcgcacatttccccgaaaagtgccacctgacgtcttattatcatgacattaacctataaaaa>9282
   1<~~~~~~~~~~~~~~~~~~~~~~~~~~~~~~~~~~~~~~~~~~~~~~~~~~~~~~~~~~~~~~~~~~~~~~~~~~~~~~~~~~~~~~~~~~~~~~~~<1
   1<~~~~~~~~~~~~~~~~~~~~~~~~~~~~~~~~~~~~~~~~~~~~~~~~~~~~~~~~~~~~~~~~~~~~~~~~~~~~~~~~~~~~~~~~~~~~~~~~<1
   1<~~~~~~~~~~~~~~~~~~~~~~~~~~~~~~~~~~~~~~~~~~~~~~~~~~~~~~~~~~~~~~~~~~~~~~~~~~~~~~~~~~~~~~~~~~~~~~~~<1
   3<----------------------------------------------------------------------------------------------------<3
   1<~~~~~~~~~~~~~~~~~~~~~~~~~~~~~~~~~~~~~~~~~~~~~~~~~~~~~~~~~~~~~~~~~~~~~~~~~~~~~~~~~~~~~~~~~~~~~~~~<1
   6<---------------------------------------------------------GTG----------------------------------------<4

              *         *
9283>taggcgtatcacgaggccctttcgtc>9308
   1<~~~~~~~~~~~~~~~~~~~~~~~~~~<1
   1<~~~~~~~~~~~~~~~~~~~~~~~~~~<1
   1<~~~~~~~~~~~~~~~~~~~~~~~~~~<1
   2<------------------------TC<1
   1<~~~~~~~~~~~~~~~~~~~~~~~~~~<1
```

3<--------------------TTT~~~~<1
