## Supplementary material for "AssemblyTron: Flexible automation of DNA assembly with Opentrons OT-2 lab robots": File 10

Mon Sep 12, 2022 11:11 EDT
-ARF7-pdar-pmas00002.gb from 1 to 10043to_F10_IVA_rxn2_C6+206_TCYC1_Seq_R.ab1--  Matches:1130; Mismatches:0;
Gaps:3632; Unattempted:5427_E10_IVA_rxn2_C5+206_TCYC1_Seq_R.ab1--  Matches:1300; Mismatches:2; Gaps:3487;
Unattempted:5256_D10_IVA_rxn2_C4+206_TCYC1_Seq_R.ab1--  Matches:996; Mismatches:1; Gaps:3777;
Unattempted:5562_H03_IVA_rxn2_C3+206_TCYC1_Seq_R.ab1--  Matches:579; Mismatches:2; Gaps:3624;
Unattempted:5983_F03_IVA_rxn2_C1+206_TCYC1_Seq_R.ab1--  Matches:1190; Mismatches:0; Gaps:3486;
Unattempted:5367_G03_IVA_rxn2_C2+206_TCYC1_Seq_R.ab1--  Matches:579; Mismatches:2; Gaps:3624; Unattempted:5983

```
                  *         *         *         *         *         *         *         *         *         *
      1>tcgcgcgtttcggtgatgacggtgaaaacctctgacacatgcagctcccggagacggtcacagcttgtctgtaagcggatgccgggagcagacaagcccg>100
   1434<T[redacted]<1434
   1450<T[redacted]<1450
   1479<~~~~~~~~~~~~~~~~~~~~~~~~~~~~~~~~~~~~~~~~~~~~~~~~~~~~~~~~~~~~~~~~~~~~~~~~~~~~~~~~~~~~~~~~~~~~~~~~~~<1479
   1395<~~~~~~~~~~~~~~~~~~~~~~~~~~~~~~~~~~~~~~~~~~~~~~~~~~~~~~~~~~~~~~~~~~~~~~~~~~~~~~~~~~~~~~~~~~~~~~~~~~<1395
   1388<~~~~~~~~~~~~~~~~~~~~~~~~~~~~~~~~~~~~~~~~~~~~~~~~~~~~~~~~~~~~~~~~~~~~~~~~~~~~~~~~~~~~~~~~~~~~~~~~~~<1388
   1395<~~~~~~~~~~~~~~~~~~~~~~~~~~~~~~~~~~~~~~~~~~~~~~~~~~~~~~~~~~~~~~~~~~~~~~~~~~~~~~~~~~~~~~~~~~~~~~~~~~<1395

                  *         *         *         *         *         *         *         *         *         *
    101>tcagggcgcgtcagcgggtgttggcgggtgtcggggctggcttaactatgcggcatcagagcagattgtactgagagtgcaccataattcgtttaaaccg>200
   1434<----------------------------------------------------------------------------------------------------<1434
   1450<----------------------------------------------------------------------------------------------------<1450
   1479<~~~~~~~~~~~~~~~~~~~~~~~~~~~~~~~~~~~~~~~~~~~~~~~~~~~~~~~~~~~~~~~~~~~~~~~~~~~~~~~~~~~~~~~~~~~~~~~~~~<1479
   1395<~~~~~~~~~~~~~~~~~~~~~~~~~~~~~~~~~~~~~~~~~~~~~~~~~~~~~~~~~~~~~~~~~~~~~~~~~~~~~~~~~~~~~~~~~~~~~~~~~~<1395
   1388<~~~~~~~~~~~~~~~~~~~~~~~~~~~~~~~~~~~~~~~~~~~~~~~~~~~~~~~~~~~~~~~~~~~~~~~~~~~~~~~~~~~~~~~~~~~~~~~~~~<1388
   1395<~~~~~~~~~~~~~~~~~~~~~~~~~~~~~~~~~~~~~~~~~~~~~~~~~~~~~~~~~~~~~~~~~~~~~~~~~~~~~~~~~~~~~~~~~~~~~~~~~~<1395

                  *         *         *         *         *         *         *         *         *         *
    201>ttttaagagcttggtgagcgctaggagtcactgccaggtatcgtttgaacacggcattagtcagggaagtcataacacagtcctttcccgcaattttctt>300
   1434<----------------------------------------------------------------------------------------------------<1434
   1450<----------------------------------------------------------------------------------------------------<1450
   1479<~~~~~~~~~~~~~~~~~~~~~~~~~~~~~~~~~~~~~~~~~~~~~~~~~~~~~~~~~~~~~~~~~~~~~~~~~~~~~~~~~~~~~~~~~~~~~~~~~~<1479
   1395<~~~~~~~~~~~~~~~~~~~~~~~~~~~~~~~~~~~~~~~~~~~~~~~~~~~~~~~~~~~~~~~~~~~~~~~~~~~~~~~~~~~~~~~~~~~~~~~~~~<1395
   1388<~~~~~~~~~~~~~~~~~~~~~~~~~~~~~~~~~~~~~~~~~~~~~~~~~~~~~~~~~~~~~~~~~~~~~~~~~~~~~~~~~~~~~~~~~~~~~~~~~~<1388
   1395<~~~~~~~~~~~~~~~~~~~~~~~~~~~~~~~~~~~~~~~~~~~~~~~~~~~~~~~~~~~~~~~~~~~~~~~~~~~~~~~~~~~~~~~~~~~~~~~~~~<1395

                  *         *         *         *         *         *         *         *         *         *
    301>tttctattactcttggcctcctctagtacactctatatttttttatgcctcggtaatgattttcattttttttttttccacctagcggatgactctttttt>400
   1434<----------------------------------------------------------------------------------------------------<1434
   1450<----------------------------------------------------------------------------------------------------<1450
   1479<~~~~~~~~~~~~~~~~~~~~~~~~~~~~~~~~~~~~~~~~~~~~~~~~~~~~~~~~~~~~~~~~~~~~~~~~~~~~~~~~~~~~~~~~~~~~~~~~~~<1479
```

```
1395<~~~~~~~~~~~~~~~~~~~~~~~~~~~~~~~~~~~~~~~~~~~~~~~~~~~~~~~~~~~~~~~~~~~~~~~~~~~~~~~~~~~~~~~~~~~~~~~~<1395
1388<~~~~~~~~~~~~~~~~~~~~~~~~~~~~~~~~~~~~~~~~~~~~~~~~~~~~~~~~~~~~~~~~~~~~~~~~~~~~~~~~~~~~~~~~~~~~~~~~<1388
1395<~~~~~~~~~~~~~~~~~~~~~~~~~~~~~~~~~~~~~~~~~~~~~~~~~~~~~~~~~~~~~~~~~~~~~~~~~~~~~~~~~~~~~~~~~~~~~~~~<1395

              *         *         *         *         *         *         *         *         *         *
 401>tttcttagcgattggcattatcacataatgaattatacattatataaagtaatgtgatttcttcgaagaatatactaaaaaatgagcaggcaagataaac>500
1434<----------------------------------------------------------------------------------------------------<1434
1450<----------------------------------------------------------------------------------------------------<1450
1479<~~~~~~~~~~~~~~~~~~~~~~~~~~~~~~~~~~~~~~~~~~~~~~~~~~~~~~~~~~~~~~~~~~~~~~~~~~~~~~~~~~~~~~~~~~~~~~~~<1479
1395<~~~~~~~~~~~~~~~~~~~~~~~~~~~~~~~~~~~~~~~~~~~~~~~~~~~~~~~~~~~~~~~~~~~~~~~~~~~~~~~~~~~~~~~~~~~~~~~~<1395
1388<~~~~~~~~~~~~~~~~~~~~~~~~~~~~~~~~~~~~~~~~~~~~~~~~~~~~~~~~~~~~~~~~~~~~~~~~~~~~~~~~~~~~~~~~~~~~~~~~<1388
1395<~~~~~~~~~~~~~~~~~~~~~~~~~~~~~~~~~~~~~~~~~~~~~~~~~~~~~~~~~~~~~~~~~~~~~~~~~~~~~~~~~~~~~~~~~~~~~~~~<1395

              *         *         *         *         *         *         *         *         *         *
 501>gaaggcaaagatgacagagcagaaagccctagtaaagcgtattacaaatgaaaccaagattcagattgcgatctctttaaagggtggtcccctagcgata>600
1434<----------------------------------------------------------------------------------------------------<1434
1450<----------------------------------------------------------------------------------------------------<1450
1479<~~~~~~~~~~~~~~~~~~~~~~~~~~~~~~~~~~~~~~~~~~~~~~~~~~~~~~~~~~~~~~~~~~~~~~~~~~~~~~~~~~~~~~~~~~~~~~~<1479
1395<~~~~~~~~~~~~~~~~~~~~~~~~~~~~~~~~~~~~~~~~~~~~~~~~~~~~~~~~~~~~~~~~~~~~~~~~~~~~~~~~~~~~~~~~~~~~~~~~<1395
1388<~~~~~~~~~~~~~~~~~~~~~~~~~~~~~~~~~~~~~~~~~~~~~~~~~~~~~~~~~~~~~~~~~~~~~~~~~~~~~~~~~~~~~~~~~~~~~~~~<1388
1395<~~~~~~~~~~~~~~~~~~~~~~~~~~~~~~~~~~~~~~~~~~~~~~~~~~~~~~~~~~~~~~~~~~~~~~~~~~~~~~~~~~~~~~~~~~~~~~~~<1395

              *         *         *         *         *         *         *         *         *         *
 601>gagcactcgatcttcccagaaaaagaggcagaagcagtagcagaacaggccacacaatcgcaagtgattaacgtccacacaggtatagggtttctggacc>700
1434<----------------------------------------------------------------------------------------------------<1434
1450<----------------------------------------------------------------------------------------------------<1450
1479<~~~~~~~~~~~~~~~~~~~~~~~~~~~~~~~~~~~~~~~~~~~~~~~~~~~~~~~~~~~~~~~~~~~~~~~~~~~~~~~~~~~~~~~~~~~~~~~~<1479
1395<~~~~~~~~~~~~~~~~~~~~~~~~~~~~~~~~~~~~~~~~~~~~~~~~~~~~~~~~~~~~~~~~~~~~~~~~~~~~~~~~~~~~~~~~~~~~~~~~<1395
1388<~~~~~~~~~~~~~~~~~~~~~~~~~~~~~~~~~~~~~~~~~~~~~~~~~~~~~~~~~~~~~~~~~~~~~~~~~~~~~~~~~~~~~~~~~~~~~~~~<1388
1395<~~~~~~~~~~~~~~~~~~~~~~~~~~~~~~~~~~~~~~~~~~~~~~~~~~~~~~~~~~~~~~~~~~~~~~~~~~~~~~~~~~~~~~~~~~~~~~~~<1395

              *         *         *         *         *         *         *         *         *         *
 701>atatgatacatgctctggccaagcattccggctggtcgctaatcgttgagtgcattggtgacttacacatagacgaccatcacaccactgaagactgcgg>800
1434<----------------------------------------------------------------------------------------------------<1434
1450<----------------------------------------------------------------------------------------------------<1450
1479<~~~~~~~~~~~~~~~~~~~~~~~~~~~~~~~~~~~~~~~~~~~~~~~~~~~~~~~~~~~~~~~~~~~~~~~~~~~~~~~~~~~~~~~~~~~~~~~~<1479
1395<~~~~~~~~~~~~~~~~~~~~~~~~~~~~~~~~~~~~~~~~~~~~~~~~~~~~~~~~~~~~~~~~~~~~~~~~~~~~~~~~~~~~~~~~~~~~~~~~<1395
1388<~~~~~~~~~~~~~~~~~~~~~~~~~~~~~~~~~~~~~~~~~~~~~~~~~~~~~~~~~~~~~~~~~~~~~~~~~~~~~~~~~~~~~~~~~~~~~~~~<1388
1395<~~~~~~~~~~~~~~~~~~~~~~~~~~~~~~~~~~~~~~~~~~~~~~~~~~~~~~~~~~~~~~~~~~~~~~~~~~~~~~~~~~~~~~~~~~~~~~~~<1395
```

```
              *         *         *         *         *         *         *         *         *         *
 801>gattgctctcggtcaagcttttaaagaggccctaggggccgtgcgtggagtaaaaaggtttggatcaggatttgcgcctttggatgaggcactttccaga>900
1434<----------------------------------------------------------------------------------------------------<1434
1450<----------------------------------------------------------------------------------------------------<1450
1479<~~~~~~~~~~~~~~~~~~~~~~~~~~~~~~~~~~~~~~~~~~~~~~~~~~~~~~~~~~~~~~~~~~~~~~~~~~~~~~~~~~~~~~~~~~~~~~~~~~~~<1479
1395<~~~~~~~~~~~~~~~~~~~~~~~~~~~~~~~~~~~~~~~~~~~~~~~~~~~~~~~~~~~~~~~~~~~~~~~~~~~~~~~~~~~~~~~~~~~~~~~~~~~~<1395
1388<~~~~~~~~~~~~~~~~~~~~~~~~~~~~~~~~~~~~~~~~~~~~~~~~~~~~~~~~~~~~~~~~~~~~~~~~~~~~~~~~~~~~~~~~~~~~~~~~~~~~<1388
1395<~~~~~~~~~~~~~~~~~~~~~~~~~~~~~~~~~~~~~~~~~~~~~~~~~~~~~~~~~~~~~~~~~~~~~~~~~~~~~~~~~~~~~~~~~~~~~~~~~~~~<1395

              *         *         *         *         *         *         *         *         *         *
 901>gcggtggtagatctttcgaacaggccgtacgcagttgtcgaacttggtttgcaaagggagaaagtaggagatctctcttgcgagatgatcccgcattttc>1000
1434<----------------------------------------------------------------------------------------------------<1434
1450<----------------------------------------------------------------------------------------------------<1450
1479<~~~~~~~~~~~~~~~~~~~~~~~~~~~~~~~~~~~~~~~~~~~~~~~~~~~~~~~~~~~~~~~~~~~~~~~~~~~~~~~~~~~~~~~~~~~~~~~~~~~~<1479
1395<~~~~~~~~~~~~~~~~~~~~~~~~~~~~~~~~~~~~~~~~~~~~~~~~~~~~~~~~~~~~~~~~~~~~~~~~~~~~~~~~~~~~~~~~~~~~~~~~~~~~<1395
1388<~~~~~~~~~~~~~~~~~~~~~~~~~~~~~~~~~~~~~~~~~~~~~~~~~~~~~~~~~~~~~~~~~~~~~~~~~~~~~~~~~~~~~~~~~~~~~~~~~~~~<1388
1395<~~~~~~~~~~~~~~~~~~~~~~~~~~~~~~~~~~~~~~~~~~~~~~~~~~~~~~~~~~~~~~~~~~~~~~~~~~~~~~~~~~~~~~~~~~~~~~~~~~~~<1395

              *         *         *         *         *         *         *         *         *         *
1001>ttgaaagctttgcagaggctagcagaattaccctccacgttgattgtctgcgaggcaagaatgatcatcaccgtagtgagagtgcgttcaaggctcttgc>1100
1434<----------------------------------------------------------------------------------------------------<1434
1450<----------------------------------------------------------------------------------------------------<1450
1479<~~~~~~~~~~~~~~~~~~~~~~~~~~~~~~~~~~~~~~~~~~~~~~~~~~~~~~~~~~~~~~~~~~~~~~~~~~~~~~~~~~~~~~~~~~~~~~~~~~~~<1479
1395<~~~~~~~~~~~~~~~~~~~~~~~~~~~~~~~~~~~~~~~~~~~~~~~~~~~~~~~~~~~~~~~~~~~~~~~~~~~~~~~~~~~~~~~~~~~~~~~~~~~~<1395
1388<~~~~~~~~~~~~~~~~~~~~~~~~~~~~~~~~~~~~~~~~~~~~~~~~~~~~~~~~~~~~~~~~~~~~~~~~~~~~~~~~~~~~~~~~~~~~~~~~~~~~<1388
1395<~~~~~~~~~~~~~~~~~~~~~~~~~~~~~~~~~~~~~~~~~~~~~~~~~~~~~~~~~~~~~~~~~~~~~~~~~~~~~~~~~~~~~~~~~~~~~~~~~~~~<1395

              *         *         *         *         *         *         *         *         *         *
1101>ggttgccataagagaagccacctcgcccaatggtaccaacgatgttccctccaccaaaggtgttcttatgtaggcgaatttcttatgatttatgattttt>1200
1434<----------------------------------------------------------------------------------------------------<1434
1450<----------------------------------------------------------------------------------------------------<1450
1479<~~~~~~~~~~~~~~~~~~~~~~~~~~~~~~~~~~~~~~~~~~~~~~~~~~~~~~~~~~~~~~~~~~~~~~~~~~~~~~~~~~~~~~~~~~~~~~~~~~~~<1479
1395<~~~~~~~~~~~~~~~~~~~~~~~~~~~~~~~~~~~~~~~~~~~~~~~~~~~~~~~~~~~~~~~~~~~~~~~~~~~~~~~~~~~~~~~~~~~~~~~~~~~~<1395
1388<~~~~~~~~~~~~~~~~~~~~~~~~~~~~~~~~~~~~~~~~~~~~~~~~~~~~~~~~~~~~~~~~~~~~~~~~~~~~~~~~~~~~~~~~~~~~~~~~~~~~<1388
1395<~~~~~~~~~~~~~~~~~~~~~~~~~~~~~~~~~~~~~~~~~~~~~~~~~~~~~~~~~~~~~~~~~~~~~~~~~~~~~~~~~~~~~~~~~~~~~~~~~~~~<1395

              *         *         *         *         *         *         *         *         *         *
```

```
1201>attattaaataagttataaaaaaaataagtgtatacaaattttaaagtgactcttaggttttaaaacgaaaattcttattcttgagtaactctttcctgt>1300
1434<----------------------------------------------------------------------------------------------------<1434
1450<----------------------------------------------------------------------------------------------------<1450
1479<~~~~~~~~~~~~~~~~~~~~~~~~~~~~~~~~~~~~~~~~~~~~~~~~~~~~~~~~~~~~~~~~~~~~~~~~~~~~~~~~~~~~~~~~~~~~~~~~~~~~<1479
1395<~~~~~~~~~~~~~~~~~~~~~~~~~~~~~~~~~~~~~~~~~~~~~~~~~~~~~~~~~~~~~~~~~~~~~~~~~~~~~~~~~~~~~~~~~~~~~~~~~~~~<1395
1388<~~~~~~~~~~~~~~~~~~~~~~~~~~~~~~~~~~~~~~~~~~~~~~~~~~~~~~~~~~~~~~~~~~~~~~~~~~~~~~~~~~~~~~~~~~~~~~~~~~~~<1388
1395<~~~~~~~~~~~~~~~~~~~~~~~~~~~~~~~~~~~~~~~~~~~~~~~~~~~~~~~~~~~~~~~~~~~~~~~~~~~~~~~~~~~~~~~~~~~~~~~~~~~~<1395

              *         *         *         *         *         *         *         *         *         *
1301>aggtcaggttgctttctcaggtatagcatgaggtcgctcttattgaccacacctcaagaaatgatggtaaatgaaataggaaatcaaggagcatgaaggc>1400
1434<----------------------------------------------------------------------------------------------------<1434
1450<----------------------------------------------------------------------------------------------------<1450
1479<~~~~~~~~~~~~~~~~~~~~~~~~~~~~~~~~~~~~~~~~~~~~~~~~~~~~~~~~~~~~~~~~~~~~~~~~~~~~~~~~~~~~~~~~~~~~~~~~~~~~<1479
1395<~~~~~~~~~~~~~~~~~~~~~~~~~~~~~~~~~~~~~~~~~~~~~~~~~~~~~~~~~~~~~~~~~~~~~~~~~~~~~~~~~~~~~~~~~~~~~~~~~~~~<1395
1388<~~~~~~~~~~~~~~~~~~~~~~~~~~~~~~~~~~~~~~~~~~~~~~~~~~~~~~~~~~~~~~~~~~~~~~~~~~~~~~~~~~~~~~~~~~~~~~~~~~~~<1388
1395<~~~~~~~~~~~~~~~~~~~~~~~~~~~~~~~~~~~~~~~~~~~~~~~~~~~~~~~~~~~~~~~~~~~~~~~~~~~~~~~~~~~~~~~~~~~~~~~~~~~~<1395

              *         *         *         *         *         *         *         *         *         *
1401>aaaagacaaatataagggtcgaacgaaaaataaagtgaaaagtgttgatatgatgtatttggctttgcggcgccgaaaaaacgagtttacgcaattgcac>1500
1434<----------------------------------------------------------------------------------------------------<1434
1450<----------------------------------------------------------------------------------------------------<1450
1479<~~~~~~~~~~~~~~~~~~~~~~~~~~~~~~~~~~~~~~~~~~~~~~~~~~~~~~~~~~~~~~~~~~~~~~~~~~~~~~~~~~~~~~~~~~~~~~~~~~~~<1479
1395<~~~~~~~~~~~~~~~~~~~~~~~~~~~~~~~~~~~~~~~~~~~~~~~~~~~~~~~~~~~~~~~~~~~~~~~~~~~~~~~~~~~~~~~~~~~~~~~~~~~~<1395
1388<~~~~~~~~~~~~~~~~~~~~~~~~~~~~~~~~~~~~~~~~~~~~~~~~~~~~~~~~~~~~~~~~~~~~~~~~~~~~~~~~~~~~~~~~~~~~~~~~~~~~<1388
1395<~~~~~~~~~~~~~~~~~~~~~~~~~~~~~~~~~~~~~~~~~~~~~~~~~~~~~~~~~~~~~~~~~~~~~~~~~~~~~~~~~~~~~~~~~~~~~~~~~~~~<1395

              *         *         *         *         *         *         *         *         *         *
1501>aatcatgctgactctgtggcggacccgcgctcttgccggcccggcgataacgctgggcgtgaggctgtgcccggcggagttttttgcgcctgcattttcc>1600
1434<----------------------------------------------------------------------------------------------------<1434
1450<----------------------------------------------------------------------------------------------------<1450
1479<~~~~~~~~~~~~~~~~~~~~~~~~~~~~~~~~~~~~~~~~~~~~~~~~~~~~~~~~~~~~~~~~~~~~~~~~~~~~~~~~~~~~~~~~~~~~~~~~~~~~<1479
1395<~~~~~~~~~~~~~~~~~~~~~~~~~~~~~~~~~~~~~~~~~~~~~~~~~~~~~~~~~~~~~~~~~~~~~~~~~~~~~~~~~~~~~~~~~~~~~~~~~~~~<1395
1388<~~~~~~~~~~~~~~~~~~~~~~~~~~~~~~~~~~~~~~~~~~~~~~~~~~~~~~~~~~~~~~~~~~~~~~~~~~~~~~~~~~~~~~~~~~~~~~~~~~~~<1388
1395<~~~~~~~~~~~~~~~~~~~~~~~~~~~~~~~~~~~~~~~~~~~~~~~~~~~~~~~~~~~~~~~~~~~~~~~~~~~~~~~~~~~~~~~~~~~~~~~~~~~~<1395

              *         *         *         *         *         *         *         *         *         *
1601>aaggtttaccctgcgctaaggggcgagattggagaagcaataagaatgccggttggggttgcgatgatgacgaccacgacaactggtgtcattatttaag>1700
1434<----------------------------------------------------------------------------------------------------<1434
1450<----------------------------------------------------------------------------------------------------<1450
```

```
1479<~~~~~~~~~~~~~~~~~~~~~~~~~~~~~~~~~~~~~~~~~~~~~~~~~~~~~~~~~~~~~~~~~~~~~~~~~~~~~~~~~~~~~~~~~~~~~~~~<1479
1395<~~~~~~~~~~~~~~~~~~~~~~~~~~~~~~~~~~~~~~~~~~~~~~~~~~~~~~~~~~~~~~~~~~~~~~~~~~~~~~~~~~~~~~~~~~~~~~~~<1395
1388<~~~~~~~~~~~~~~~~~~~~~~~~~~~~~~~~~~~~~~~~~~~~~~~~~~~~~~~~~~~~~~~~~~~~~~~~~~~~~~~~~~~~~~~~~~~~~~~~<1388
1395<~~~~~~~~~~~~~~~~~~~~~~~~~~~~~~~~~~~~~~~~~~~~~~~~~~~~~~~~~~~~~~~~~~~~~~~~~~~~~~~~~~~~~~~~~~~~~~~~<1395

              *         *         *         *         *         *         *         *         *         *
1701>ttgccgaaagaacctgagtgcatttgcaacatgagtatactagaagaatgagccaagacttgcgagacgcgagtttgccggtggtgcgaacaatagagcg>1800
1434<----------------------------------------------------------------------------------------------------<1434
1450<----------------------------------------------------------------------------------------------------<1450
1479<~~~~~~~~~~~~~~~~~~~~~~~~~~~~~~~~~~~~~~~~~~~~~~~~~~~~~~~~~~~~~~~~~~~~~~~~~~~~~~~~~~~~~~~~~~~~~~~~<1479
1395<~~~~~~~~~~~~~~~~~~~~~~~~~~~~~~~~~~~~~~~~~~~~~~~~~~~~~~~~~~~~~~~~~~~~~~~~~~~~~~~~~~~~~~~~~~~~~~~~<1395
1388<~~~~~~~~~~~~~~~~~~~~~~~~~~~~~~~~~~~~~~~~~~~~~~~~~~~~~~~~~~~~~~~~~~~~~~~~~~~~~~~~~~~~~~~~~~~~~~~~<1388
1395<~~~~~~~~~~~~~~~~~~~~~~~~~~~~~~~~~~~~~~~~~~~~~~~~~~~~~~~~~~~~~~~~~~~~~~~~~~~~~~~~~~~~~~~~~~~~~~~~<1395

              *         *         *         *         *         *         *         *         *         *
1801>accatgaccttgaaggtgagacgcgcataaccgctagagtactttgaagaggaaacagcaatagggttgctaccagtataaatagacaggtacatacaac>1900
1434<----------------------------------------------------------------------------------------------------<1434
1450<----------------------------------------------------------------------------------------------------<1450
1479<~~~~~~~~~~~~~~~~~~~~~~~~~~~~~~~~~~~~~~~~~~~~~~~~~~~~~~~~~~~~~~~~~~~~~~~~~~~~~~~~~~~~~~~~~~~~~~~~<1479
1395<~~~~~~~~~~~~~~~~~~~~~~~~~~~~~~~~~~~~~~~~~~~~~~~~~~~~~~~~~~~~~~~~~~~~~~~~~~~~~~~~~~~~~~~~~~~~~~~~<1395
1388<~~~~~~~~~~~~~~~~~~~~~~~~~~~~~~~~~~~~~~~~~~~~~~~~~~~~~~~~~~~~~~~~~~~~~~~~~~~~~~~~~~~~~~~~~~~~~~~~<1388
1395<~~~~~~~~~~~~~~~~~~~~~~~~~~~~~~~~~~~~~~~~~~~~~~~~~~~~~~~~~~~~~~~~~~~~~~~~~~~~~~~~~~~~~~~~~~~~~~~~<1395

              *         *         *         *         *         *         *         *         *         *
1901>actggaaatggttgtctgtttgagtacgctttcaattcatttgggtgtgcactttattatgttacaatatggaagggaactttacacttctcctatgcac>2000
1434<----------------------------------------------------------------------------------------------------<1434
1450<----------------------------------------------------------------------------------------------------<1450
1479<~~~~~~~~~~~~~~~~~~~~~~~~~~~~~~~~~~~~~~~~~~~~~~~~~~~~~~~~~~~~~~~~~~~~~~~~~~~~~~~~~~~~~~~~~~~~~~~~<1479
1395<~~~~~~~~~~~~~~~~~~~~~~~~~~~~~~~~~~~~~~~~~~~~~~~~~~~~~~~~~~~~~~~~~~~~~~~~~~~~~~~~~~~~~~~~~~~~~~~~<1395
1388<~~~~~~~~~~~~~~~~~~~~~~~~~~~~~~~~~~~~~~~~~~~~~~~~~~~~~~~~~~~~~~~~~~~~~~~~~~~~~~~~~~~~~~~~~~~~~~~~<1388
1395<~~~~~~~~~~~~~~~~~~~~~~~~~~~~~~~~~~~~~~~~~~~~~~~~~~~~~~~~~~~~~~~~~~~~~~~~~~~~~~~~~~~~~~~~~~~~~~~~<1395

              *         *         *         *         *         *         *         *         *         *
2001>atatattaattaaagtccaatgctagtagagaaggggggtaacacccctccgcgctcttttccgatttttttctaaaccgtggaatatttcggatatcct>2100
1434<----------------------------------------------------------------------------------------------------<1434
1450<----------------------------------------------------------------------------------------------------<1450
1479<~~~~~~~~~~~~~~~~~~~~~~~~~~~~~~~~~~~~~~~~~~~~~~~~~~~~~~~~~~~~~~~~~~~~~~~~~~~~~~~~~~~~~~~~~~~~~~~~<1479
1395<~~~~~~~~~~~~~~~~~~~~~~~~~~~~~~~~~~~~~~~~~~~~~~~~~~~~~~~~~~~~~~~~~~~~~~~~~~~~~~~~~~~~~~~~~~~~~~~~<1395
1388<~~~~~~~~~~~~~~~~~~~~~~~~~~~~~~~~~~~~~~~~~~~~~~~~~~~~~~~~~~~~~~~~~~~~~~~~~~~~~~~~~~~~~~~~~~~~~~~~<1388
```

```
1395<~~~~~~~~~~~~~~~~~~~~~~~~~~~~~~~~~~~~~~~~~~~~~~~~~~~~~~~~~~~~~~~~~~~~~~~~~~~~~~~~~~~~~~~~~~~~~~~~~<1395

              *         *         *         *         *         *         *         *         *         *
2101>tttgttgtttccgggtgtacaatatggacttcctcttttctggcaaccaaacccatacatcgggattcctataataccttcgttggtctccctaacatgt>2200
1434<----------------------------------------------------------------------------------------------------<1434
1450<----------------------------------------------------------------------------------------------------<1450
1479<~~~~~~~~~~~~~~~~~~~~~~~~~~~~~~~~~~~~~~~~~~~~~~~~~~~~~~~~~~~~~~~~~~~~~~~~~~~~~~~~~~~~~~~~~~~~~~~~~~<1479
1395<~~~~~~~~~~~~~~~~~~~~~~~~~~~~~~~~~~~~~~~~~~~~~~~~~~~~~~~~~~~~~~~~~~~~~~~~~~~~~~~~~~~~~~~~~~~~~~~~~<1395
1388<~~~~~~~~~~~~~~~~~~~~~~~~~~~~~~~~~~~~~~~~~~~~~~~~~~~~~~~~~~~~~~~~~~~~~~~~~~~~~~~~~~~~~~~~~~~~~~~~~<1388
1395<~~~~~~~~~~~~~~~~~~~~~~~~~~~~~~~~~~~~~~~~~~~~~~~~~~~~~~~~~~~~~~~~~~~~~~~~~~~~~~~~~~~~~~~~~~~~~~~~~<1395

              *         *         *         *         *         *         *         *         *         *
2201>aggtggcggaggggagatatacaatagaacagataccagacaagacataatgggctaaacaagactacaccaattacactgcctcattgatggtggtaca>2300
1434<----------------------------------------------------------------------------------------------------<1434
1450<----------------------------------------------------------------------------------------------------<1450
1479<~~~~~~~~~~~~~~~~~~~~~~~~~~~~~~~~~~~~~~~~~~~~~~~~~~~~~~~~~~~~~~~~~~~~~~~~~~~~~~~~~~~~~~~~~~~~~~~~~~<1479
1395<~~~~~~~~~~~~~~~~~~~~~~~~~~~~~~~~~~~~~~~~~~~~~~~~~~~~~~~~~~~~~~~~~~~~~~~~~~~~~~~~~~~~~~~~~~~~~~~~~<1395
1388<~~~~~~~~~~~~~~~~~~~~~~~~~~~~~~~~~~~~~~~~~~~~~~~~~~~~~~~~~~~~~~~~~~~~~~~~~~~~~~~~~~~~~~~~~~~~~~~~~<1388
1395<~~~~~~~~~~~~~~~~~~~~~~~~~~~~~~~~~~~~~~~~~~~~~~~~~~~~~~~~~~~~~~~~~~~~~~~~~~~~~~~~~~~~~~~~~~~~~~~~~<1395

              *         *         *         *         *         *         *         *         *         *
2301>taacgaactaatactgtagccctagacttgatagccatcatcatatcgaagtttcactaccctttttccatttgccatctattgaagtaataataggcgc>2400
1434<----------------------------------------------------------------------------------------------------<1434
1450<----------------------------------------------------------------------------------------------------<1450
1479<~~~~~~~~~~~~~~~~~~~~~~~~~~~~~~~~~~~~~~~~~~~~~~~~~~~~~~~~~~~~~~~~~~~~~~~~~~~~~~~~~~~~~~~~~~~~~~~~~~<1479
1395<~~~~~~~~~~~~~~~~~~~~~~~~~~~~~~~~~~~~~~~~~~~~~~~~~~~~~~~~~~~~~~~~~~~~~~~~~~~~~~~~~~~~~~~~~~~~~~~~~<1395
1388<~~~~~~~~~~~~~~~~~~~~~~~~~~~~~~~~~~~~~~~~~~~~~~~~~~~~~~~~~~~~~~~~~~~~~~~~~~~~~~~~~~~~~~~~~~~~~~~~~<1388
1395<~~~~~~~~~~~~~~~~~~~~~~~~~~~~~~~~~~~~~~~~~~~~~~~~~~~~~~~~~~~~~~~~~~~~~~~~~~~~~~~~~~~~~~~~~~~~~~~~~<1395

              *         *         *         *         *         *         *         *         *         *
2401>atgcaacttcttttcttttttttttcttttctctctccccgttgttgtctcaccatatccgcaatgacaaaaaaaatgatggaagacactaaaggaaaaaa>2500
1434<----------------------------------------------------------------------------------------------------<1434
1450<----------------------------------------------------------------------------------------------------<1450
1479<~~~~~~~~~~~~~~~~~~~~~~~~~~~~~~~~~~~~~~~~~~~~~~~~~~~~~~~~~~~~~~~~~~~~~~~~~~~~~~~~~~~~~~~~~~~~~~~~~~<1479
1395<~~~~~~~~~~~~~~~~~~~~~~~~~~~~~~~~~~~~~~~~~~~~~~~~~~~~~~~~~~~~~~~~~~~~~~~~~~~~~~~~~~~~~~~~~~~~~~~~~<1395
1388<~~~~~~~~~~~~~~~~~~~~~~~~~~~~~~~~~~~~~~~~~~~~~~~~~~~~~~~~~~~~~~~~~~~~~~~~~~~~~~~~~~~~~~~~~~~~~~~~~<1388
1395<~~~~~~~~~~~~~~~~~~~~~~~~~~~~~~~~~~~~~~~~~~~~~~~~~~~~~~~~~~~~~~~~~~~~~~~~~~~~~~~~~~~~~~~~~~~~~~~~~<1395
```

```
              *         *         *         *         *         *         *         *         *         *
2501>ttaacgacaaagacagcaccaacagatgtcgttgttccagagctgatgaggggtatctcgaagcacacgaaactttttccttccttcattcacgcacact>2600
1434<----------------------------------------------------------------------------------------------------<1434
1450<----------------------------------------------------------------------------------------------------<1450
1479<~~~~~~~~~~~~~~~~~~~~~~~~~~~~~~~~~~~~~~~~~~~~~~~~~~~~~~~~~~~~~~~~~~~~~~~~~~~~~~~~~~~~~~~~~~~~~~~~~~~~<1479
1395<~~~~~~~~~~~~~~~~~~~~~~~~~~~~~~~~~~~~~~~~~~~~~~~~~~~~~~~~~~~~~~~~~~~~~~~~~~~~~~~~~~~~~~~~~~~~~~~~~~~~<1395
1388<~~~~~~~~~~~~~~~~~~~~~~~~~~~~~~~~~~~~~~~~~~~~~~~~~~~~~~~~~~~~~~~~~~~~~~~~~~~~~~~~~~~~~~~~~~~~~~~~~~~~<1388
1395<~~~~~~~~~~~~~~~~~~~~~~~~~~~~~~~~~~~~~~~~~~~~~~~~~~~~~~~~~~~~~~~~~~~~~~~~~~~~~~~~~~~~~~~~~~~~~~~~~~~~<1395

              *         *         *         *         *         *         *         *         *         *
2601>actctctaatgagcaacggtatacggccttccttccagttacttgaatttgaaataaaaaaaagtttgctgtcttgctatcaagtataaatagacctgca>2700
1434<----------------------------------------------------------------------------------------------------<1434
1450<----------------------------------------------------------------------------------------------------<1450
1479<~~~~~~~~~~~~~~~~~~~~~~~~~~~~~~~~~~~~~~~~~~~~~~~~~~~~~~~~~~~~~~~~~~~~~~~~~~~~~~~~~~~~~~~~~~~~~~~~~~~~<1479
1395<~~~~~~~~~~~~~~~~~~~~~~~~~~~~~~~~~~~~~~~~~~~~~~~~~~~~~~~~~~~~~~~~~~~~~~~~~~~~~~~~~~~~~~~~~~~~~~~~~~~~<1395
1388<~~~~~~~~~~~~~~~~~~~~~~~~~~~~~~~~~~~~~~~~~~~~~~~~~~~~~~~~~~~~~~~~~~~~~~~~~~~~~~~~~~~~~~~~~~~~~~~~~~~~<1388
1395<~~~~~~~~~~~~~~~~~~~~~~~~~~~~~~~~~~~~~~~~~~~~~~~~~~~~~~~~~~~~~~~~~~~~~~~~~~~~~~~~~~~~~~~~~~~~~~~~~~~~<1395

              *         *         *         *         *         *         *         *         *         *
2701>attattaatcttttgtttcctcgtcattgttctcgttccctttcttccttgtttctttttctgcacaatatttcaagctataccaagcatacaatcaact>2800
1434<----------------------------------------------------------------------------------------------------<1434
1450<----------------------------------------------------------------------------------------------------<1450
1479<~~~~~~~~~~~~~~~~~~~~~~~~~~~~~~~~~~~~~~~~~~~~~~~~~~~~~~~~~~~~~~~~~~~~~~~~~~~~~~~~~~~~~~~~~~~~~~~~~~~~<1479
1395<~~~~~~~~~~~~~~~~~~~~~~~~~~~~~~~~~~~~~~~~~~~~~~~~~~~~~~~~~~~~~~~~~~~~~~~~~~~~~~~~~~~~~~~~~~~~~~~~~~~~<1395
1388<~~~~~~~~~~~~~~~~~~~~~~~~~~~~~~~~~~~~~~~~~~~~~~~~~~~~~~~~~~~~~~~~~~~~~~~~~~~~~~~~~~~~~~~~~~~~~~~~~~~~<1388
1395<~~~~~~~~~~~~~~~~~~~~~~~~~~~~~~~~~~~~~~~~~~~~~~~~~~~~~~~~~~~~~~~~~~~~~~~~~~~~~~~~~~~~~~~~~~~~~~~~~~~~<1395

              *         *         *         *         *         *         *         *         *         *
2801>atctcatatacatctagaactagtggatccccccatcacaagtttgtacaaaaaagcaggcttcaaaatgaaagctccttcatcaaatggagtttctccta>2900
1434<----------------------------------------------------------------------------------------------------<1434
1450<----------------------------------------------------------------------------------------------------<1450
1479<~~~~~~~~~~~~~~~~~~~~~~~~~~~~~~~~~~~~~~~~~~~~~~~~~~~~~~~~~~~~~~~~~~~~~~~~~~~~~~~~~~~~~~~~~~~~~~~~~~~~<1479
1395<~~~~~~~~~~~~~~~~~~~~~~~~~~~~~~~~~~~~~~~~~~~~~~~~~~~~~~~~~~~~~~~~~~~~~~~~~~~~~~~~~~~~~~~~~~~~~~~~~~~~<1395
1388<~~~~~~~~~~~~~~~~~~~~~~~~~~~~~~~~~~~~~~~~~~~~~~~~~~~~~~~~~~~~~~~~~~~~~~~~~~~~~~~~~~~~~~~~~~~~~~~~~~~~<1388
1395<~~~~~~~~~~~~~~~~~~~~~~~~~~~~~~~~~~~~~~~~~~~~~~~~~~~~~~~~~~~~~~~~~~~~~~~~~~~~~~~~~~~~~~~~~~~~~~~~~~~~<1395

              *         *         *         *         *         *         *         *         *         *
2901>atcctgttgaaggagaaaggagaaatataaactcagagctatggcacgcttgtgctgggccattgatttcgttgcctccagcaggaagtcttgttgttta>3000
1434<----------------------------------------------------------------------------------------------------<1434
```

```
1450<----------------------------------------------------------------------------------------------------<1450
1479<~~~~~~~~~~~~~~~~~~~~~~~~~~~~~~~~~~~~~~~~~~~~~~~~~~~~~~~~~~~~~~~~~~~~~~~~~~~~~~~~~~~~~~~~~~~~~~~~~~~~<1479
1395<~~~~~~~~~~~~~~~~~~~~~~~~~~~~~~~~~~~~~~~~~~~~~~~~~~~~~~~~~~~~~~~~~~~~~~~~~~~~~~~~~~~~~~~~~~~~~~~~~~~~<1395
1388<~~~~~~~~~~~~~~~~~~~~~~~~~~~~~~~~~~~~~~~~~~~~~~~~~~~~~~~~~~~~~~~~~~~~~~~~~~~~~~~~~~~~~~~~~~~~~~~~~~~~<1388
1395<~~~~~~~~~~~~~~~~~~~~~~~~~~~~~~~~~~~~~~~~~~~~~~~~~~~~~~~~~~~~~~~~~~~~~~~~~~~~~~~~~~~~~~~~~~~~~~~~~~~~<1395

              *         *         *         *         *         *         *         *         *         *
3001>cttccctcaaggtcacagtgagcaagtcgcggcttcaatgcagaagcagactgatttcataccaagttacccgaatcttccttccaagctcatatgcatg>3100
1434<----------------------------------------------------------------------------------------------------<1434
1450<----------------------------------------------------------------------------------------------------<1450
1479<~~~~~~~~~~~~~~~~~~~~~~~~~~~~~~~~~~~~~~~~~~~~~~~~~~~~~~~~~~~~~~~~~~~~~~~~~~~~~~~~~~~~~~~~~~~~~~~~~~~~<1479
1395<~~~~~~~~~~~~~~~~~~~~~~~~~~~~~~~~~~~~~~~~~~~~~~~~~~~~~~~~~~~~~~~~~~~~~~~~~~~~~~~~~~~~~~~~~~~~~~~~~~~~<1395
1388<~~~~~~~~~~~~~~~~~~~~~~~~~~~~~~~~~~~~~~~~~~~~~~~~~~~~~~~~~~~~~~~~~~~~~~~~~~~~~~~~~~~~~~~~~~~~~~~~~~~~<1388
1395<~~~~~~~~~~~~~~~~~~~~~~~~~~~~~~~~~~~~~~~~~~~~~~~~~~~~~~~~~~~~~~~~~~~~~~~~~~~~~~~~~~~~~~~~~~~~~~~~~~~~<1395

              *         *         *         *         *         *         *         *         *         *
3101>ctccacaatgttacactgaatgctgatcctgagacggatgaggtctatgcgcagatgactcttcagccagtaaacaaatatgacagagatgcattgcttg>3200
1434<----------------------------------------------------------------------------------------------------<1434
1450<----------------------------------------------------------------------------------------------------<1450
1479<~~~~~~~~~~~~~~~~~~~~~~~~~~~~~~~~~~~~~~~~~~~~~~~~~~~~~~~~~~~~~~~~~~~~~~~~~~~~~~~~~~~~~~~~~~~~~~~~~~~~<1479
1395<~~~~~~~~~~~~~~~~~~~~~~~~~~~~~~~~~~~~~~~~~~~~~~~~~~~~~~~~~~~~~~~~~~~~~~~~~~~~~~~~~~~~~~~~~~~~~~~~~~~~<1395
1388<~~~~~~~~~~~~~~~~~~~~~~~~~~~~~~~~~~~~~~~~~~~~~~~~~~~~~~~~~~~~~~~~~~~~~~~~~~~~~~~~~~~~~~~~~~~~~~~~~~~~<1388
1395<~~~~~~~~~~~~~~~~~~~~~~~~~~~~~~~~~~~~~~~~~~~~~~~~~~~~~~~~~~~~~~~~~~~~~~~~~~~~~~~~~~~~~~~~~~~~~~~~~~~~<1395

              *         *         *         *         *         *         *         *         *         *
3201>cttctgacatgggtcttaagctaaacagacaacctaatgaatttttctgcaaaaccctcacggcgagtgacacaagtactcacggtggattttctgtacc>3300
1434<----------------------------------------------------------------------------------------------------<1434
1450<----------------------------------------------------------------------------------------------------<1450
1479<~~~~~~~~~~~~~~~~~~~~~~~~~~~~~~~~~~~~~~~~~~~~~~~~~~~~~~~~~~~~~~~~~~~~~~~~~~~~~~~~~~~~~~~~~~~~~~~~~~~~<1479
1395<~~~~~~~~~~~~~~~~~~~~~~~~~~~~~~~~~~~~~~~~~~~~~~~~~~~~~~~~~~~~~~~~~~~~~~~~~~~~~~~~~~~~~~~~~~~~~~~~~~~~<1395
1388<~~~~~~~~~~~~~~~~~~~~~~~~~~~~~~~~~~~~~~~~~~~~~~~~~~~~~~~~~~~~~~~~~~~~~~~~~~~~~~~~~~~~~~~~~~~~~~~~~~~~<1388
1395<~~~~~~~~~~~~~~~~~~~~~~~~~~~~~~~~~~~~~~~~~~~~~~~~~~~~~~~~~~~~~~~~~~~~~~~~~~~~~~~~~~~~~~~~~~~~~~~~~~~~<1395

              *         *         *         *         *         *         *         *         *         *
3301>ccgacgagctgctgagaaaatctttcctgctctggatttctcgatgcaaccaccttgtcaggagcttgttgctaaggatattcatgacaacacatggact>3400
1434<----------------------------------------------------------------------------------------------------<1434
1450<----------------------------------------------------------------------------------------------------<1450
1479<~~~~~~~~~~~~~~~~~~~~~~~~~~~~~~~~~~~~~~~~~~~~~~~~~~~~~~~~~~~~~~~~~~~~~~~~~~~~~~~~~~~~~~~~~~~~~~~~~~~~<1479
1395<~~~~~~~~~~~~~~~~~~~~~~~~~~~~~~~~~~~~~~~~~~~~~~~~~~~~~~~~~~~~~~~~~~~~~~~~~~~~~~~~~~~~~~~~~~~~~~~~~~~~<1395
```

```
1388<~~~~~~~~~~~~~~~~~~~~~~~~~~~~~~~~~~~~~~~~~~~~~~~~~~~~~~~~~~~~~~~~~~~~~~~~~~~~~~~~~~~~~~~~~~~~~~~~~<1388
1395<~~~~~~~~~~~~~~~~~~~~~~~~~~~~~~~~~~~~~~~~~~~~~~~~~~~~~~~~~~~~~~~~~~~~~~~~~~~~~~~~~~~~~~~~~~~~~~~~~<1395

              *         *         *         *         *         *         *         *         *         *
3401>ttcagacatatttatcgaggtcaaccaaaaaggcacttgctaactacaggctggagtgtgtttgtcagcacgaaaaggctctttgctggagactctgttc>3500
1434<----------------------------------------------------------------------------------------------------<1434
1450<----------------------------------------------------------------------------------------------------<1450
1479<~~~~~~~~~~~~~~~~~~~~~~~~~~~~~~~~~~~~~~~~~~~~~~~~~~~~~~~~~~~~~~~~~~~~~~~~~~~~~~~~~~~~~~~~~~~~~~~~~<1479
1395<~~~~~~~~~~~~~~~~~~~~~~~~~~~~~~~~~~~~~~~~~~~~~~~~~~~~~~~~~~~~~~~~~~~~~~~~~~~~~~~~~~~~~~~~~~~~~~~~~<1395
1388<~~~~~~~~~~~~~~~~~~~~~~~~~~~~~~~~~~~~~~~~~~~~~~~~~~~~~~~~~~~~~~~~~~~~~~~~~~~~~~~~~~~~~~~~~~~~~~~~~<1388
1395<~~~~~~~~~~~~~~~~~~~~~~~~~~~~~~~~~~~~~~~~~~~~~~~~~~~~~~~~~~~~~~~~~~~~~~~~~~~~~~~~~~~~~~~~~~~~~~~~~<1395

              *         *         *         *         *         *         *         *         *         *
3501>tttttataagagatggaaaggcgcaacttctgttggggataagacgtgcaaatagacaacagcctgcactttcttcatctgtaatatcaagtgatagcat>3600
1434<----------------------------------------------------------------------------------------------------<1434
1450<----------------------------------------------------------------------------------------------------<1450
1479<~~~~~~~~~~~~~~~~~~~~~~~~~~~~~~~~~~~~~~~~~~~~~~~~~~~~~~~~~~~~~~~~~~~~~~~~~~~~~~~~~~~~~~~~~~~~~~~~~<1479
1395<~~~~~~~~~~~~~~~~~~~~~~~~~~~~~~~~~~~~~~~~~~~~~~~~~~~~~~~~~~~~~~~~~~~~~~~~~~~~~~~~~~~~~~~~~~~~~~~~~<1395
1388<~~~~~~~~~~~~~~~~~~~~~~~~~~~~~~~~~~~~~~~~~~~~~~~~~~~~~~~~~~~~~~~~~~~~~~~~~~~~~~~~~~~~~~~~~~~~~~~~~<1388
1395<~~~~~~~~~~~~~~~~~~~~~~~~~~~~~~~~~~~~~~~~~~~~~~~~~~~~~~~~~~~~~~~~~~~~~~~~~~~~~~~~~~~~~~~~~~~~~~~~~<1395

              *         *         *         *         *         *         *         *         *         *
3601>gcacatcggagttcttgcagctgcagctcatgctaatgctaataacagtcctttcaccattttctacaacccgaggtgggctgctcctgctgagtttgtg>3700
1434<----------------------------------------------------------------------------------------------------<1434
1450<----------------------------------------------------------------------------------------------------<1450
1479<~~~~~~~~~~~~~~~~~~~~~~~~~~~~~~~~~~~~~~~~~~~~~~~~~~~~~~~~~~~~~~~~~~~~~~~~~~~~~~~~~~~~~~~~~~~~~~~~~<1479
1395<~~~~~~~~~~~~~~~~~~~~~~~~~~~~~~~~~~~~~~~~~~~~~~~~~~~~~~~~~~~~~~~~~~~~~~~~~~~~~~~~~~~~~~~~~~~~~~~~~<1395
1388<~~~~~~~~~~~~~~~~~~~~~~~~~~~~~~~~~~~~~~~~~~~~~~~~~~~~~~~~~~~~~~~~~~~~~~~~~~~~~~~~~~~~~~~~~~~~~~~~~<1388
1395<~~~~~~~~~~~~~~~~~~~~~~~~~~~~~~~~~~~~~~~~~~~~~~~~~~~~~~~~~~~~~~~~~~~~~~~~~~~~~~~~~~~~~~~~~~~~~~~~~<1395

              *         *         *         *         *         *         *         *         *         *
3701>gttcctttagccaagtataccaaagcgatgtacgctcaagtttccctcggtatgcggtttagaatgatatttgagactgaagaatgtggagttcgtcggt>3800
1434<----------------------------------------------------------------------------------------------------<1434
1450<----------------------------------------------------------------------------------------------------<1450
1479<~~~~~~~~~~~~~~~~~~~~~~~~~~~~~~~~~~~~~~~~~~~~~~~~~~~~~~~~~~~~~~~~~~~~~~~~~~~~~~~~~~~~~~~~~~~~~~~~~<1479
1395<~~~~~~~~~~~~~~~~~~~~~~~~~~~~~~~~~~~~~~~~~~~~~~~~~~~~~~~~~~~~~~~~~~~~~~~~~~~~~~~~~~~~~~~~~~~~~~~~~<1395
1388<~~~~~~~~~~~~~~~~~~~~~~~~~~~~~~~~~~~~~~~~~~~~~~~~~~~~~~~~~~~~~~~~~~~~~~~~~~~~~~~~~~~~~~~~~~~~~~~~~<1388
1395<~~~~~~~~~~~~~~~~~~~~~~~~~~~~~~~~~~~~~~~~~~~~~~~~~~~~~~~~~~~~~~~~~~~~~~~~~~~~~~~~~~~~~~~~~~~~~~~~~<1395
```

```
              *         *         *         *         *         *         *         *         *         *
3801>atatgggtacagttaccggtatcagtgatcttgatccagtgagatggaaaaactctcagtggcggaatcttcagattggatgggatgagtcagctgctgg>3900
1434<----------------------------------------------------------------------------------------------------<1434
1450<----------------------------------------------------------------------------------------------------<1450
1479<~~~~~~~~~~~~~~~~~~~~~~~~~~~~~~~~~~~~~~~~~~~~~~~~~~~~~~~~~~~~~~~~~~~~~~~~~~~~~~~~~~~~~~~~~~~~~~~~~~<1479
1395<~~~~~~~~~~~~~~~~~~~~~~~~~~~~~~~~~~~~~~~~~~~~~~~~~~~~~~~~~~~~~~~~~~~~~~~~~~~~~~~~~~~~~~~~~~~~~~~~~~<1395
1388<~~~~~~~~~~~~~~~~~~~~~~~~~~~~~~~~~~~~~~~~~~~~~~~~~~~~~~~~~~~~~~~~~~~~~~~~~~~~~~~~~~~~~~~~~~~~~~~~~~<1388
1395<~~~~~~~~~~~~~~~~~~~~~~~~~~~~~~~~~~~~~~~~~~~~~~~~~~~~~~~~~~~~~~~~~~~~~~~~~~~~~~~~~~~~~~~~~~~~~~~~~~<1395

              *         *         *         *         *         *         *         *         *         *
3901>tgataggcccagtcgagtttcagtttgggacattgaaccggttttaactcctttctacatatgtcctcctccatttttccgacctcgcttttctggacaa>4000
1434<----------------------------------------------------------------------------------------------------<1434
1450<----------------------------------------------------------------------------------------------------<1450
1479<~~~~~~~~~~~~~~~~~~~~~~~~~~~~~~~~~~~~~~~~~~~~~~~~~~~~~~~~~~~~~~~~~~~~~~~~~~~~~~~~~~~~~~~~~~~~~~~~~~<1479
1395<~~~~~~~~~~~~~~~~~~~~~~~~~~~~~~~~~~~~~~~~~~~~~~~~~~~~~~~~~~~~~~~~~~~~~~~~~~~~~~~~~~~~~~~~~~~~~~~~~~<1395
1388<~~~~~~~~~~~~~~~~~~~~~~~~~~~~~~~~~~~~~~~~~~~~~~~~~~~~~~~~~~~~~~~~~~~~~~~~~~~~~~~~~~~~~~~~~~~~~~~~~~<1388
1395<~~~~~~~~~~~~~~~~~~~~~~~~~~~~~~~~~~~~~~~~~~~~~~~~~~~~~~~~~~~~~~~~~~~~~~~~~~~~~~~~~~~~~~~~~~~~~~~~~~<1395

              *         *         *         *         *         *         *         *         *         *
4001>cctggaatgccagatgatgagactgacatggagtctgcactgaagagagcaatgccatggcttgataatagcttagagatgaaagacccttcgagtacta>4100
1434<----------------------------------------------------------------------------------------------------<1434
1450<----------------------------------------------------------------------------------------------------<1450
1479<~~~~~~~~~~~~~~~~~~~~~~~~~~~~~~~~~~~~~~~~~~~~~~~~~~~~~~~~~~~~~~~~~~~~~~~~~~~~~~~~~~~~~~~~~~~~~~~~~~<1479
1395<~~~~~~~~~~~~~~~~~~~~~~~~~~~~~~~~~~~~~~~~~~~~~~~~~~~~~~~~~~~~~~~~~~~~~~~~~~~~~~~~~~~~~~~~~~~~~~~~~~<1395
1388<~~~~~~~~~~~~~~~~~~~~~~~~~~~~~~~~~~~~~~~~~~~~~~~~~~~~~~~~~~~~~~~~~~~~~~~~~~~~~~~~~~~~~~~~~~~~~~~~~~<1388
1395<~~~~~~~~~~~~~~~~~~~~~~~~~~~~~~~~~~~~~~~~~~~~~~~~~~~~~~~~~~~~~~~~~~~~~~~~~~~~~~~~~~~~~~~~~~~~~~~~~~<1395

              *         *         *         *         *         *         *         *         *         *
4101>tctttcctggtctgagtttagttcagtggatgaatatgcagcagcagaacggccagctaccctctgctgctgcacagccaggtttcttcccatcaatgct>4200
1434<----------------------------------------------------------------------------------------------------<1434
1450<----------------------------------------------------------------------------------------------------<1450
1479<~~~~~~~~~~~~~~~~~~~~~~~~~~~~~~~~~~~~~~~~~~~~~~~~~~~~~~~~~~~~~~~~~~~~~~~~~~~~~~~~~~~~~~~~~~~~~~~~~~<1479
1395<~~~~~~~~~~~~~~~~~~~~~~~~~~~~~~~~~~~~~~~~~~~~~~~~~~~~~~~~~~~~~~~~~~~~~~~~~~~~~~~~~~~~~~~~~~~~~~~~~~<1395
1388<~~~~~~~~~~~~~~~~~~~~~~~~~~~~~~~~~~~~~~~~~~~~~~~~~~~~~~~~~~~~~~~~~~~~~~~~~~~~~~~~~~~~~~~~~~~~~~~~~~<1388
1395<~~~~~~~~~~~~~~~~~~~~~~~~~~~~~~~~~~~~~~~~~~~~~~~~~~~~~~~~~~~~~~~~~~~~~~~~~~~~~~~~~~~~~~~~~~~~~~~~~~<1395

              *         *         *         *         *         *         *         *         *         *
4201>ttcgccaaccgcggcgctgcacaacaatcttggcggcactgatgatccctccaagttactgagctttcagacgccgcacgggggatttcctcctcaaat>4300
```

```
1434<----------------------------------------------------------------------------------------------------<1434
1450<----------------------------------------------------------------------------------------------------<1450
1479<~~~~~~~~~~~~~~~~~~~~~~~~~~~~~~~~~~~~~~~~~~~~~~~~~~~~~~~~~~~~~~~~~~~~~~~~~~~~~~~~~~~~~~~~~~~~~~~~~~~~<1479
1395<~~~~~~~~~~~~~~~~~~~~~~~~~~~~~~~~~~~~~~~~~~~~~~~~~~~~~~~~~~~~~~~~~~~~~~~~~~~~~~~~~~~~~~~~~~~~~~~~~~~~<1395
1388<~~~~~~~~~~~~~~~~~~~~~~~~~~~~~~~~~~~~~~~~~~~~~~~~~~~~~~~~~~~~~~~~~~~~~~~~~~~~~~~~~~~~~~~~~~~~~~~~~~~~<1388
1395<~~~~~~~~~~~~~~~~~~~~~~~~~~~~~~~~~~~~~~~~~~~~~~~~~~~~~~~~~~~~~~~~~~~~~~~~~~~~~~~~~~~~~~~~~~~~~~~~~~~~<1395

              *         *         *         *         *         *         *         *         *         *
4301>ctccaatttaacaaacagaatcagcaagccccaatgtctcagttgcctcagccaccaactacgttgtcccaacaacagcagctgcagcaattgttgcact>4400
1434<----------------------------------------------------------------------------------------------------<1434
1450<----------------------------------------------------------------------------------------------------<1450
1479<~~~~~~~~~~~~~~~~~~~~~~~~~~~~~~~~~~~~~~~~~~~~~~~~~~~~~~~~~~~~~~~~~~~~~~~~~~~~~~~~~~~~~~~~~~~~~~~~~~~~<1479
1395<~~~~~~~~~~~~~~~~~~~~~~~~~~~~~~~~~~~~~~~~~~~~~~~~~~~~~~~~~~~~~~~~~~~~~~~~~~~~~~~~~~~~~~~~~~~~~~~~~~~~<1395
1388<~~~~~~~~~~~~~~~~~~~~~~~~~~~~~~~~~~~~~~~~~~~~~~~~~~~~~~~~~~~~~~~~~~~~~~~~~~~~~~~~~~~~~~~~~~~~~~~~~~~~<1388
1395<~~~~~~~~~~~~~~~~~~~~~~~~~~~~~~~~~~~~~~~~~~~~~~~~~~~~~~~~~~~~~~~~~~~~~~~~~~~~~~~~~~~~~~~~~~~~~~~~~~~~<1395

              *         *         *         *         *         *         *         *         *         *
4401>cctctttgaaccatcaacaacagcaatcgcagtctcaacaacagcaacaacaacaacagttgctgcagcagcaacaacaattgcagtctcaacaacacag>4500
1434<----------------------------------------------------------------------------------------------------<1434
1450<----------------------------------------------------------------------------------------------------<1450
1479<~~~~~~~~~~~~~~~~~~~~~~~~~~~~~~~~~~~~~~~~~~~~~~~~~~~~~~~~~~~~~~~~~~~~~~~~~~~~~~~~~~~~~~~~~~~~~~~~~~~~<1479
1395<~~~~~~~~~~~~~~~~~~~~~~~~~~~~~~~~~~~~~~~~~~~~~~~~~~~~~~~~~~~~~~~~~~~~~~~~~~~~~~~~~~~~~~~~~~~~~~~~~~~~<1395
1388<~~~~~~~~~~~~~~~~~~~~~~~~~~~~~~~~~~~~~~~~~~~~~~~~~~~~~~~~~~~~~~~~~~~~~~~~~~~~~~~~~~~~~~~~~~~~~~~~~~~~<1388
1395<~~~~~~~~~~~~~~~~~~~~~~~~~~~~~~~~~~~~~~~~~~~~~~~~~~~~~~~~~~~~~~~~~~~~~~~~~~~~~~~~~~~~~~~~~~~~~~~~~~~~<1395

              *         *         *         *         *         *         *         *         *         *
4501>caacaacaatcaatcgcagtctcagcaacaacaacaattgctccagcagcaacaacaacaacaactgcagcaacaacatcaacaaccgttacagcaacag>4600
1434<----------------------------------------------------------------------------------------------------<1434
1450<----------------------------------------------------------------------------------------------------<1450
1479<~~~~~~~~~~~~~~~~~~~~~~~~~~~~~~~~~~~~~~~~~~~~~~~~~~~~~~~~~~~~~~~~~~~~~~~~~~~~~~~~~~~~~~~~~~~~~~~~~~~~<1479
1395<~~~~~~~~~~~~~~~~~~~~~~~~~~~~~~~~~~~~~~~~~~~~~~~~~~~~~~~~~~~~~~~~~~~~~~~~~~~~~~~~~~~~~~~~~~~~~~~~~~~~<1395
1388<~~~~~~~~~~~~~~~~~~~~~~~~~~~~~~~~~~~~~~~~~~~~~~~~~~~~~~~~~~~~~~~~~~~~~~~~~~~~~~~~~~~~~~~~~~~~~~~~~~~~<1388
1395<~~~~~~~~~~~~~~~~~~~~~~~~~~~~~~~~~~~~~~~~~~~~~~~~~~~~~~~~~~~~~~~~~~~~~~~~~~~~~~~~~~~~~~~~~~~~~~~~~~~~<1395

              *         *         *         *         *         *         *         *         *         *
4601>actcagcagcagcagctaagaacacagccattgcaatctcactcgcatccacagccacaacagttacaacaacataagttgcagcaacttcaggttccac>4700
1434<----------------------------------------------------------------------------------------------------<1434
1450<----------------------------------------------------------------------------------------------------<1450
1479<~~~~~~~~~~~~~~~~~~~~~~~~~~~~~~~~~~~~~~~~~~~~~~~~~~~~~~~~~~~~~~~~~~~~~~~~~~~~~~~~~~~~~~~~~~~~~~~~~~~~<1479
```

```
1395<~~~~~~~~~~~~~~~~~~~~~~~~~~~~~~~~~~~~~~~~~~~~~~~~~~~~~~~~~~~~~~~~~~~~~~~~~~~~~~~~~~~~~~~~~~~~~~~~~~~~<1395
1388<~~~~~~~~~~~~~~~~~~~~~~~~~~~~~~~~~~~~~~~~~~~~~~~~~~~~~~~~~~~~~~~~~~~~~~~~~~~~~~~~~~~~~~~~~~~~~~~~~~~~<1388
1395<~~~~~~~~~~~~~~~~~~~~~~~~~~~~~~~~~~~~~~~~~~~~~~~~~~~~~~~~~~~~~~~~~~~~~~~~~~~~~~~~~~~~~~~~~~~~~~~~~~~~<1395

              *         *         *         *         *         *         *         *         *         *
4701>agaatcagctttacaatggtcaacaagcagcgcagcagcatcagtcgcaacaagcatctacacatcatttgcaaccacaattagtttcgggatcaatggc>4800
1434<----------------------------------------------------------------------------------------------------<1434
1450<----------------------------------------------------------------------------------------------------<1450
1479<~~~~~~~~~~~~~~~~~~~~~~~~~~~~~~~~~~~~~~~~~~~~~~~~~~~~~~~~~~~~~~~~~~~~~~~~~~~~~~~~~~~~~~~~~~~~~~~~~~~~<1479
1395<~~~~~~~~~~~~~~~~~~~~~~~~~~~~~~~~~~~~~~~~~~~~~~~~~~~~~~~~~~~~~~~~~~~~~~~~~~~~~~~~~~~~~~~~~~~~~~~~~~~~<1395
1388<~~~~~~~~~~~~~~~~~~~~~~~~~~~~~~~~~~~~~~~~~~~~~~~~~~~~~~~~~~~~~~~~~~~~~~~~~~~~~~~~~~~~~~~~~~~~~~~~~~~~<1388
1395<~~~~~~~~~~~~~~~~~~~~~~~~~~~~~~~~~~~~~~~~~~~~~~~~~~~~~~~~~~~~~~~~~~~~~~~~~~~~~~~~~~~~~~~~~~~~~~~~~~~~<1395

              *         *         *         *         *         *         *         *         *         *
4801>aagcagtgtcatcacgcctccgtccagctcccttaatcaaagctttcaacagcaacaacaacagtctaagcaacttcaacaagcacatcaccatttaggt>4900
1434<----------------------------------------------------------------------------------------------------<1434
1450<----------------------------------------------------------------------------------------------------<1450
1479<~~~~~~~~~~~~~~~~~~~~~~~~~~~~~~~~~~~~~~~~~~~~~~~~~~~~~~~~~~~~~~~~~~~~~~~~~~~~~~~~~~~~~~~~~~~~~~~~~~~~<1479
1395<~~~~~~~~~~~~~~~~~~~~~~~~~~~~~~~~~~~~~~~~~~~~~~~~~~~~~~~~~~~~~~~~~~~~~~~~~~~~~~~~~~~~~~~~~~~~~~~~~~~~<1395
1388<~~~~~~~~~~~~~~~~~~~~~~~~~~~~~~~~~~~~~~~~~~~~~~~~~~~~~~~~~~~~~~~~~~~~~~~~~~~~~~~~~~~~~~~~~~~~~~~~~~~~<1388
1395<~~~~~~~~~~~~~~~~~~~~~~~~~~~~~~~~~~~~~~~~~~~~~~~~~~~~~~~~~~~~~~~~~~~~~~~~~~~~~~~~~~~~~~~~~~~~~~~~~~~~<1395

              *         *         *         *         *         *         *         *         *         *
4901>gctagcactagccagagtagtgtaattgaaaccagcaagtcttcatccaatctgatgtccgcaccgccgcaagagacacagttttcacgacaagtagaac>5000
1434<----------------------------------------------------------------------------------------------------<1434
1450<----------------------------------------------------------------------------------------------------<1450
1479<~~~~~~~~~~~~~~~~~~~~~~~~~~~~~~~~~~~~~~~~~~~~~~~~~~~~~~~~~~~~~~~~~~~~~~~~~~~~~~~~~~~~~~~~~~~~~~~~~~~~<1479
1395<~~~~~~~~~~~~~~~~~~~~~~~~~~~~~~~~~~~~~~~~~~~~~~~~~~~~~~~~~~~~~~~~~~~~~~~~~~~~~~~~~~~~~~~~~~~~~~~~~~~~<1395
1388<~~~~~~~~~~~~~~~~~~~~~~~~~~~~~~~~~~~~~~~~~~~~~~~~~~~~~~~~~~~~~~~~~~~~~~~~~~~~~~~~~~~~~~~~~~~~~~~~~~~~<1388
1395<~~~~~~~~~~~~~~~~~~~~~~~~~~~~~~~~~~~~~~~~~~~~~~~~~~~~~~~~~~~~~~~~~~~~~~~~~~~~~~~~~~~~~~~~~~~~~~~~~~~~<1395

              *         *         *         *         *         *         *         *         *         *
5001>agcagcagcctcctggtctcaacgggcagaatcagcaaacacttttgcagcagaaagctcaccaggcacaggcccaacagatattccagcagagtctctt>5100
1434<----------------------------------------------------------------------------------------------------<1434
1450<----------------------------------------------------------------------------------------------------<1450
1479<~~~~~~~~~~~~~~~~~~~~~~~~~~~~~~~~~~~~~~~~~~~~~~~~~~~~~~~~~~~~~~~~~~~~~~~~~~~~~~~~~~~~~~~~~~~~~~~~~~~~<1479
1395<~~~~~~~~~~~~~~~~~~~~~~~~~~~~~~~~~~~~~~~~~~~~~~~~~~~~~~~~~~~~~~~~~~~~~~~~~~~~~~~~~~~~~~~~~~~~~~~~~~~~<1395
1388<~~~~~~~~~~~~~~~~~~~~~~~~~~~~~~~~~~~~~~~~~~~~~~~~~~~~~~~~~~~~~~~~~~~~~~~~~~~~~~~~~~~~~~~~~~~~~~~~~~~~<1388
1395<~~~~~~~~~~~~~~~~~~~~~~~~~~~~~~~~~~~~~~~~~~~~~~~~~~~~~~~~~~~~~~~~~~~~~~~~~~~~~~~~~~~~~~~~~~~~~~~~~~~~<1395
```

```
          *         *         *         *         *         *         *         *         *         *
5101>ggaacagccgcatatacagtttcagctgttacagagattacaacagcaacagcagcaatttctttcgccgcagtctcagttaccacaccatcaattg>5200
1434<----------------------------------------------------------------------------------------------------<1434
1449<-----------ACACGCAGTCTGTTACCAGAGAGATTTCAACGGCAAACGGCAGGCAGCATTCTCGCCGCCAGTTTCAAGTTTAAGCACCCATCAATTGC<1361
1479<~~~~~~~~~~~~~~~~~~~~~~~~~~~~~~~~~~~~~~~~~~~~~~~~~~~~~~~~~~~~~~~~~~~~~~~~~~~~~~~~~~~~~~~~~~~~~~~~~~~~<1479
1395<~~~~~~~~~~~~~~~~~~~~~~~~~~~~~~~~~~~~~~~~~~~~~~~~~~~~~~~~~~~~~~~~~~~~~~~~~~~~~~~~~~~~~~~~~~~~~~~~~~~~<1395
1387<~~~~~~~~~~~~~~~~~~~~~~~~~~~~~~~~~~~~~~~~~~~~~~~~~~~~~~~~~~~~~~~~~~~~~~GGCAGCAATTTCTTGACCGCAAGTTCCAGT<1358
1395<~~~~~~~~~~~~~~~~~~~~~~~~~~~~~~~~~~~~~~~~~~~~~~~~~~~~~~~~~~~~~~~~~~~~~~~~~~~~~~~~~~~~~~~~~~~~~~~~~~~~<1395

          *         *         *         *         *         *         *         *         *
5201>caaagccagcagttgcaacagctgcctactctctctcaaggtcatcagtttccgtcatcttgcactaacaatggcttatcgacgttgcaaccacct-caa>5299
1433<----------------------------------------------------------------------GCCACACCACCTCTCAAATGTCGTGT-TGT<1405
1360<CAAGCCAGCAGTTGCAACCAGCTGCTACTTTTCTCTCAAAGGTCATCAGTTCGTCATTCTTGCACTAACAATGGCTTATCGACGT-GCAGCCACCTCCAA<1262
1479<~~~~~~~~~~~~~~~~~~~~~~~~~~~~~~~~~~~~~~~~~~~~~~~~~~~~~~~~~~~~~~~~~~~~~~~~~~~~~~~~~~~~~~~~~~~~~~~~~~~~<1479
1395<~~~~~~~~~~~~~~~~~~~~~~~~~~~~~~~~~~~~~~~~~~~~~~~~~~~~~~~~~~~~~~~~~~~~~~~~~~~~~~~~~~~~~~~~~~~~~~~~~~~~<1395
1357<TACCAACCATTCATTGCAAGGCAGCAGTGCAACAGGTGCCTATCTTTCTTCCAGGTCATCAGTTCGTCATCTGCACTAACAATGCTATCGACGTGC-AAT<1259
1395<~~~~~~~~~~~~~~~~~~~~~~~~~~~~~~~~~~~~~~~~~~~~~~~~~~~~~~~~~~~~~~~~~~~~~~~~~~~~~~~~~~~~~~~~~~~~~~~~~~~~<1395

*          *         *         *         *         *         *         *         *         *
5300>atgctggt-gagccgacctcaggaaaaacaaaacccaccggttgggggaggggtcaaagcttattcaggcatcacagatggaggagatgcaccttcctct>5398
1404<AGAGCGCG-ACCTCTCAGGAAAAAACAAAACCCACCCGGTTGGGGGAGGGTCAAGCTTATTCAGCATCACCAGAATTGAGGGAGAGATGGCACCTCCTCT<1306
1261<ATGCTGGTGGAGCCGA-CTCA-GAAAAACAAAACCCACCGGTTGGGGAGGGGTCAAAGCTTATTCAGGCATCACAGATG-AGGAGATGCACCTTCCTC-<1166
1478<~~~~~~~~~~~~~~~~~~~~~~~~~~~~~~~~~~~~~~~~~~~~~~~~~~~~~~~~~~~~~~~~~~~~~~~~~~~~AACAAATAGTTGGCGCCTCTTGCA<1455
1394<~~~~~~~~~~~~~~~~~CGAAAAACGGTAATTCAGTATCGAAGCGGGTGTCGTGGCTGTCAACAAGGCTGATCAAGTCTTGACATATCTCATCAGATAGC<1312
1258<CACTCAAT-GCTGTGAGCGACTCAGAAAAAACAAAATCCACCGTTGGGGAGGGGGTCAAGCTATCAGCCATCACAGATGGAGGAGATGCAC--T-CTC-<1164
1394<~~~~~~~~~~~~~~~~~GTATCAACATTCCACAGTGTGTTATTGAGGGAAGTATCAGTACTCGAAACCGTGCTCGTGCCTTACAGCTGATCAGTTCTGAC<1312

 *         *         *         *         *         *         *         *         *         *
5399>tcaacgtcgccttccaccaacaactgtcagatctcttcttcaggctttctcaacagaagccaaagcgggccagcgatcttgatacctgatgcagcgattg>5498
1305<TCACGTCGCCTTCCACCAACAACTGTCAGAATCTCTTCTTCAGGCTTTCTCAACAGAAGCCAAAGCGG-CCAGCGATC-TGATA-CTGATGCAGCGATTG<1209
1165<TCAACGTCGCCTTCCACCAACAACTGTCAGATCTCTTCTTCAGGCTTTCTCAACAGAAGCCAAAGCGGGCCAGCGATCTTGATACCTGATGCAGCGATTG<1066
1454<CACACAACAACCGTCAGAATTCTCTTTTCAAGCTTTCACAAACAGGAGGCCAAAGCGGCAAGCGATTCTTGAATACCTGAGTGGCAGGCGGATGATATGT<1355
1311<CAGCACAGTCTGATGCAATTAATTGAATTCTCTTTTGCTGGAACTGATGAGGCTGACATCACCAAGTTCTCTTCTGCAGTCTTCTGCGATCATACAAGCA<1212
1163<TCAACGTCGCCTTCCACCAACAACTGTCAGATCTCTTCT-CAGGCTTTCTCAACAGAAGCCAAAGCGGGCCAGCGATCT-GATACCTGATGCAGCGATTG<1066
1311<CATACTCATCAGATACACCACAGTCTGATCAATAAATTGATCTCTTCTGAAACTGATGAGGCTGACATCACAGTCTCTCTGCAGTCTCTGCCGATCATAC<1212

 *          *         *         *         *         *         *         *         *         *
```

```
5499>atatg-tctggtaatcttgttcaggatctttacagcaaatccgatatgcggctaaaacaagaactcgtgggtcagcaaaagtccaaagctagtttaacag>5597
1208<ATATGTTCTGGTAATC-TG-TCAGGATC-TTACAGCAAATCCGATATGCGGCTAAAACAAGAACTCGTGGGTCAGCAAAAGTCCAAAGCTAGTTTAACAG<1112
1065<ATATG-TCTGGTAATCTTGTTCAGGATCTTTACAGCAAATCCGATATGCGGCTAAAACAAGAACTCGTGGGTCAGCAAAAGTCCAAAGCTAGTTTAACAG<967
1354<[redacted]-[redacted]CGTGGGTCAGCAAAAGTCCAAAGCTAGTT-AACAG<1257
1211<[redacted]-[redacted]<1113
1065<ATATG-TCTGGTAATCTTGTTCAGGATCTTTACAGCAAATCCGATATGCGGCTAAAACAAGAACTCGTGGGTCAGCAAAAGTCCAAAGCTAGTTTAACAG<967
1211<[redacted]-[redacted]<1113

         *         *         *         *         *         *         *         *         *         *
5598>atcatcaactagaagcatctgcctc-tggaacttcttacggtttagatggaggcgaaaacaac-agacaacaaaatttcttggctccaacttttggcctt>5695
1111<ATCATCAACTAGAAGCATCTGCCTC-TGGAACTTCTTACGGTTTAGATGGAGGCGAAAACAAC-AGACAACAAAATTTCTTGGCTCCAACTTTTGGCCTT<1014
 966<ATCATCAACTAGAAGCATCTGCCTC-TGGAACTTCTTACGGTTTAGATGGAGGCGAAAACAAC-AGACAACAAAATTTCTTGGCTCCAACTTTTGGCCTT<869
1256<ATCATCAACTAGAAGCATCTGCCTCCTGGAACTTCTTACGGTTTAGATG-A-GCGAAAACAACAAGACAACAAAATTTCTTGGCTCCAACTTTTGGCCTT<1159
1112<[redacted]-[redacted]-[redacted]<1015
 966<ATCATCAACTAGAAGCATCTGCCTC-TGGAACTTCTTACGGTTTAGATGGAGGCGAAAACAAC-AGACAACAAAATTTCTTGGCTCCAACTTTTGGCCTT<869
1112<[redacted]-[redacted]-[redacted]<1015

         *         *         *         *         *         *         *         *         *         *
5696>gacggtgattccaggaacagcttgctcggtggagctaatgttgataatggctttgtgcctgacacgctactctcgaggggatatgactcccagaaagatc>5795
1013<GACGGTGATTCCAGGAACAGCTTGCTCGGTGGAGCTAATGTTGATAATGGCTTTGTGCCTGACACGCTACTCTCGAGGGGATATGACTCCCAGAAAGATC<914
 868<GACGGTGATTCCAGGAACAGCTTGCTCGGTGGAGCTAATGTTGATAATGGCTTTGTGCCTGACACGCTACTCTCGAGGGGATATGACTCCCAGAAAGATC<769
1158<GACGGTGAT-CCAGGAACAGCTTGCTCGGTGGAGCTAATGTTGATAATGGCTTTGTGCCTGACACGCTACTCTCGAGGGGATATGACTCCCAGAAAGATC<1060
1014<[redacted]<915
 868<GACGGTGATTCCAGGAACAGCTTGCTCGGTGGAGCTAATGTTGATAATGGCTTTGTGCCTGACACGCTACTCTCGAGGGGATATGACTCCCAGAAAGATC<769
1014<[redacted]<915

         *         *         *         *         *         *         *         *         *         *
5796>ttcagaacatgctttcaaactatggaggagtgacaaatgacattggtacagagatgtctacttcagctgtaagaactcaatcttttggtgtccccaatgt>5895
 913<TTCAGAACATGCTTTCAAACTATGGAGGAGTGACAAATGACATTGGTACAGAGATGTCTACTTCAGCTGTAAGAACTCAATCTTTTGGTGTCCCCAATGT<814
 768<TTCAGAACATGCTTTCAAACTATGGAGGAGTGACAAATGACATTGGTACAGAGATGTCTACTTCAGCTGTAAGAACTCAATCTTTTGGTGTCCCCAATGT<669
1059<TTCAGAACATGCTTTCAAACTATGGAGGAGTGACAAATGACATTGGTACAGAGATGTCTACTTCAGCTGTAAGAACTCAATCTTTTGGTGTCCCCAATGT<960
 914<[redacted]<815
 768<TTCAGAACATGCTTTCAAACTATGGAGGAGTGACAAATGACATTGGTACAGAGATGTCTACTTCAGCTGTAAGAACTCAATCTTTTGGTGTCCCCAATGT<669
 914<[redacted]<815

         *         *         *         *         *         *         *         *         *         *
5896>gcccgccatttcgaacgatctagctgtcaacgatgctggagttcttggtggtggattgtggccagctcagactcagcgaatgcgaactggcagcgatctg>5995
 813<GCCCGCCATTTCGAACGATCTAGCTGTCAACGATGCTGGAGTTCTTGGTGGTGGATTGTGGCCAGCTCAGACTCAGCGAATGCGAACTGGCAGCGATCTG<714
 668<GCCCGCCATTTCGAACGATCTAGCTGTCAACGATGCTGGAGTTCTTGGTGGTGGATTGTGGCCAGCTCAGACTCAGCGAATGCGAACTGGCAGCGATCTG<569
```

```
 959<GCCCGCCATTTCGAACGATCTAGCTGTCAACGATGCTGGAGTTCTTGGTGGTGGATTGTGGCCAGCTCAGACTCAGCGAATGCGAACTGGCAGCGATCTG<860
 814<[redacted]GGCAGCGATCTG<715
 668<GCCCGCCATTTCGAACGATCTAGCTGTCAACGATGCTGGAGTTCTTGGTGGTGGATTGTGGCCAGCTCAGACTCAGCGAATGCGAACTGGCAGCGATCTG<569
 814<[redacted]GGCAGCGATCTG<715

         *         *         *         *         *         *         *         *         *         *
5996>ggtaaaaagctgctggaagcagccgcggccggccaagatgatgaggtgcgtattctgatggcgaatggggccgatgttaacgcaaccgacgacgatggcc>6095
 713<GGTAAAAAGCTGCTGGAAGCAGCCGCGGCCGGCCAAGATGATGAGGTGCGTATTCTGATGGCGAATGGGGCCGATGTTAACGCAACCGACGACGATGGCC<614
 568<GGTAAAAAGCTGCTGGAAGCAGCCGCGGCCGGCCAAGATGATGAGGTGCGTATTCTGATGGCGAATGGGGCCGATGTTAACGCAACCGACGACGATGGCC<469
 859<GGTAAAAAGCTGCTGGAAGCAGCCGCGGCCGGCCAAGATGATGAGGTGCGTATTCTGATGGCGAATGGGGCCGATGTTAACGCAACCGACGACGATGGCC<760
 714<GGTAAAAAGCTGCTGGAAGCAGCCGCGGCCGGCCAAGATGATGAGGTGCGTATTCTGATGGCGAATGGGGCCGATGTTAACGCAACCGACGACAATGGCC<615
 568<GGTAAAAAGCTGCTGGAAGCAGCCGCGGCCGGCCAAGATGATGAGGTGCGTATTCTGATGGCGAATGGGGCCGATGTTAACGCAACCGACGACGATGGCC<469
 714<GGTAAAAAGCTGCTGGAAGCAGCCGCGGCCGGCCAAGATGATGAGGTGCGTATTCTGATGGCGAATGGGGCCGATGTTAACGCAACCGACGACAATGGCC<615

         *         *         *         *         *         *         *
6096>tgactccgctgcacctggcggctgcaaacgggcaactggaaatcgtagaggtactgctgaaaaatggcgc------------------------------>6165
 613<TGACTCCGCTGCACCTGGCGGCTGCAAACGGGCAACTGGAAATCGTAGAGGTACTGCTGAAAAATGGCGC------------------------------<544
 468<TGACTCCGCTGCACCTGGCGGCTGCAAACGGGCAACTGGAAATCGTAGAGGTACTGCTGAAAAATGGCGC------------------------------<399
 759<TGACTCCGCTGCACCTGGCGGCTGCAAACGGGCAACTGGAAATCGTAGAGGTACTGCTGAAAAATGGCGCCGATGTTAACGCAACCGACGACGATGGCCT<660
 614<TGACTCCGCTGCACCTGGCGGCTGCAAACGGGCAACTGGAAATCGTAGAGGTACTGCTGAAAAATGGCGC------------------------------<545
 468<TGACTCCGCTGCACCTGGCGGCTGCAAACGGGCAACTGGAAATCGTAGAGGTACTGCTGAAAAATGGCGC------------------------------<399
 614<TGACTCCGCTGCACCTGGCGGCTGCAAACGGGCAACTGGAAATCGTAGAGGTACTGCTGAAAAATGGCGC------------------------------<545

6165>---------------------------------------------------------------------------------------------------->6165
 544<----------------------------------------------------------------------------------------------------<544
 399<----------------------------------------------------------------------------------------------------<399
 659<GACTCCGCTGCACCTGCCGATGTTAACGCAACCGACGACGATGGCCTGACTCCGCTGCACCTGGCGGCTGCAAACGGGCAACTGGAAATCGTAGAGGTAC<560
 545<----------------------------------------------------------------------------------------------------<545
 399<----------------------------------------------------------------------------------------------------<399
 545<----------------------------------------------------------------------------------------------------<545

                         *         *         *         *         *         *         *         *
6166>----------------cgatgtgaacgcttctgatagtgcgggtattactccgctgcacctggccgcttatgacggccatctggagattgtcgaagtcct>6249
 543<----------------CGATGTGAACGCTTCTGATAGTGCGGGTATTACTCCGCTGCACCTGGCCGCTTATGACGGCCATCTGGAGATTGTCGAAGTCCT<460
 398<----------------CGATGTGAACGCTTCTGATAGTGCGGGTATTACTCCGCTGCACCTGGCCGCTTATGACGGCCATCTGGAGATTGTCGAAGTCCT<315
 559<TGCTGAAAAATGGCGCCGATGTGAACGCTTCTGATAGTGCGGGTATTACTCCGCTGCACCTGGCCGCTTATGACGGCCATCTGGAGATTGTCGAAGTCCT<460
 544<----------------CGATGTGAACGCTTCTGATAGTGCGGGTATTACTCCGCTGCACCTGGCCGCTTATGACGGCCATCTGGAGATTGTCGAAGTCCT<461
 398<----------------CGATGTGAACGCTTCTGATAGTGCGGGTATTACTCCGCTGCACCTGGCCGCTTATGACGGCCATCTGGAGATTGTCGAAGTCCT<315
```

```
 544<----------------CGATGTGAACGCTTCTGATAGTGCGGGTATTACTCCGCTGCACCTGGCCGCTTATGACGGCCATCTGGAGATTGTCGAAGTCCT<461

     *         *         *         *         *         *         *         *         *         *
6250>gctgaagcacggggctgacgttaatgcgtacgaccgcgccgggtggacaccgctgcacctagcagcgctgagtggccaactggagattgtggaagttctg>6349
 459<GCTGAAGCACGGGGCTGACGTTAATGCGTACGACCGCGCCGGGTGGACACCGCTGCACCTAGCAGCGCTGAGTGGCCAACTGGAGATTGTGGAAGTTCTG<360
 314<GCTGAAGCACGGGGCTGACGTTAATGCGTACGACCGCGCCGGGTGGACACCGCTGCACCTAGCAGCGCTGAGTGGCCAACTGGAGATTGTGGAAGTTCTG<215
 459<GCTGAAGCACGGGGCTGACGTTAATGCGTACGACCGCGCCGGGTGGACACCGCTGCACCTAGCAGCGCTGAGTGGCCAACTGGAGATTGTGGAAGTTCTG<360
 460<GCTGAAGCACGGGGCTGACGTTAATGCGTACGACCGCGCCGGGTGGACACCGCTGCACCTAGCAGCGCTGAGTGGCCAACTGGAGATTGTGGAAGTTCTG<361
 314<GCTGAAGCACGGGGCTGACGTTAATGCGTACGACCGCGCCGGGTGGACACCGCTGCACCTAGCAGCGCTGAGTGGCCAACTGGAGATTGTGGAAGTTCTG<215
 460<GCTGAAGCACGGGGCTGACGTTAATGCGTACGACCGCGCCGGGTGGACACCGCTGCACCTAGCAGCGCTGAGTGGCCAACTGGAGATTGTGGAAGTTCTG<361

     *         *         *         *         *         *         *         *         *         *
6350>ctgaaacacggcgcagatgtcaacgcccaagacgcactgggcctgaccgcgtttgatatctcgattaatcaaggtcaggaagatctggcagagatcctgc>6449
 359<CTGAAACACGGCGCAGATGTCAACGCCCAAGACGCACTGGGCCTGACCGCGTTTGATATCTCGATTAATCAAGGTCAGGAAGATCTGGCAGAGATCCTGC<260
 214<CTGAAACACGGCGCAGATGTCAACGCCCAAGACGCACTGGGCCTGACCGCGTTTGATATCTCGATTAATCAAGGTCAGGAAGATCTGGCAGAGATCCTGC<115
 359<CTGAAACACGGCGCAGATGTCAACGCCCAAGACGCACTGGGCCTGACCGCGTTTGATATCTCGATTAATCAAGGTCAGGAAGATCTGGCAGAGATCCTGC<260
 360<CTGAAACACGGCGCAGATGTCAACGCCCAAGACGCACTGGGCCTGACCGCGTTTGATATCTCGATTAATCAAGGTCAGGAAGATCTGGCAGAGATCCTGC<261
 214<CTGAAACACGGCGCAGATGTCAACGCCCAAGACGCACTGGGCCTGACCGCGTTTGATATCTCGATTAATCAAGGTCAGGAAGATCTGGCAGAGATCCTGC<115
 360<CTGAAACACGGCGCAGATGTCAACGCCCAAGACGCACTGGGCCTGACCGCGTTTGATATCTCGATTAATCAAGGTCAGGAAGATCTGGCAGAGATCCTGC<261

     *         *         *         *
              H  H  H  H  H  H      ---------------------------------------------------------------------
             CACcacCACcacCACcac     ---------------------------------------------------------------------
6450>aactcgagcaccaccaccaccaccactgaca--------------------------------------------------------------------->6480
 259<AACTCGAGCACCACCACCACCACCACTGACACCCAGCTTTCTTGTCCACCACCACTGACACCCAGCTTTCTTGTCCACCACCACTGACACCCAGCTTTCT<160
 114<AACTCGAGCACCACCACCACCACCACTGACA---------------------------------------------------------------------<84
 259<AACTCGAGCACCACCACCACCACCACTGACACCCAGCTTTCTTGTCCACCACCACTGACACCCAGCTTTCTTGTCCACCACCACTGACACCCAGCTTTCT<160
 260<AACTCGAGCACCACCACCACCACCACTGACACCCAGCTTTCTTGTCCACCACCACTGACACCCAGCTTTCTTGTCCACCACCACTGACACCCAGCTTTCT<161
 114<AACTCGAGCACCACCACCACCACCACTGACA---------------------------------------------------------------------<84
 260<AACTCGAGCACCACCACCACCACCACTGACACCCAGCTTTCTTGTCCACCACCACTGACACCCAGCTTTCTTGTCCACCACCACTGACACCCAGCTTTCT<161

                                                                                          *         *
6481>----------------------------------------------------------------------------cccagctttcttgtacaaagtggt>6504
 159<TGTCCACCACCACTGACACCCAGCTTTCTTGTCCACCACCACTGACACCCAGCTTTCTTGTCCACCACCACTGACACCCAGCTTTCTTGT----------<70
  83<----------------------------------------------------------------------------CCCAGCTTTCTTGTAC--------<68
 159<TGTCCACCACCACTGACACCCAGCTTTCTTGTCCACCACCACTGACACCCAGCTTTCTTGTCCACCACCACTGACACCCAGCTTTCTTGT----------<70
 160<TGTCCACCACCACTGACACCCAGCTTTCTTGTCCACCACCACTGACACCCAGCTTTCTTGTCCACCACCACTGACACCCAGCTTTCTTGT----------<71
  83<----------------------------------------------------------------------------CCCAGCTTTCTTGTAC--------<68
 160<TGTCCACCACCACTGACACCCAGCTTTCTTGTCCACCACCACTGACACCCAGCTTTCTTGTCCACCACCACTGACACCCAGCTTTCTTGT----------<71
```

```
         *         *         *         *         *         *         *         *         *         *
6505>tcttgtacaaagtggtgatgggctgcaggaattcgatatcaagcttatcgataccgtcgacctcgagtcatgtaattagttatgtcacgcttacattcac>6604
  69<------ACAAAGTGGTGATGGGCTGCAGGAATTCGAT-TCAAGCTTATCGATACCGTCGACCTCGAGTCA------------------------------<7
  67<--------AAAGTGGTGATGGGCTGCAGGAATTCGATTTCAAGCTTATCGATACCGTCGACCTCGAGTCA------------------------------<6
  69<------ACAAAGTGGTGATGGGCTGCAGGAATTCGATTTCAAGCTTATCGATACCGTCGACCTCGAGTCA------------------------------<6
  70<------ACAAAGTGGTGATGGGCTGCAGGAATTCGATTTCAAGCTTATCGATACCGTCGACCTCGAGTC-------------------------------<8
  67<--------AAAGTGGTGATGGGCTGCAGGAATTCGAT-TCAAGCTTATCGATACCGTCGACCTCGAGTCA------------------------------<7
  70<------ACAAAGTGGTGATGGGCTGCAGGAATTCGATTTCAAGCTTATCGATACCGTCGACCTCGAGTCA------------------------------<7

         *         *         *         *         *         *         *         *         *         *
6605>gccctccccccacatccgctctaaccgaaaaggaaggagttagacaacctgaagtctaggtccctatttattttttatagttatgttagtattaagaac>6704
   7<----------------------------------------------------------------------------------------------------<7
   6<----------------------------------------------------------------------------------------------------<6
   6<----------------------------------------------------------------------------------------------------<6
   8<----------------------------------------------------------------------------------------------------<8
   7<----------------------------------------------------------------------------------------------------<7
   7<----------------------------------------------------------------------------------------------------<7

         *         *         *         *         *         *         *         *         *         *
6705>gttatttatatttcaaattttctttttttttctgtacagacgcgtgtacgcatgtaacattatactgaaaaccttgcttgagaaggttttgggacgctcg>6804
   7<----------------------------------------------------------------------------------------------------<7
   6<----------------------------------------------------------------------------------------------------<6
   6<----------------------------------------------------------------------------------------------------<6
   8<----------------------------------------------------------------------------------------------------<8
   7<----------------------------------------------------------------------------------------------------<7
   7<----------------------------------------------------------------------------------------------------<7

         *         *         *         *         *         *         *         *         *         *
6805>aaggctttaatttgtgacaccgattatttaaagctgcagcatacgatatatatacatgtgtatatatgtatacctatgaatgtcagtaagtatgtatacg>6904
   7<----------------------------------------------------------------------------------------------------<7
   6<----------------------------------------------------------------------------------------------------<6
   6<----------------------------------------------------------------------------------------------------<6
   8<----------------------------------------------------------------------------------------------------<8
   7<----------------------------------------------------------------------------------------------------<7
   7<----------------------------------------------------------------------------------------------------<7

         *         *         *         *         *         *         *         *         *         *
```

```
6905>aacagtatgatactgaagatgacaaggtaatgcatcattctatacgtgtcattctgaacgaggcgcgctttccttttttctttttgctttttcttttttt>7004
   7<----------------------------------------------------------------------------------------------------<7
   5<----GTATG~~~~~~~~~~~~~~~~~~~~~~~~~~~~~~~~~~~~~~~~~~~~~~~~~~~~~~~~~~~~~~~~~~~~~~~~~~~~~~~~~~~~~~~~~~<1
   6<----------------------------------------------------------------------------------------------------<6
   8<----------------------------------------------------------------------------------------------------<8
   7<----------------------------------------------------------------------------------------------------<7
   7<----------------------------------------------------------------------------------------------------<7

          *         *         *         *         *         *         *         *         *         *
7005>ttctcttgaactcgagaaaaaaaatataaaagagatggaggaacgggaaaaagttagttgtggtgataggtggcaagtggtattccgtaagaacaacaag>7104
   7<----------------------------------------------------------------------------------------------------<7
   1<~~~~~~~~~~~~~~~~~~~~~~~~~~~~~~~~~~~~~~~~~~~~~~~~~~~~~~~~~~~~~~~~~~~~~~~~~~~~~~~~~~~~~~~~~~~~~~~~~~~~<1
   5<--------------------------------------------------------------------------------------GTAAG~~~~~~~~~<1
   8<----------------------------------------------------------------------------------------------------<8
   7<----------------------------------------------------------------------------------------------------<7
   6<------------------------------------------------------------------------------GGTATT~~~~~~~~~~~~~~~~<1

          *         *         *         *         *         *         *         *         *         *
7105>aaaagcatttcatattatggctgaactgagcgaacaagtgcaaaatttaagcatcaacgacaacaacgagaatggttatgttcctcctcacttaagagga>7204
   7<----------------------------------------------------------------------------------------------------<7
   1<~~~~~~~~~~~~~~~~~~~~~~~~~~~~~~~~~~~~~~~~~~~~~~~~~~~~~~~~~~~~~~~~~~~~~~~~~~~~~~~~~~~~~~~~~~~~~~~~~~~<1
   1<~~~~~~~~~~~~~~~~~~~~~~~~~~~~~~~~~~~~~~~~~~~~~~~~~~~~~~~~~~~~~~~~~~~~~~~~~~~~~~~~~~~~~~~~~~~~~~~~~~~<1
   8<----------------------------------------------------------------------------------------------------<8
   7<----------------------------------------------------------------------------------------------------<7
   1<~~~~~~~~~~~~~~~~~~~~~~~~~~~~~~~~~~~~~~~~~~~~~~~~~~~~~~~~~~~~~~~~~~~~~~~~~~~~~~~~~~~~~~~~~~~~~~~~~~~<1

          *         *         *         *         *         *         *         *         *         *
7205>aaaccaagaagtgccagaaataacagtagcaactacaataacaacaacggcggctacaacggtggccgtggcggtggcagcttctttagcaacaaccgtc>7304
   7<----------------------------------------------------------------------------------------------------<7
   1<~~~~~~~~~~~~~~~~~~~~~~~~~~~~~~~~~~~~~~~~~~~~~~~~~~~~~~~~~~~~~~~~~~~~~~~~~~~~~~~~~~~~~~~~~~~~~~~~~~~<1
   1<~~~~~~~~~~~~~~~~~~~~~~~~~~~~~~~~~~~~~~~~~~~~~~~~~~~~~~~~~~~~~~~~~~~~~~~~~~~~~~~~~~~~~~~~~~~~~~~~~~~<1
   8<----------------------------------------------------------------------------------------------------<8
   7<----------------------------------------------------------------------------------------------------<7
   1<~~~~~~~~~~~~~~~~~~~~~~~~~~~~~~~~~~~~~~~~~~~~~~~~~~~~~~~~~~~~~~~~~~~~~~~~~~~~~~~~~~~~~~~~~~~~~~~~~~~<1

          *         *         *         *         *         *         *         *         *         *
7305>gtggtggttacggcaacggtggtttcttcggtggaaacaacggtggcagcagatctaacggccgttctggtggtagatggatcgatggcaaacatgtccc>7404
   7<----------------------------------------------------------------------------------------------------<7
   1<~~~~~~~~~~~~~~~~~~~~~~~~~~~~~~~~~~~~~~~~~~~~~~~~~~~~~~~~~~~~~~~~~~~~~~~~~~~~~~~~~~~~~~~~~~~~~~~~~~~<1
```

```
   1<~~~~~~~~~~~~~~~~~~~~~~~~~~~~~~~~~~~~~~~~~~~~~~~~~~~~~~~~~~~~~~~~~~~~~~~~~~~~~~~~~~~~~~~~~~~~~~~~<1
   8<----------------------------------------------------------------------------------------------------<8
   7<----------------------------------------------------------------------------------------------------<7
   1<~~~~~~~~~~~~~~~~~~~~~~~~~~~~~~~~~~~~~~~~~~~~~~~~~~~~~~~~~~~~~~~~~~~~~~~~~~~~~~~~~~~~~~~~~~~~~~~~<1

              *         *         *         *         *         *         *         *         *         *
7405>agctccaagaaacgaaaaggccgagatcgccatatttggtgtccccgaggatccaaatttccaatcttctggtattaacttcgataactacgatgatatt>7504
   7<----------------------------------------------------------------------------------------------------<7
   1<~~~~~~~~~~~~~~~~~~~~~~~~~~~~~~~~~~~~~~~~~~~~~~~~~~~~~~~~~~~~~~~~~~~~~~~~~~~~~~~~~~~~~~~~~~~~~~~~<1
   1<~~~~~~~~~~~~~~~~~~~~~~~~~~~~~~~~~~~~~~~~~~~~~~~~~~~~~~~~~~~~~~~~~~~~~~~~~~~~~~~~~~~~~~~~~~~~~~~~<1
   8<----------------------------------------------------------------------------------------------------<8
   7<----------------------------------------------------------------------------------------------------<7
   1<~~~~~~~~~~~~~~~~~~~~~~~~~~~~~~~~~~~~~~~~~~~~~~~~~~~~~~~~~~~~~~~~~~~~~~~~~~~~~~~~~~~~~~~~~~~~~~~~<1

              *         *         *         *         *         *         *         *         *         *
7505>ccagtggacgcctctggtaaggatgttcctgaaccaatcacagaatttacctcacctccattggacggattgttattggaaaacatcaaattggcccgtt>7604
   7<----------------------------------------------------------------------------------------------------<7
   1<~~~~~~~~~~~~~~~~~~~~~~~~~~~~~~~~~~~~~~~~~~~~~~~~~~~~~~~~~~~~~~~~~~~~~~~~~~~~~~~~~~~~~~~~~~~~~~~~<1
   1<~~~~~~~~~~~~~~~~~~~~~~~~~~~~~~~~~~~~~~~~~~~~~~~~~~~~~~~~~~~~~~~~~~~~~~~~~~~~~~~~~~~~~~~~~~~~~~~~<1
   8<----------------------------------------------------------------------------------------------------<8
   7<----------------------------------------------------------------------------------------------------<7
   1<~~~~~~~~~~~~~~~~~~~~~~~~~~~~~~~~~~~~~~~~~~~~~~~~~~~~~~~~~~~~~~~~~~~~~~~~~~~~~~~~~~~~~~~~~~~~~~~~<1

              *         *         *         *         *         *         *         *         *         *
7605>tcaccaagccaacacctgtgcaaaaatactccgtccctatcgttgccaacggcagagatttgatggcctgtgcgcagaccggttctggtaagactggtgg>7704
   7<----------------------------------------------------------------------------------------------------<7
   1<~~~~~~~~~~~~~~~~~~~~~~~~~~~~~~~~~~~~~~~~~~~~~~~~~~~~~~~~~~~~~~~~~~~~~~~~~~~~~~~~~~~~~~~~~~~~~~~~<1
   1<~~~~~~~~~~~~~~~~~~~~~~~~~~~~~~~~~~~~~~~~~~~~~~~~~~~~~~~~~~~~~~~~~~~~~~~~~~~~~~~~~~~~~~~~~~~~~~~~<1
   8<----------------------------------------------------------------------------------------------------<8
   7<----------------------------------------------------------------------------------------------------<7
   1<~~~~~~~~~~~~~~~~~~~~~~~~~~~~~~~~~~~~~~~~~~~~~~~~~~~~~~~~~~~~~~~~~~~~~~~~~~~~~~~~~~~~~~~~~~~~~~~~<1

              *         *         *         *         *         *         *         *         *         *
7705>gtttttattcccagtgttgtccgaatcatttaagactggaccatctcctcaaccagagtctcaaggctccttttaccaaagaaaggcctacccaactgct>7804
   7<----------------------------------------------------------------------------------------------------<7
   1<~~~~~~~~~~~~~~~~~~~~~~~~~~~~~~~~~~~~~~~~~~~~~~~~~~~~~~~~~~~~~~~~~~~~~~~~~~~~~~~~~~~~~~~~~~~~~~~~<1
   1<~~~~~~~~~~~~~~~~~~~~~~~~~~~~~~~~~~~~~~~~~~~~~~~~~~~~~~~~~~~~~~~~~~~~~~~~~~~~~~~~~~~~~~~~~~~~~~~~<1
   8<----------------------------------------------------------------------------------------------------<8
   7<----------------------------------------------------------------------------------------------------<7
```

```
   1<~~~~~~~~~~~~~~~~~~~~~~~~~~~~~~~~~~~~~~~~~~~~~~~~~~~~~~~~~~~~~~~~~~~~~~~~~~~~~~~~~~~~~~~~~~~~~~~~~<1

          *         *         *         *         *         *         *         *         *         *
7805>gtcattatggctccagtttaaaccatggtcatagctgtttcctgtgtgaaattgttatccgctcacaattccacacaacataggagccggaagcataaag>7904
   7<----------------------------------------------------------------------------------------------------<7
   1<~~~~~~~~~~~~~~~~~~~~~~~~~~~~~~~~~~~~~~~~~~~~~~~~~~~~~~~~~~~~~~~~~~~~~~~~~~~~~~~~~~~~~~~~~~~~~~~~~<1
   1<~~~~~~~~~~~~~~~~~~~~~~~~~~~~~~~~~~~~~~~~~~~~~~~~~~~~~~~~~~~~~~~~~~~~~~~~~~~~~~~~~~~~~~~~~~~~~~~~~<1
   8<----------------------------------------------------------------------------------------------------<8
   7<----------------------------------------------------------------------------------------------------<7
   1<~~~~~~~~~~~~~~~~~~~~~~~~~~~~~~~~~~~~~~~~~~~~~~~~~~~~~~~~~~~~~~~~~~~~~~~~~~~~~~~~~~~~~~~~~~~~~~~~~<1

          *         *         *         *         *         *         *         *         *         *
7905>tgtaaagcctggggtgcctaatgagtgaggtaactcacattaattgcgttgcgctcactgcccgctttccagtcgggaaacctgtcgtgccagctgcatt>8004
   6<-GTAAAG~~~~~~~~~~~~~~~~~~~~~~~~~~~~~~~~~~~~~~~~~~~~~~~~~~~~~~~~~~~~~~~~~~~~~~~~~~~~~~~~~~~~~~<1
   1<~~~~~~~~~~~~~~~~~~~~~~~~~~~~~~~~~~~~~~~~~~~~~~~~~~~~~~~~~~~~~~~~~~~~~~~~~~~~~~~~~~~~~~~~~~~~~~~~~<1
   1<~~~~~~~~~~~~~~~~~~~~~~~~~~~~~~~~~~~~~~~~~~~~~~~~~~~~~~~~~~~~~~~~~~~~~~~~~~~~~~~~~~~~~~~~~~~~~~~~~<1
   8<----------------------------------------------------------------------------------------------------<8
   7<----------------------------------------------------------------------------------------------------<7
   1<~~~~~~~~~~~~~~~~~~~~~~~~~~~~~~~~~~~~~~~~~~~~~~~~~~~~~~~~~~~~~~~~~~~~~~~~~~~~~~~~~~~~~~~~~~~~~~~~~<1

          *         *         *         *         *         *         *         *         *         *
8005>aatgaatcggccaacgcgcggggagaggcggtttgcgtattgggcgctcttccgcttcctcgctcactgactcgctgcgctcggtcgttcggctgcggcg>8104
   1<~~~~~~~~~~~~~~~~~~~~~~~~~~~~~~~~~~~~~~~~~~~~~~~~~~~~~~~~~~~~~~~~~~~~~~~~~~~~~~~~~~~~~~~~~~~~~~~~~<1
   1<~~~~~~~~~~~~~~~~~~~~~~~~~~~~~~~~~~~~~~~~~~~~~~~~~~~~~~~~~~~~~~~~~~~~~~~~~~~~~~~~~~~~~~~~~~~~~~~~~<1
   1<~~~~~~~~~~~~~~~~~~~~~~~~~~~~~~~~~~~~~~~~~~~~~~~~~~~~~~~~~~~~~~~~~~~~~~~~~~~~~~~~~~~~~~~~~~~~~~~~~<1
   8<----------------------------------------------------------------------------------------------------<8
   7<----------------------------------------------------------------------------------------------------<7
   1<~~~~~~~~~~~~~~~~~~~~~~~~~~~~~~~~~~~~~~~~~~~~~~~~~~~~~~~~~~~~~~~~~~~~~~~~~~~~~~~~~~~~~~~~~~~~~~~~~<1

          *         *         *         *         *         *         *         *         *         *
8105>agcggtatcagctcactcaaaggcggtaatacggttatccacagaatcaggggataacgcaggaaagaacatgtgagcaaaaggccagcaaaaggccagg>8204
   1<~~~~~~~~~~~~~~~~~~~~~~~~~~~~~~~~~~~~~~~~~~~~~~~~~~~~~~~~~~~~~~~~~~~~~~~~~~~~~~~~~~~~~~~~~~~~~~~~~<1
   1<~~~~~~~~~~~~~~~~~~~~~~~~~~~~~~~~~~~~~~~~~~~~~~~~~~~~~~~~~~~~~~~~~~~~~~~~~~~~~~~~~~~~~~~~~~~~~~~~~<1
   1<~~~~~~~~~~~~~~~~~~~~~~~~~~~~~~~~~~~~~~~~~~~~~~~~~~~~~~~~~~~~~~~~~~~~~~~~~~~~~~~~~~~~~~~~~~~~~~~~~<1
   8<----------------------------------------------------------------------------------------------------<8
   7<----------------------------------------------------------------------------------------------------<7
   1<~~~~~~~~~~~~~~~~~~~~~~~~~~~~~~~~~~~~~~~~~~~~~~~~~~~~~~~~~~~~~~~~~~~~~~~~~~~~~~~~~~~~~~~~~~~~~~~~~<1
```

```
          *         *         *         *         *         *         *         *         *         *
8205>aaccgtaaaaaggccgcgttgctggcgtttttccataggctcggcccccctgacgagcatcacaaaaatcgacgctcaagtcagaggtggcgaaacccga>8304
   1<~~~~~~~~~~~~~~~~~~~~~~~~~~~~~~~~~~~~~~~~~~~~~~~~~~~~~~~~~~~~~~~~~~~~~~~~~~~~~~~~~~~~~~~~~~~~~~~~<1
   1<~~~~~~~~~~~~~~~~~~~~~~~~~~~~~~~~~~~~~~~~~~~~~~~~~~~~~~~~~~~~~~~~~~~~~~~~~~~~~~~~~~~~~~~~~~~~~~~~<1
   1<~~~~~~~~~~~~~~~~~~~~~~~~~~~~~~~~~~~~~~~~~~~~~~~~~~~~~~~~~~~~~~~~~~~~~~~~~~~~~~~~~~~~~~~~~~~~~~~~<1
   8<----------------------------------------------------------------------------------------------------<8
   7<----------------------------------------------------------------------------------------------------<7
   1<~~~~~~~~~~~~~~~~~~~~~~~~~~~~~~~~~~~~~~~~~~~~~~~~~~~~~~~~~~~~~~~~~~~~~~~~~~~~~~~~~~~~~~~~~~~~~~~~~<1

          *         *         *         *         *         *         *         *         *         *
8305>caggactataaagataccaggcgttcccccctggaagctccctcgtgcgctctcctgttccgaccctgccgcttaccggatacctgtccgcctttctccc>8404
   1<~~~~~~~~~~~~~~~~~~~~~~~~~~~~~~~~~~~~~~~~~~~~~~~~~~~~~~~~~~~~~~~~~~~~~~~~~~~~~~~~~~~~~~~~~~~~~~~~<1
   1<~~~~~~~~~~~~~~~~~~~~~~~~~~~~~~~~~~~~~~~~~~~~~~~~~~~~~~~~~~~~~~~~~~~~~~~~~~~~~~~~~~~~~~~~~~~~~~~~<1
   1<~~~~~~~~~~~~~~~~~~~~~~~~~~~~~~~~~~~~~~~~~~~~~~~~~~~~~~~~~~~~~~~~~~~~~~~~~~~~~~~~~~~~~~~~~~~~~~~~<1
   8<----------------------------------------------------------------------------------------------------<8
   7<----------------------------------------------------------------------------------------------------<7
   1<~~~~~~~~~~~~~~~~~~~~~~~~~~~~~~~~~~~~~~~~~~~~~~~~~~~~~~~~~~~~~~~~~~~~~~~~~~~~~~~~~~~~~~~~~~~~~~~~~<1

          *         *         *         *         *         *         *         *         *         *
8405>ttcgggaagcgtggcgctttctcaatgctcacgctgtaggtatctcagttcggtgtaggtcgttcgctccaagctgggctgtgtgcacgaaccccccgtt>8504
   1<~~~~~~~~~~~~~~~~~~~~~~~~~~~~~~~~~~~~~~~~~~~~~~~~~~~~~~~~~~~~~~~~~~~~~~~~~~~~~~~~~~~~~~~~~~~~~~~~<1
   1<~~~~~~~~~~~~~~~~~~~~~~~~~~~~~~~~~~~~~~~~~~~~~~~~~~~~~~~~~~~~~~~~~~~~~~~~~~~~~~~~~~~~~~~~~~~~~~~~<1
   1<~~~~~~~~~~~~~~~~~~~~~~~~~~~~~~~~~~~~~~~~~~~~~~~~~~~~~~~~~~~~~~~~~~~~~~~~~~~~~~~~~~~~~~~~~~~~~~~~<1
   8<----------------------------------------------------------------------------------------------------<8
   7<----------------------------------------------------------------------------------------------------<7
   1<~~~~~~~~~~~~~~~~~~~~~~~~~~~~~~~~~~~~~~~~~~~~~~~~~~~~~~~~~~~~~~~~~~~~~~~~~~~~~~~~~~~~~~~~~~~~~~~~~<1

          *         *         *         *         *         *         *         *         *         *
8505>cagcccgaccgctgcgccttatccggtaactatcgtcttgagtccaacccggtaagacacgacttatcgccactggcagcagccactggtaacaggatta>8604
   1<~~~~~~~~~~~~~~~~~~~~~~~~~~~~~~~~~~~~~~~~~~~~~~~~~~~~~~~~~~~~~~~~~~~~~~~~~~~~~~~~~~~~~~~~~~~~~~~~<1
   1<~~~~~~~~~~~~~~~~~~~~~~~~~~~~~~~~~~~~~~~~~~~~~~~~~~~~~~~~~~~~~~~~~~~~~~~~~~~~~~~~~~~~~~~~~~~~~~~~<1
   1<~~~~~~~~~~~~~~~~~~~~~~~~~~~~~~~~~~~~~~~~~~~~~~~~~~~~~~~~~~~~~~~~~~~~~~~~~~~~~~~~~~~~~~~~~~~~~~~~<1
   8<----------------------------------------------------------------------------------------------------<8
   7<----------------------------------------------------------------------------------------------------<7
   1<~~~~~~~~~~~~~~~~~~~~~~~~~~~~~~~~~~~~~~~~~~~~~~~~~~~~~~~~~~~~~~~~~~~~~~~~~~~~~~~~~~~~~~~~~~~~~~~~~<1

          *         *         *         *         *         *         *         *         *         *
8605>gcagagcgaggtatgtaggcggtgctacagagttcttgaagtggtggcctaactacggctacactagaaggacagtatttggtatctgcgctctgctgaa>8704
   1<~~~~~~~~~~~~~~~~~~~~~~~~~~~~~~~~~~~~~~~~~~~~~~~~~~~~~~~~~~~~~~~~~~~~~~~~~~~~~~~~~~~~~~~~~~~~~~~~<1
```

```
   1<~~~~~~~~~~~~~~~~~~~~~~~~~~~~~~~~~~~~~~~~~~~~~~~~~~~~~~~~~~~~~~~~~~~~~~~~~~~~~~~~~~~~~~~~~~~~~~~~<1
   1<~~~~~~~~~~~~~~~~~~~~~~~~~~~~~~~~~~~~~~~~~~~~~~~~~~~~~~~~~~~~~~~~~~~~~~~~~~~~~~~~~~~~~~~~~~~~~~~<1
   8<----------------------------------------------------------------------------------------------------<8
   7<----------------------------------------------------------------------------------------------------<7
   1<~~~~~~~~~~~~~~~~~~~~~~~~~~~~~~~~~~~~~~~~~~~~~~~~~~~~~~~~~~~~~~~~~~~~~~~~~~~~~~~~~~~~~~~~~~~~~~~<1


          *         *         *         *         *         *         *         *         *         *    
8705>gccagttaccttcggaaaaagagttggtagctcttgatccggcaaacaaaccaccgctggtagcggtggtttttttgtttgcaagcagcagattacgcgc>8804
   1<~~~~~~~~~~~~~~~~~~~~~~~~~~~~~~~~~~~~~~~~~~~~~~~~~~~~~~~~~~~~~~~~~~~~~~~~~~~~~~~~~~~~~~~~~~~~~~~<1
   1<~~~~~~~~~~~~~~~~~~~~~~~~~~~~~~~~~~~~~~~~~~~~~~~~~~~~~~~~~~~~~~~~~~~~~~~~~~~~~~~~~~~~~~~~~~~~~~~<1
   1<~~~~~~~~~~~~~~~~~~~~~~~~~~~~~~~~~~~~~~~~~~~~~~~~~~~~~~~~~~~~~~~~~~~~~~~~~~~~~~~~~~~~~~~~~~~~~~~<1
   8<----------------------------------------------------------------------------------------------------<8
   7<----------------------------------------------------------------------------------------------------<7
   1<~~~~~~~~~~~~~~~~~~~~~~~~~~~~~~~~~~~~~~~~~~~~~~~~~~~~~~~~~~~~~~~~~~~~~~~~~~~~~~~~~~~~~~~~~~~~~~~<1


          *         *         *         *         *         *         *         *         *         *    
8805>agaaaaaaaggatctcaagaagatcctttgatcttttctacggggtctgacgctcagtggaacgaaaactcacgttaagggattttggtcatgagattat>8904
   1<~~~~~~~~~~~~~~~~~~~~~~~~~~~~~~~~~~~~~~~~~~~~~~~~~~~~~~~~~~~~~~~~~~~~~~~~~~~~~~~~~~~~~~~~~~~~~~~<1
   1<~~~~~~~~~~~~~~~~~~~~~~~~~~~~~~~~~~~~~~~~~~~~~~~~~~~~~~~~~~~~~~~~~~~~~~~~~~~~~~~~~~~~~~~~~~~~~~~<1
   1<~~~~~~~~~~~~~~~~~~~~~~~~~~~~~~~~~~~~~~~~~~~~~~~~~~~~~~~~~~~~~~~~~~~~~~~~~~~~~~~~~~~~~~~~~~~~~~~<1
   8<----------------------------------------------------------------------------------------------------<8
   7<----------------------------------------------------------------------------------------------------<7
   1<~~~~~~~~~~~~~~~~~~~~~~~~~~~~~~~~~~~~~~~~~~~~~~~~~~~~~~~~~~~~~~~~~~~~~~~~~~~~~~~~~~~~~~~~~~~~~~~<1


          *         *         *         *         *         *         *         *         *         *    
                                                                                              *  W  H  K
                                                                                             TTAccaATGctt
8905>caaaaaggatcttcacctagatccttttaaattaaaaatgaagttttaaatcaatctaaagtatatatgagtaaacttggtctgacagttaccaatgctt>9004
   1<~~~~~~~~~~~~~~~~~~~~~~~~~~~~~~~~~~~~~~~~~~~~~~~~~~~~~~~~~~~~~~~~~~~~~~~~~~~~~~~~~~~~~~~~~~~~~~~<1
   1<~~~~~~~~~~~~~~~~~~~~~~~~~~~~~~~~~~~~~~~~~~~~~~~~~~~~~~~~~~~~~~~~~~~~~~~~~~~~~~~~~~~~~~~~~~~~~~~<1
   1<~~~~~~~~~~~~~~~~~~~~~~~~~~~~~~~~~~~~~~~~~~~~~~~~~~~~~~~~~~~~~~~~~~~~~~~~~~~~~~~~~~~~~~~~~~~~~~~<1
   8<----------------------------------------------------------------------------------------------------<8
   7<----------------------------------------------------------------------------------------------------<7
   1<~~~~~~~~~~~~~~~~~~~~~~~~~~~~~~~~~~~~~~~~~~~~~~~~~~~~~~~~~~~~~~~~~~~~~~~~~~~~~~~~~~~~~~~~~~~~~~~<1


          *         *         *         *         *         *         *         *         *         *    
      I  L  S  A  G  I  E  A  I  Q  R  N  R  E  D  M  T  A  Q  S  G  T  T  Y  I  V  V  I  R  S  P  K  G
     AATcagTGAggcACCtatCTCagcGATctgTCTatttCGttcATCcatAGTtgcCTGactGCCcgtCGTgtaGATaacTACgatACGggaGGGcttACCa
9005>aatcagtgaggcacctatctcagcgatctgtctatttcgttcatccatagttgcctgactgcccgtcgtgtagataactacgatacgggagggcttacca>9104
```

```
   1<~~~~~~~~~~~~~~~~~~~~~~~~~~~~~~~~~~~~~~~~~~~~~~~~~~~~~~~~~~~~~~~~~~~~~~~~~~~~~~~~~~~~~~~~~~~~~~~~~~~~<1
   1<~~~~~~~~~~~~~~~~~~~~~~~~~~~~~~~~~~~~~~~~~~~~~~~~~~~~~~~~~~~~~~~~~~~~~~~~~~~~~~~~~~~~~~~~~~~~~~~~~~~~<1
   1<~~~~~~~~~~~~~~~~~~~~~~~~~~~~~~~~~~~~~~~~~~~~~~~~~~~~~~~~~~~~~~~~~~~~~~~~~~~~~~~~~~~~~~~~~~~~~~~~~~~~<1
   8<----------------------------------------------------------------------------------------------------<8
   7<----------------------------------------------------------------------------------------------------<7
   1<~~~~~~~~~~~~~~~~~~~~~~~~~~~~~~~~~~~~~~~~~~~~~~~~~~~~~~~~~~~~~~~~~~~~~~~~~~~~~~~~~~~~~~~~~~~~~~~~~~~~<1

          *         *         *         *         *         *         *         *         *         *
     D  P  G  L  A  A  I  I  G  R  S  G  R  E  G  A  G  S  K  D  A  I  F  W  G  A  P  L  A  S  R  L  L  P
     tcTGGccccCAGtgcTGCaatGATaccGCGagaCCCacgCTCaccGGCtccAGAtttATCagcAATaaaCCAgccAGCcggAAGggcCGAgcgCAGaagTG
9105>tctggccccagtgctgcaatgataccgcgagacccacgctcaccggctccagatttatcagcaataaaccagccagccggaagggccgagcgcagaagtg>9204
   1<~~~~~~~~~~~~~~~~~~~~~~~~~~~~~~~~~~~~~~~~~~~~~~~~~~~~~~~~~~~~~~~~~~~~~~~~~~~~~~~~~~~~~~~~~~~~~~~~~~~~<1
   1<~~~~~~~~~~~~~~~~~~~~~~~~~~~~~~~~~~~~~~~~~~~~~~~~~~~~~~~~~~~~~~~~~~~~~~~~~~~~~~~~~~~~~~~~~~~~~~~~~~~~<1
   1<~~~~~~~~~~~~~~~~~~~~~~~~~~~~~~~~~~~~~~~~~~~~~~~~~~~~~~~~~~~~~~~~~~~~~~~~~~~~~~~~~~~~~~~~~~~~~~~~~~~~<1
   8<----------------------------------------------------------------------------------------------------<8
   7<----------------------------------------------------------------------------------------------------<7
   1<~~~~~~~~~~~~~~~~~~~~~~~~~~~~~~~~~~~~~~~~~~~~~~~~~~~~~~~~~~~~~~~~~~~~~~~~~~~~~~~~~~~~~~~~~~~~~~~~~~~~<1

          *         *         *         *         *         *         *         *         *         *
       G  A  V  K  D  A  E  M  W  D  I  L  Q  Q  R  S  A  L  T  L  L  E  G  T  L  L  K  R  L  T  T  A  M
     GtccTGCaacTTTatcCGCctcCATccaGTCtatTAAttgTTGccgGGAagcTAGagtAAGtagTTCgccAGTtaaTAGtttGCGcaaCGTtgtTGCcat
9205>gtcctgcaactttatccgcctccatccagtctattaattgttgccgggaagctagagtaagtagttcgccagttaatagtttgcgcaacgttgttgccat>9304
   1<~~~~~~~~~~~~~~~~~~~~~~~~~~~~~~~~~~~~~~~~~~~~~~~~~~~~~~~~~~~~~~~~~~~~~~~~~~~~~~~~~~~~~~~~~~~~~~~~~~~~<1
   1<~~~~~~~~~~~~~~~~~~~~~~~~~~~~~~~~~~~~~~~~~~~~~~~~~~~~~~~~~~~~~~~~~~~~~~~~~~~~~~~~~~~~~~~~~~~~~~~~~~~~<1
   1<~~~~~~~~~~~~~~~~~~~~~~~~~~~~~~~~~~~~~~~~~~~~~~~~~~~~~~~~~~~~~~~~~~~~~~~~~~~~~~~~~~~~~~~~~~~~~~~~~~~~<1
   8<----------------------------------------------------------------------------------------------------<8
   7<----------------------------------------------------------------------------------------------------<7
   1<~~~~~~~~~~~~~~~~~~~~~~~~~~~~~~~~~~~~~~~~~~~~~~~~~~~~~~~~~~~~~~~~~~~~~~~~~~~~~~~~~~~~~~~~~~~~~~~~~~~~<1

          *         *         *         *         *         *         *         *         *         *
      A  V  P  M  T  T  D  R  E  D  N  P  I  A  E  N  L  E  P  E  W  R  D  L  R  T  V  H  D  G  M  N  H
     TGCtacAGGcatCGTggtGTCacgCTCgtcGTTtggTATggcTTCattCAGctcCGGttcCCAacgATCaagGCGagtTACatgATCcccCATgttGTGa
9305>tgctacaggcatcgtggtgtcacgctcgtcgtttggtatggcttcattcagctccggttcccaacgatcaaggcgagttacatgatcccccatgttgtga>9404
   1<~~~~~~~~~~~~~~~~~~~~~~~~~~~~~~~~~~~~~~~~~~~~~~~~~~~~~~~~~~~~~~~~~~~~~~~~~~~~~~~~~~~~~~~~~~~~~~~~~~~~<1
   1<~~~~~~~~~~~~~~~~~~~~~~~~~~~~~~~~~~~~~~~~~~~~~~~~~~~~~~~~~~~~~~~~~~~~~~~~~~~~~~~~~~~~~~~~~~~~~~~~~~~~<1
   1<~~~~~~~~~~~~~~~~~~~~~~~~~~~~~~~~~~~~~~~~~~~~~~~~~~~~~~~~~~~~~~~~~~~~~~~~~~~~~~~~~~~~~~~~~~~~~~~~~~~~<1
   8<----------------------------------------------------------------------------------------------------<8
   7<----------------------------------------------------------------------------------------------------<7
   1<~~~~~~~~~~~~~~~~~~~~~~~~~~~~~~~~~~~~~~~~~~~~~~~~~~~~~~~~~~~~~~~~~~~~~~~~~~~~~~~~~~~~~~~~~~~~~~~~~~~~<1
```

```
          *         *         *         *         *         *         *         *         *         *
     F  F  A  T  L  E  K  P  G  G  I  T  T  L  L  N  A  A  T  N  D  S  M  T  I  A  A  S  C  L  E  R  V
     aaAAAagcGGTtagCTCcttCGGtccTCCgatCGTtgtCAGaagTAAgttGGCcgcAGTgttATCactCATggtTATggcAGCactGCAtaaTTCtctTA
9405>aaaaaagcggttagctccttcggtcctccgatcgttgtcagaagtaagttggccgcagtgttatcactcatggttatggcagcactgcataattctctta>9504
   1<~~~~~~~~~~~~~~~~~~~~~~~~~~~~~~~~~~~~~~~~~~~~~~~~~~~~~~~~~~~~~~~~~~~~~~~~~~~~~~~~~~~~~~~~~~~~~~~~~~<1
   1<~~~~~~~~~~~~~~~~~~~~~~~~~~~~~~~~~~~~~~~~~~~~~~~~~~~~~~~~~~~~~~~~~~~~~~~~~~~~~~~~~~~~~~~~~~~~~~~~~~<1
   1<~~~~~~~~~~~~~~~~~~~~~~~~~~~~~~~~~~~~~~~~~~~~~~~~~~~~~~~~~~~~~~~~~~~~~~~~~~~~~~~~~~~~~~~~~~~~~~~~~~<1
   8<----------------------------------------------------------------------------------------------------<8
   7<----------------------------------------------------------------------------------------------------<7
   1<~~~~~~~~~~~~~~~~~~~~~~~~~~~~~~~~~~~~~~~~~~~~~~~~~~~~~~~~~~~~~~~~~~~~~~~~~~~~~~~~~~~~~~~~~~~~~~~~~~<1

          *         *         *         *         *         *         *         *         *         *
       T  M  G  D  T  L  H  K  E  T  V  P  S  Y  E  V  L  D  N  Q  S  Y  H  I  R  R  G  L  Q  E  Q  G  A
     CtgtCATgccATCcgtAAGatgCTTttcTGTgacTGGtgaGTActcAACcaaGTCattCTGagaATAgtgTATgcgGCGaccGAGttgCTCttgCCCggc
9505>ctgtcatgccatccgtaagatgcttttctgtgactggtgagtactcaaccaagtcattctgagaatagtgtatgcggcgaccgagttgctcttgcccggc>9604
   1<~~~~~~~~~~~~~~~~~~~~~~~~~~~~~~~~~~~~~~~~~~~~~~~~~~~~~~~~~~~~~~~~~~~~~~~~~~~~~~~~~~~~~~~~~~~~~~~~~~<1
   1<~~~~~~~~~~~~~~~~~~~~~~~~~~~~~~~~~~~~~~~~~~~~~~~~~~~~~~~~~~~~~~~~~~~~~~~~~~~~~~~~~~~~~~~~~~~~~~~~~~<1
   1<~~~~~~~~~~~~~~~~~~~~~~~~~~~~~~~~~~~~~~~~~~~~~~~~~~~~~~~~~~~~~~~~~~~~~~~~~~~~~~~~~~~~~~~~~~~~~~~~~~<1
   8<----------------------------------------------------------------------------------------------------<8
   7<----------------------------------------------------------------------------------------------------<7
   1<~~~~~~~~~~~~~~~~~~~~~~~~~~~~~~~~~~~~~~~~~~~~~~~~~~~~~~~~~~~~~~~~~~~~~~~~~~~~~~~~~~~~~~~~~~~~~~~~~~<1

          *         *         *         *         *         *         *         *         *         *
      D  I  R  S  L  V  A  G  C  L  L  V  K  F  T  S  M  M  P  F  R  E  E  P  R  F  S  E  L  I  K  G  S
     GTCaatACGggaTAAtacCGCgccACAtagCAGaacTTTaaaAGTgctCATcatTGGaaaACGttcTTCgggGCGaaaACTctcAAGgatCTTaccGCTg
9605>gtcaatacgggataataccgcgccacatagcagaactttaaaagtgctcatcattggaaaacgttcttcggggcgaaaactctcaaggatcttaccgctg>9704
   1<~~~~~~~~~~~~~~~~~~~~~~~~~~~~~~~~~~~~~~~~~~~~~~~~~~~~~~~~~~~~~~~~~~~~~~~~~~~~~~~~~~~~~~~~~~~~~~~~~~<1
   1<~~~~~~~~~~~~~~~~~~~~~~~~~~~~~~~~~~~~~~~~~~~~~~~~~~~~~~~~~~~~~~~~~~~~~~~~~~~~~~~~~~~~~~~~~~~~~~~~~~<1
   1<~~~~~~~~~~~~~~~~~~~~~~~~~~~~~~~~~~~~~~~~~~~~~~~~~~~~~~~~~~~~~~~~~~~~~~~~~~~~~~~~~~~~~~~~~~~~~~~~~~<1
   8<----------------------------------------------------------------------------------------------------<8
   7<----------------------------------------------------------------------------------------------------<7
   1<~~~~~~~~~~~~~~~~~~~~~~~~~~~~~~~~~~~~~~~~~~~~~~~~~~~~~~~~~~~~~~~~~~~~~~~~~~~~~~~~~~~~~~~~~~~~~~~~~~<1

          *         *         *         *         *         *         *         *         *         *
     N  L  D  L  E  I  Y  G  V  R  A  G  L  Q  D  E  A  D  K  V  K  V  L  T  E  P  H  A  F  V  P  L  C  F
     ttGAGatcCAGttcGATgtaACCcacTCGtgcACCcaaCTGatcTTCagcATCtttTACtttCACcagCGTttcTGGgtgAGCaaaAACaggAAGgcaAA
9705>ttgagatccagttcgatgtaacccactcgtgcacccaactgatcttcagcatctttactttcaccagcgtttctgggtgagcaaaaacaggaaggcaaa>9804
   1<~~~~~~~~~~~~~~~~~~~~~~~~~~~~~~~~~~~~~~~~~~~~~~~~~~~~~~~~~~~~~~~~~~~~~~~~~~~~~~~~~~~~~~~~~~~~~~~~~~<1
   1<~~~~~~~~~~~~~~~~~~~~~~~~~~~~~~~~~~~~~~~~~~~~~~~~~~~~~~~~~~~~~~~~~~~~~~~~~~~~~~~~~~~~~~~~~~~~~~~~~~<1
```

```
    1<~~~~~~~~~~~~~~~~~~~~~~~~~~~~~~~~~~~~~~~~~~~~~~~~~~~~~~~~~~~~~~~~~~~~~~~~~~~~~~~~~~~~~~~~~~~~~~~~~~~<1
    8<----------------------------------------------------------------------------------------------------<8
    7<----------------------------------------------------------------------------------------------------<7
    1<~~~~~~~~~~~~~~~~~~~~~~~~~~~~~~~~~~~~~~~~~~~~~~~~~~~~~~~~~~~~~~~~~~~~~~~~~~~~~~~~~~~~~~~~~~~~~~~~~~~<1


           *         *         *         *         *         *         *         *         *         *
        A  A  F  F  P  I  L  A  V  R  F  H  Q  I  S  M
      AtgcCGCaaaAAAgggAATaagGGCgacACGgaaATGttgAATactCAT
 9805>atgccgcaaaaaagggaataagggcgacacggaaatgttgaatactcatactcttcctttttcaatattattgaagcatttatcagggttattgtctcat>9904
    1<~~~~~~~~~~~~~~~~~~~~~~~~~~~~~~~~~~~~~~~~~~~~~~~~~~~~~~~~~~~~~~~~~~~~~~~~~~~~~~~~~~~~~~~~~~~~~~~~<1
    1<~~~~~~~~~~~~~~~~~~~~~~~~~~~~~~~~~~~~~~~~~~~~~~~~~~~~~~~~~~~~~~~~~~~~~~~~~~~~~~~~~~~~~~~~~~~~~~~~<1
    1<~~~~~~~~~~~~~~~~~~~~~~~~~~~~~~~~~~~~~~~~~~~~~~~~~~~~~~~~~~~~~~~~~~~~~~~~~~~~~~~~~~~~~~~~~~~~~~~~<1
    8<----------------------------------------------------------------------------------------------------<8
    7<----------------------------------------------------------------------------------------------------<7
    1<~~~~~~~~~~~~~~~~~~~~~~~~~~~~~~~~~~~~~~~~~~~~~~~~~~~~~~~~~~~~~~~~~~~~~~~~~~~~~~~~~~~~~~~~~~~~~~~~<1


           *         *         *         *         *         *         *         *         *         *
 9905>gagcggatacatatttgaatgtatttagaaaaataaacaaataggggttccgcgcacatttccccgaaaagtgccacctgacgtcttattatcatgacat>10004
    1<~~~~~~~~~~~~~~~~~~~~~~~~~~~~~~~~~~~~~~~~~~~~~~~~~~~~~~~~~~~~~~~~~~~~~~~~~~~~~~~~~~~~~~~~~~~~~~~~<1
    1<~~~~~~~~~~~~~~~~~~~~~~~~~~~~~~~~~~~~~~~~~~~~~~~~~~~~~~~~~~~~~~~~~~~~~~~~~~~~~~~~~~~~~~~~~~~~~~~~<1
    1<~~~~~~~~~~~~~~~~~~~~~~~~~~~~~~~~~~~~~~~~~~~~~~~~~~~~~~~~~~~~~~~~~~~~~~~~~~~~~~~~~~~~~~~~~~~~~~~~<1
    7<-------------------------------------C----AGGGGT~~~~~~~~~~~~~~~~~~~~~~~~~~~~~~~~~~~~~~~~~~~~~~~~~~~~<1
    7<----------------------------------------------------------------------------------------------------<7
    1<~~~~~~~~~~~~~~~~~~~~~~~~~~~~~~~~~~~~~~~~~~~~~~~~~~~~~~~~~~~~~~~~~~~~~~~~~~~~~~~~~~~~~~~~~~~~~~~~<1


           *         *         *         *
10005>taacctataaaaataggcgtatcacgaggccctttcgtc>10043
    1<~~~~~~~~~~~~~~~~~~~~~~~~~~~~~~~~~~~~~~~<1
    1<~~~~~~~~~~~~~~~~~~~~~~~~~~~~~~~~~~~~~~~<1
    1<~~~~~~~~~~~~~~~~~~~~~~~~~~~~~~~~~~~~~~~<1
    1<~~~~~~~~~~~~~~~~~~~~~~~~~~~~~~~~~~~~~~~<1
    6<------------------GTAT---------------TC<1
    1<~~~~~~~~~~~~~~~~~~~~~~~~~~~~~~~~~~~~~~~<1
```
